# Integrated stress response associated with dark microglia promotes microglial lipogenesis and contributes to neurodegeneration

**DOI:** 10.1101/2024.03.04.582965

**Authors:** Anna Flury, Leen Aljayousi, Siaresh Aziz, Hye-Jin Park, Mohammadparsa Khakpour, Colby Sandberg, Fernando González Ibáñez, Olivia Braniff, Pragney Deme, Jackson D. McGrath, Thi Ngo, Jack Mechler, Denice Moran Ramirez, Dvir Avnon-Klein, John W. Murray, Jia Liu, Norman J. Haughey, Sebastian Werneburg, Marie-Ève Tremblay, Pinar Ayata

## Abstract

Microglia, the brain’s primary resident immune cells, are a heterogeneous population and can assume phenotypes with diverse functional outcomes on brain homeostasis. In Alzheimer’s disease (AD), where microglia are a leading causal cell type, microglia subsets with protective functions have been well characterized. Yet, the identity of microglia subsets that drive neurodegeneration remains unresolved. Here, we identify a neurodegenerative microglia phenotype that is characterized by a conserved stress signaling pathway, the integrated stress response (ISR). Using mouse models to activate or inhibit ISR in microglia, we show that ISR underlies the ultrastructurally distinct “dark” microglia subset linked to pathological synapse loss. Inducing microglial ISR in murine AD models exacerbates neurodegenerative pathologies, such as Tau pathology and synaptic terminal loss. Conversely, inhibiting microglial ISR in AD models ameliorates these pathologies. Mechanistically, we present evidence that ISR promotes the secretion of toxic long-chain lipids that impair neuron and oligodendrocyte homeostasis in vitro. Accordingly, small molecule-based inhibition of lipid synthesis in AD models ameliorates synaptic terminal loss. Our results demonstrate that activation of ISR within microglia represents a novel pathway contributing to neurodegeneration and suggest that this may be sustained, at least in part, by the secretion of long-chain lipids from ISR-activated microglia.

## Main

Alzheimer’s disease (AD) risk factors gleaned by genome-wide association studies largely map to microglia—the brain’s innate immune cells, thus strongly implicating these cells in the etiology of the disease^1,2^. Microglia co-exist in diverse functional states with distinct molecular signatures and functional outcomes on brain homeostasis^3,4^. A well-characterized subset of disease-associated microglia (DAM) is found at sites of neurodegeneration in AD mouse models, restricting neuropathologies and thereby protecting the brain^5–7^. Other microglia subsets likely play an active role in the neurodegenerative process by excessively pruning synapses^8–10^, exacerbating Tau pathology^11–13^, or impairing glial support^14–16^, yet their identity remains elusive.

Ultrastructural studies in humans and mice have attributed neurodegenerative properties to “dark microglia,” a subset that exhibits markers of cellular stress^17–19^. These microglia appear in greater numbers in disease conditions and frequently interact with synaptic structures, connoting a neurodegenerative faculty. They are further characterized by dilated endoplasmic reticulum and Golgi, an electron-dense, condensed, dark cyto- and nucleoplasm, and altered mitochondria—collectively, signs of cellular stress^17–19^. Cells (from prokaryotes to mammalian cells) can cope with stress by activating various stress response pathways. Among these pathways, the integrated stress response (ISR) has been heavily implicated in AD pathology^20,21^, where the level of ISR strongly correlates with patients’ cognitive decline^22,23^. ISR is also implicated in genetic AD risk. The strongest genetic risk factor for sporadic AD, the *APOE4* allele^24^, induces ISR in microglia^25^ and leads to the expansion of a microglia subset characterized by cellular stress^26^. Moreover, *EIF2B,* a direct ISR target, has been recently identified as an AD risk gene^27^. Together, these studies implicate microglial ISR and, more generally, microglial stress in AD causality and pathology.

In this study, we show that ISR is activated in dark microglia and that intrinsic activation of ISR leads microglia to autonomously acquire early features of a dark microglia phenotype. Using mouse models to turn on and off ISR in microglia, we show that ISR upregulates genes associated with diverse cellular stress responses and metabolic pathways that are uncoupled from inflammatory networks. In AD models, microglial ISR significantly worsens Tau accumulation and presynaptic protein loss, while its inhibition blunts the development of these pathological features. ISR promotes lipid synthesis pathways in vivo and microglial secretion of long-chain lipids in vitro. The microglia-secreted lipids impair neuron and oligodendrocyte progenitor cell (OPC) homeostasis and survival. Small molecule inhibition of fatty acid synthesis in AD models prevents neurodegenerative synaptic protein loss. Together, these results identify ISR-activated microglia as a neurodegenerative phenotype and reveal lipid secretion as a novel mechanism of microglia-mediated neurotoxicity.

### ISR is activated in dark microglia

Dark microglia—a microglia subset whose frequent interactions with synapses hint at the potential role of stressed microglia in excessive synaptic pruning—are rarely detected in healthy mice but frequently in the APP/PS1 mouse model of amyloid pathology (∼43% of plaque-associated microglia)^17,18^ and aged individuals^18,19^. Yet, their presence in AD patients has not been shown. To address this, we performed ultrastructural analysis by scanning electron microscopy on the postmortem brain of a 79-year-old female AD patient. We found dark microglia characterized by dark cytoplasm and nucleus, fragmented and dilated ER, aberrant mitochondria, and a loss of chromatin pattern induced in this patient’s brain (Fig. 1a, Supplementary Fig. 1).

**Figure 1.**
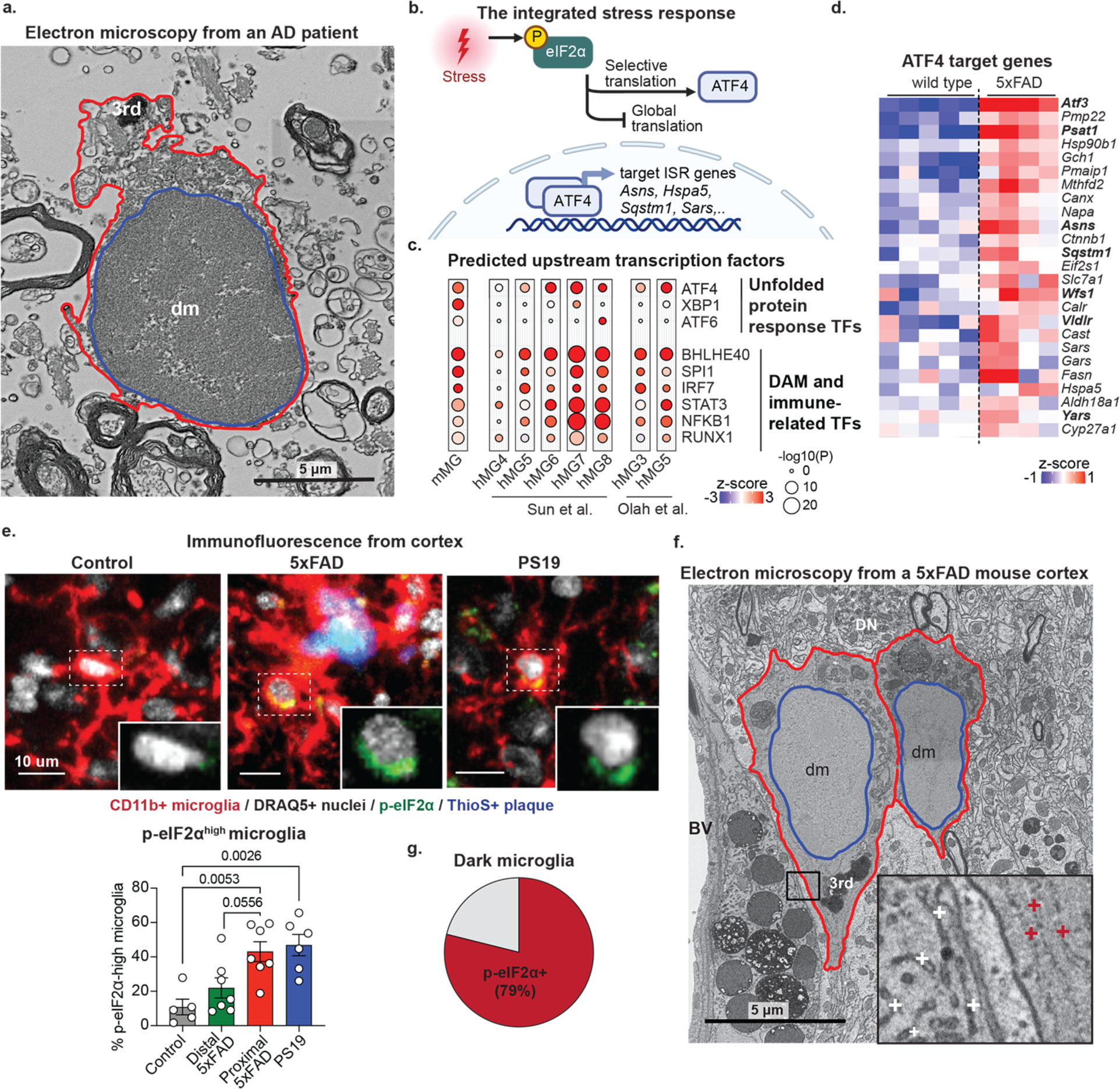
ISR is activated in the microglia of AD mouse models and dark microglia. **(a)** Representative electron microscopy image shows dark microglia (dm) containing tertiary lysosomes (3rd) in the hippocampus of a 79-year-old female patient with AD. **(b)** Schematic shows the main events in integrated stress response (ISR). Adapted from “Stress Response Mechanism in Cells” by BioRender.com (2023). **(c)** The balloon plot represents the p-value (size) and z-score (color) for Ingenuity Pathway Analysis (IPA) upstream regulator enrichment for differentially regulated genes (p value<0.05, baseMean>20 by DESeq2) in translating ribosome affinity purification (TRAP) sequencing from 7-month-old control (n=5) and 5xFAD (n=4) mouse microglia (mMG) and for human microglia subsets (hMG) identified by Sun et al.^36^ and Olah et al^35^. **(d)** Heatmap shows gene z-scored variance stabilizing transformation (vst, gene expression score) of ATF4 target ISR genes that are differentially regulated genes, identified as in (c). Canonical ATF4 targets^33^ are shown in bold. **(e)** Representative immunofluorescence images (top) and quantification (bottom) of p-eIF2α levels in microglia. p-eIF2α: green, CD11b+ microglia: red, ThioS+ dense-core plaques: blue, DRAQ5+ nuclei: gray. The bar graph shows the percentage of p-eIF2α^high^ microglia (above the dotted line in the dot plot in Supplementary Fig. 2f) in the cortex of 6-month-old control (n=5), 6-month-old 5xFAD (n=7), and 8-month-old control PS19 mice (n=6). In 5xFAD mice, p-eIF2α intensity in microglia proximal to plaques (microglial surfaces less than 5 μm away from a plaque) and distal microglia (microglial surfaces more than 5 μm away from a plaque) are shown separately—one-way ANOVA with multiple comparisons. Bar graph with individual data points shows mean ± s.e.m. **(f)** Representative electron microscopy image shows two adjacent dark microglia (dm) contacting a dystrophic neurite (DN) located near a blood vessel (BV) in a 5xFAD mouse. Red outline: cellular membrane. Blue outline: nuclear membrane. The inset shows a zoomed-in view illustrating examples of p-eIF2α immunostaining as dot patterns decorating the ER cisternae in dark microglia (red plus) and p-eIF2α-negative ER in an adjacent pericyte (white plus). 3rd: tertiary lysosome (scale bars = 5 μm). **(g)** The pie chart shows the percentage of p-eIF2α+ dark microglia in red (n=53 cells from a female 5xFAD mouse).

Due to the strong association of ISR in AD pathology and causality, we postulated that ISR may be activated in dark microglia. ISR is induced tissue-wide in postmortem brains of AD patients and AD mouse models^28–31^, yet its induction specifically in microglia has not been documented. ISR is initiated when one of the four stress sensor kinases (PERK, PKR, GCN2, and HRI) is activated and phosphorylates translation initiation factor eIF2α (Fig. 1b, Supplementary Fig. 2a). This event results in the attenuation of translation via EIF2B sequestration, thereby alleviating the burden of protein synthesis. Simultaneously, it induces the selective translation of transcripts containing an upstream open reading frame, most notably transcription factor ATF4^32^. ATF4—the main effector of ISR—induces the transcription of specific ISR genes involved in cellular adaptation by cooperating with one of its dimerization partners^32,33^. The magnitude of eIF2α phosphorylation and ATF4 binding partners determine the outcome of ISR.

We first checked whether ATF4 target networks are enriched in AD patients, specifically in microglia. Single-cell sequencing studies of human AD brains have consistently reported microglial subsets characterized by cellular stress and altered metabolism.^34–36^ To probe these datasets for ATF4 network enrichment, we used the upstream regulator analysis tool of Ingenuity Pathway Analysis^37^. We found that many disease-associated human microglia subsets from two recent studies^35,36^ showed a selective enrichment of the ATF4 target gene network (Fig. 1c, Supplementary Fig. 2b). The enrichment of other ER stress and unfolded protein response (UPR) transcription factor target networks, including ATF6 and XBP1, was relatively lower.

We next asked whether ISR genes are induced in microglia in AD mouse models. Although the field lacks an AD model that fully recapitulates the pathological spectrum observed in humans, we chose the commonly used 5xFAD model to test if AD-associated amyloid β (Aβ) accumulation can induce the ISR signature in microglia. This model presents amyloid pathology starting at two months in the subiculum but lacks Tau pathology^38,39^. To obtain microglial gene expression signatures from these mice, we performed cortical microglia-specific ribosomal profiling by translating ribosome affinity purification (TRAP) at 6 months, when 5xFAD mice have significant amyloid pathology in the cortex^40–43^. This analysis revealed that 5xFAD cortical microglia upregulated the expression of canonical ATF4 target genes, such as *Asns, Sqstm1, Wfs1, Atf3,* and *Vldlr* (Fig. 1c, d, Supplementary Fig. 2c-e). Together, these results revealed that ISR gene expression is induced in the microglia of AD patients and the 5xFAD mouse model.

Phosphorylation of eIF2α is the core event in the ISR pathway and serves as a robust proxy for ISR activation^20^ (Fig. 1b, Supplementary Fig. 2a). To solidify our finding that ISR is activated in microglia in AD, we assessed phosphorylated eIF2α (p-eIF2α) levels in microglia. Using immunofluorescence, we found that microglia in 6-month-old 5xFAD brains exhibited higher levels of p-eIF2α, specifically in microglia proximal to plaques (microglia that are closer than 5 μm to plaques, Supplementary Fig. 2f). p-eIF2α^high^ microglia (as defined in Supplementary Fig. 2f) comprised ∼40% of proximal microglia (Fig. 1e), similar to dark microglia in the APP/PS1 model^19^. This finding raised the question of whether ISR induction in microglia was limited to Aβ accumulation or whether other disease-associated protein aggregates could induce the same pathway in microglia. To address this, we quantified p-eIF2α^high^ microglia in the PS19 Tau pathology model^44^. In these mice, neurofibrillary Tau tangles are detectable in the cortex at ∼6 months, with significant neuronal loss and brain atrophy by 8 months^44^. In 8-month-old PS19 Tau mice, p-eIF2α^high^ microglia were present at ∼40%—similar levels to microglia proximal to Aβ plaques (Fig. 1e, Supplementary Fig. 2f).

To test whether ISR is activated in dark microglia, we performed immunostaining for p-eIF2α and ultrastructural analysis by electron microscopy. We found that dark microglia in the 5xFAD mice contained strong punctate p-eIF2α staining localized on their ER membrane (in 79% of dark microglia imaged (n=76 cells), Fig. 1f, g, Supplementary Fig. 3). Our results revealed that ISR is induced in the microglia of AD patients and AD mouse models, and specifically within the neurodegeneration-associated dark microglia.

### Generation and validation of microglia-specific ISR models

We reasoned that ISR induction within microglia without confounding factors might provide critical insight into their functional outcome on the brain. To regulate microglial ISR, we generated two opposing *Cx3cr1^CreErt2/+^*-driven mouse models (which target microglia with ∼99% efficiency)^45–47^:

1. The chemogenetic ISR-ON model phosphorylates eIF2α via a drug-inducible eIF2α kinase PKR (iPKR)^48,49^ and temporarily turns on ISR (Fig. 2a). This is achieved by co-expressing the C-terminal kinase domain of eIF2α kinase PKR with an engineered target site for NS3 (from the Hepatitis C virus) together with this protease^48,49^. The kinase is typically degraded by NS3, but upon inhibition of NS3 by the Hepatitis C drug asunaprevir (ASV), PKR is free to phosphorylate eIF2α (Fig. 2a). eIF2α kinases sense different cellular stresses due to their unique regulatory domains, and they all lead to eIF2α phosphorylation due to their structurally similar kinase domains, for which the only validated substrate is eIF2α^20,32,50^. The advantage of the iPKR model is that it uses the C-terminal kinase domain of PKR to selectively induce eIF2α phosphorylation. Thus, off-target effects of PKR’s regulatory N-terminal domain are omitted. Moreover, with this design, endogenous PKR is not obstructed^48,49^.
2. Serving the opposite role is an ISR-OFF model^51,52^, where eIF2α phosphorylation, and therefore ISR, is prevented via a phosphomutant (Fig. 2a). The *Eif2^A^* phosphomutant model, which is homozygous for a Serine to Alanine mutation at the primary phosphorylation site (Ser 51) of eIF2α, simultaneously carries a Cre-excisable wild-type *Eif2s1* transgene (encoding eIF2α, Fig. 2a). Cre expression removes the wild-type copy of *Eif2s1* in microglia, leaving only the phosphomutant, while other cells express both.

**Figure 2.**
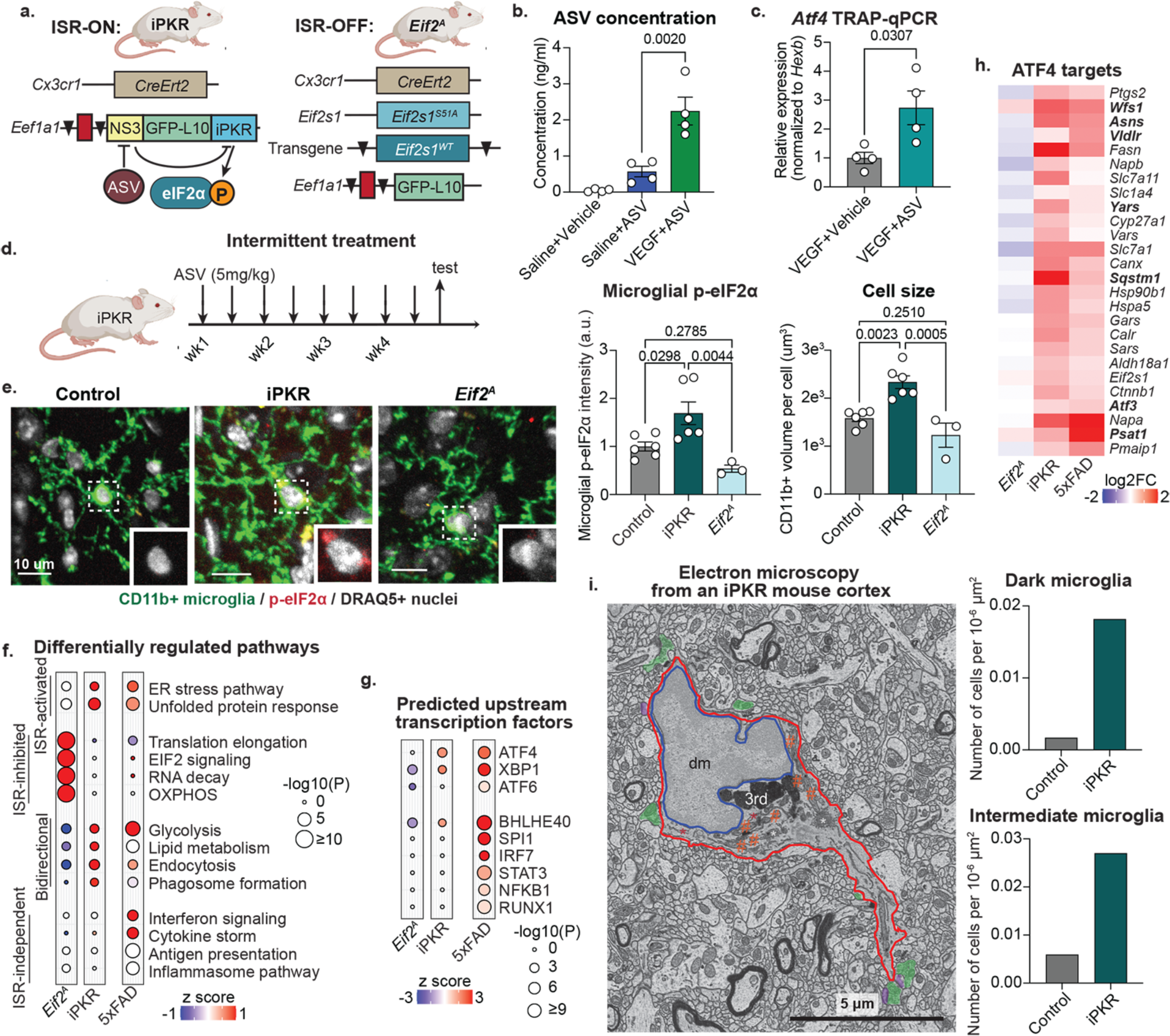
Microglial ISR impacts gene expression profiles and generates dark microglia. **(a)** The schematic shows the constructs for the iPKR (ISR-ON) and *Eif2^A^* (ISR-OFF) mice. **(b)** The bar graph shows mass spectrometry-based quantification of ASV in wild-type mice (n=4 mice/group) 2 hours after oral treatment with ASV —one-way ANOVA with multiple comparisons. **(c)** The bar graph shows the qPCR-based measurement of *Atf4* in microglia-specific TRAP experiments from animals exposed to VEGF followed by a single oral vehicle administration and VEGF followed by a single oral ASV administration (n=4/group)—unpaired two-tailed t-test. Vehicle or ASV administration was performed 30 minutes after VEGF. Tissue was collected 2.5 hours after VEGF administration. **(d)** The schematic shows the bi-weekly month-long treatment of iPKR mice in the intermittent treatment paradigm. **(e)** Representative immunofluorescence images (left) and quantifications (right). p-eIF2α: red, CD11b+ microglia: green, DRAQ5+ nuclei: gray. The bar graphs show the mean p-eIF2α intensity in microglial cytosol and the microglial cell volume (cell size) in the cortex of 6-month-old ASV-treated control (n=6), ASV-treated iPKR (n=6), and *Eif2^A^* mice (n=3)—one-way ANOVA with multiple comparisons. Bar graphs with individual data points show mean ± s.e.m. **(f, g)** The balloon plots represent the p-value (size) and z-score (color) for Ingenuity Pathway Analysis (IPA) pathways (f) and upstream regulators (g) of differentially regulated genes identified by DESeq2 (p value<0.05, baseMean>20 by DESeq2) in microglia TRAP sequencing from 6-month-old mice. Microglia-specific TRAP mice (n=5) were used as controls for the *Eif2^A^* (n=4) and 5xFAD mice (n=4). Vehicle-treated iPKR mice (VEGF followed by DMSO:PEG, n=4) were used as controls for ASV-treated iPKR mice (n=5). **(h)** Heatmap shows log2 (fold change) of ATF4 target genes by DESeq2. **(i)** Representative electron microscopy image (left) and quantification (right) from an iPKR mouse. A dark microglia (dm) is adjacent to axon terminals (green) and dendritic spines (purple) in an iPKR mouse, containing several tertiary lysosomes (3rd). Red outline: cellular membrane. Blue outline: nuclear membrane. Orange hash: mitochondria. White asterisk: Golgi apparatus. Red asterisk: dilated ER. The bar graphs show the number of dark (top) and intermediate microglia (bottom) in a control and an iPKR mouse.

To induce ASV-mediated activation of iPKR, we orally administered ASV after temporarily permeabilizing the blood-brain barrier (BBB) using low-dose (1 μg/kg) VEGF injection^53,54^ (Supplementary Fig. 4a, b). 30 minutes after VEGF injection, we orally administered the clinically established dose of 5 mg/kg ASV^55,56^. Pre-clinical pharmacokinetics studies found that ASV levels increase steeply 1 hour after oral administration and peak between 2-4 hours^55,56^. Mass spectrometry analysis of ASV 2 hours after ASV administration revealed that VEGF increased BBB penetrance of ASV ∼5-fold (Fig. 2b). Leveraging the iPKR design, which contains the *Rpl10a-eGFP* gene in the construct^42^, we performed microglia-specific TRAP-qPCR analysis 2 hours after oral delivery^42^. This analysis showed that the ribosomal association of *Atf4* increased by ∼3-fold in iPKR mice (Fig. 2c) but not in wild-type mice (Supplementary Fig. 4c, d).

Aging is a strong risk factor for AD, suggesting the gradual accumulation of minor insults^57^. Notably, repeated ISR activation can result in gene expression changes that reprogram the cellular identity^58–61^. To mimic these processes, we induced a mild, intermittent ISR in microglia in the iPKR mice. We focused on relatively mild changes in ISR that are expected to reflect the neurodegenerative environment during aging rather than include blunt tools such as continuous ISR and translation arrest that could lead to programmed cell death^32^. We administered ASV biweekly for four weeks (following VEGF administration) and harvested mouse brains 2 hours after the last administration at 6 months of age (Fig. 2d). By immunofluorescence, we detected a significant increase in p-eIF2α levels in iPKR mice compared to wild-type and *Eif2^A^* mice (Fig. 2e).

### ISR regulates metabolic, engulfment, and stress response pathways and induces dark microglia

Having established these microglia-specific ISR models, we determined how activation or inhibition of ISR impacts the molecular signature of microglia. We performed microglia-specific TRAP-sequencing and detected four categories of differentially regulated pathways:

1. ISR-activated pathways, including UPR and ER stress response pathways and ATF4 and XBP1 target genes. These were induced in iPKR microglia but not in *Eif2^A^* microglia (Fig. 2f-h, Supplementary Fig. 4e-g). Notably, chemogenetic induction of PKR could sufficiently induce other stress pathways, such as those downstream XBP1, validating crosstalk among these pathways (Supplementary Fig. 4h). Furthermore, the induction of these pathways was comparable in 5xFAD microglia, showing the relevance of the iPKR model to AD.
2. ISR-inhibited pathways, including EIF2 signaling, translation elongation, RNA decay, and oxidative phosphorylation (OXPHOS). These were upregulated in *Eif2^A^* microglia and were unchanged in iPKR and 5xFAD microglia.
3. Bidirectionally regulated pathways, including glycolysis, lipid metabolism, endocytosis, and phagocytosis. These were upregulated in the iPKR and 5xFAD microglia and reduced in *Eif2^A^* microglia.
4. ISR-independent pathways, including antigen representation, cytokine storm, interferon signaling, and inflammasome pathways, as well as SPI1, IRF7, STAT3, NFKB1, and RUNX1 target networks. These immune-related pathways were strongly upregulated in 5xFAD microglia but remained unaffected in iPKR or *Eif2^A^* microglia (Fig. 2f, g). One exception to these target networks was the BHLHE40 target network, which was bidirectionally regulated at moderate levels. When interrogating the affected BHLHE40 target genes, we found that these included largely glycolytic and fatty acid metabolism genes, including *Hk1, Hk2, Pfkp, Pla2g4a,* and *St3gal5*. Collectively, the gene expression signatures impacted by ISR involved protein homeostasis (proteostasis) and metabolic genes without affecting inflammatory pathways.

Morphologically, we found that the intermittent ISR led to a significant increase in cell size with no difference in cell number (Fig. 2e, Supplementary Fig. 5a), thus prompting us to perform an ultrastructural analysis of microglia. This analysis revealed that ISR induction by intermittent ASV treatment of iPKR mice led to a ∼10-fold increase in the density of intermediate and dark microglia compared to the ASV-treated control *Cx3cr1^CreErt2/+^* mice (Fig. 2i, Supplementary Fig. 5b, c). Of note, we found more intermediate microglia than dark microglia in the iPKR mice. Intermediate microglia harbor canonical dark microglia features of aberrant organelles, namely ER dilation and altered mitochondria.

Furthermore, they exhibit a less-defined heterochromatin pattern and dark cytoplasm. However, they do not fully recapitulate the latter two phenotypes to the degree of canonical dark microglia^19^. This finding validated that ISR alone is sufficient to induce the early features of dark microglia. Still, other stress response pathways likely contribute to the full establishment of the dark microglia phenotype.

### ISR leads to the expansion of stressed microglia subsets in amyloidosis

To determine the impact of ISR on the 5xFAD background, we crossed the microglia-specific iPKR and *Eif2^A^* mice to 5xFAD mice, generating 5xFADiPKR and 5xFAD*Eif2^A^* mice, respectively. In 5xFADiPKR mice, which were treated with ASV biweekly for four weeks, microglial p-eIF2α levels did not increase compared to 5xFAD mice (Supplementary Fig. 6a). Yet, the expression of ISR-activated pathways (UPR, ER stress response, as well as ATF4, XBP1, and ATF6 target genes) and bidirectionally regulated pathways at baseline (glycolysis, lipid metabolism, endocytosis, and phagocytosis) were upregulated (Fig. 3a-c, Supplementary Fig. 6b-d, 7). ISR-independent immune pathways remained comparable to 5xFAD microglia.

**Figure 3.**
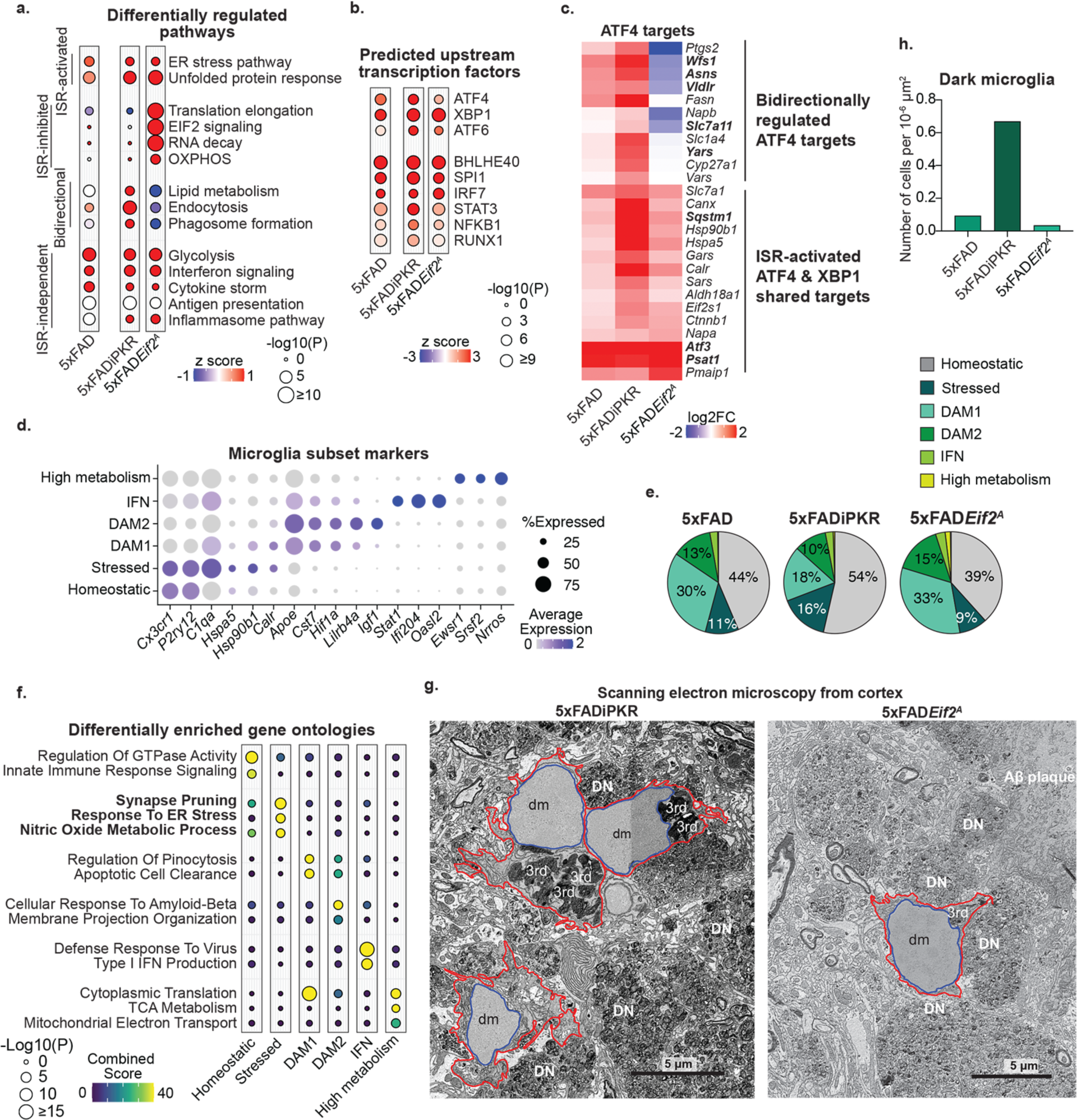
Microglial ISR impacts gene expression profiles and dark microglia phenotype in the 5xFAD model. **(a, b)** The balloon plots represent the p-value (size) and z-score (color) for Ingenuity Pathway Analysis (IPA) pathways (a) and upstream regulators (b) of differentially regulated genes identified by DESeq2 (p value<0.05, baseMean>20 by DESeq2) in microglia TRAP sequencing from 6-month-old mice. Microglia-specific TRAP mice (n=5) were used as controls for the 5xFAD*Eif2^A^* (n=3) and 5xFAD mice (n=4). Vehicle-treated iPKR mice (VEGF followed by DMSO:PEG, n=4) were used as controls for ASV-treated 5xFADiPKR mice (n=4). **(c)** Heatmap shows log2 (fold change) of ATF4 target genes that are differentially regulated (p value<0.05, baseMean>20 by DESeq2) by microglia-specific TRAP sequencing from the same mice. **(d)** The dot plot shows the average expression levels per cluster (color) and the percentage of cells (size) from each cluster identified by microglia-specific single nuclei sequencing from 5xFAD, 5xFADiPKR, and 5xFAD*Eif2^A^* mice (n=2 pooled mice per sample). **(e)** The pie charts show the proportions of microglial clusters in each genotype. **(f)** The balloon plot represents the p-value (size) and combined score (color) for gene ontologies of cluster markers in microglia-specific single nuclei sequencing. **(g)** Representative electron microscopy images. Left: Dark microglia (dm) in the cortex of a 6-month-old female 5xFADiPKR mouse that juxtaposes dystrophic neurites (DN). Right: Dark microglia (dm) near numerous dystrophic neurites (DN) and an amyloid beta (Aβ) plaque observed in the cortex of a 6-month-old male 5xFAD*Eif2^A^* mouse. Red outline: cellular membrane. Blue outline: nuclear membrane. 3rd: tertiary lysosome. **(h)** The bar graphs show the number of dark microglia in a 5xFAD, a 5xFADiPKR, and a 5xFAD*Eif2^A^* mouse. Bar graphs with individual data points show mean ± s.e.m.

In 5xFAD*Eif2^A^* microglia, p-eIF2α levels were significantly lower with a concomitant reduction in ATF4 target enrichment (Fig. 3b, c, Supplementary Fig. 6a). Paradoxically, unlike *Eif2^A^* microglia, ISR-activated ER stress and UPR pathways (such as those downstream XBP1 and ATF6) were upregulated in 5xFAD*Eif2^A^* microglia, suggesting that these pathways may compensate for the absence of ISR (Fig. 3a, b, Supplementary Fig. 6b-d, 7). Like in the *Eif2^A^* microglia, ISR-inhibited pathways associated with EIF2 signaling, translation elongation, RNA decay, and OXPHOS were induced. Of the bidirectionally regulated pathways, lipid metabolism, endocytosis, and phagocytosis were downregulated in 5xFAD*Eif2^A^* microglia, but glycolysis was strongly upregulated. Finally, ISR-independent immune pathway signatures remained unchanged.

Our TRAP sequencing results led us to consider how activation or repression of ISR could impact the amyloidosis-induced microglia subsets. To address this, we performed microglia-specific single nuclei sequencing and identified several well-characterized microglia subsets represented in all our mouse models (Fig. 3d-f, Supplementary Fig. 8). We found that the stressed and homeostatic microglia subsets were represented at considerably higher levels in 5xFADiPKR mice but at slightly lower levels in 5xFAD*Eif2^A^* mice. In 5xFADiPKR mice, the increase in these populations was counterbalanced by a reduction of the DAM subset, which was slightly increased in 5xFAD*Eif2^A^* mice. Notably, the reduction in DAM numbers in the 5xFADiPKR mice was primarily attributable to the TREM2-independent DAM1 population^5^, whereas the mild increase in 5xFAD*Eif2^A^* mice was equally represented in DAM1 and TREM2-dependent mature DAM2 subsets^5^. The IFN subset remained unchanged in all groups. Unexpectedly, we found that unleashing translation in 5xFAD*Eif2^A^* mice generated a small novel subset (that we coined “high metabolism subset”) characterized by genes associated with protein translation, TCA cycle, and mitochondrial metabolism (Fig. 3d-f, Supplementary Fig. 8). Together, these results reveal ISR as a regulator of microglial subset identity in amyloidosis models.

Baseline autonomous activation of ISR in microglia triggered the generation of dark microglia in the iPKR mice, concomitant with gene expression changes associated with increased cellular stress response pathways. In 5xFADiPKR mice, these pathways and the stressed microglia subset were also induced, but these were only nominally reduced in 5xFAD*Eif2^A^* mice. Therefore, we asked how activation or inhibition of ISR would impact the dark microglia phenotype in the 5xFAD model. We found many clusters of dark microglia in a 5xFAD-iPKR mouse (Fig. 3g, h, Supplementary Fig. 9). The prevalence of dark microglia in 5xFAD*Eif2^A^* mice remained comparable to 5xFAD mice (Fig. 3g, h, Supplementary Fig. 10). These observations build upon our earlier findings that ISR activation in the absence of other stimuli induces intermediate and dark microglia, clarifying that ISR is sufficient but not required to induce the dark microglia phenotype.

### Modulation of microglial ISR impacts AD-associated neurodegenerative pathologies

AD is characterized by two major pathologies—Aβ plaques and intracellular Tau aggregates^62^. Microglia, particularly DAM, play critical protective roles in restricting amyloid burden and Aβ-induced axonal dystrophy^62^. We found that ISR impaired the protective function of microglia on amyloid pathology. Total dense-core plaque numbers and volume, as well as axonal dystrophy, increased in 5xFADiPKR mice with no significant changes to plaque size or shape (Fig. 4a, Supplementary Fig. 11a). Surprisingly, these amyloid-associated pathologies were not mitigated in 5xFAD*Eif2^A^* mice. In line with our single-cell findings, we found that ISR modestly impacted microglial engagement with the plaques (assessed by plaque-associated microglia numbers and microglial plaque infiltration), with a mild loss in 5xFADiPKR mice and a mild rescue in 5xFAD*Eif2^A^* mice, without significant changes to overall microglia numbers or size (Fig. 4b, Supplementary Fig. 11b). These results imply that moderate microglial ISR is dispensable for microglia’s protective functions against amyloid pathology, but excessive ISR is detrimental.

**Figure 4.**
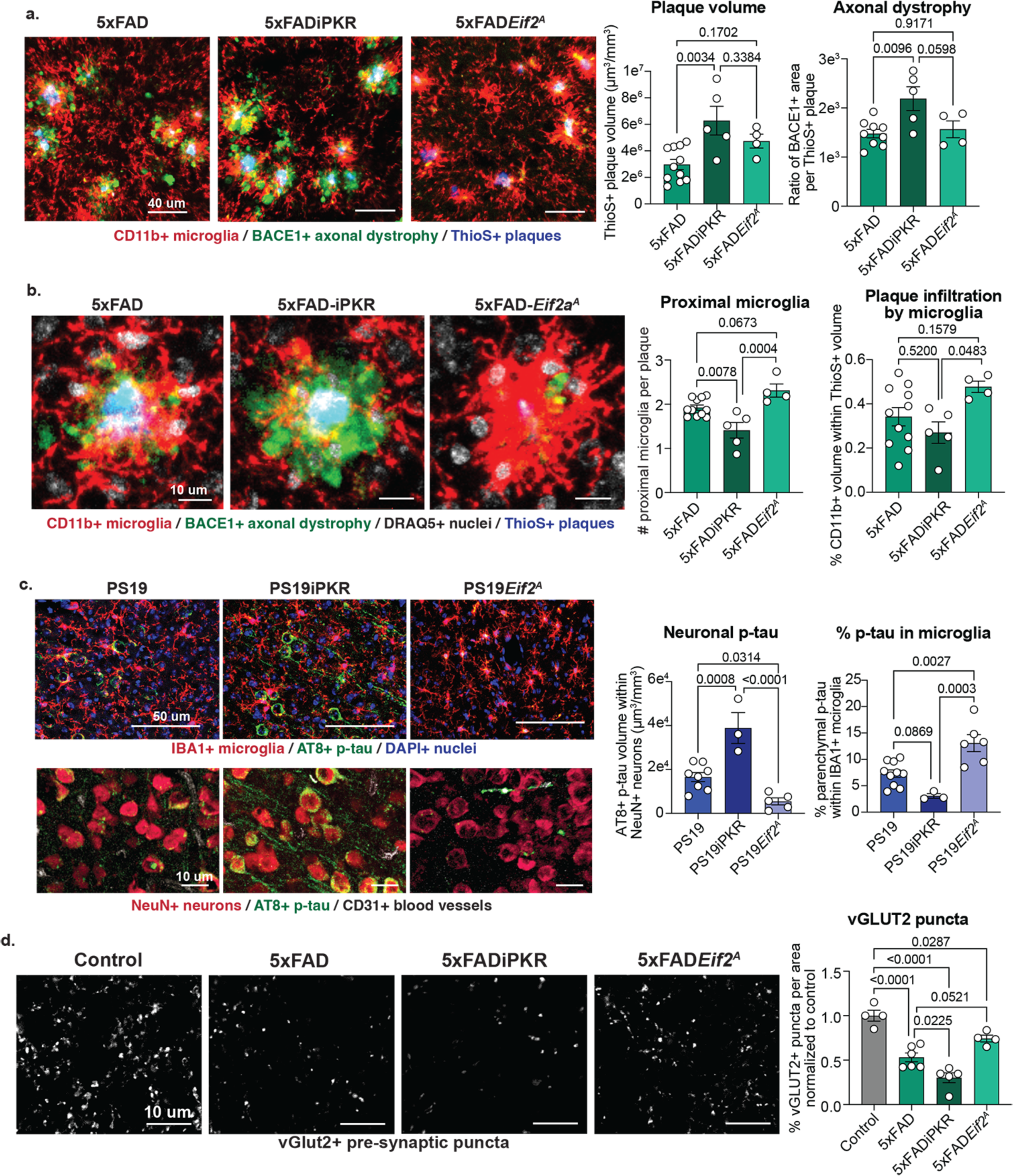
Microglial ISR has a strong impact on AD-associated pathologies. **(a)** Representative immunofluorescence images (left) and quantification (right). CD11b+ microglia: red, BACE1+ axonal dystrophies: green, ThioS+ dense-core plaques: blue. The first bar graph shows the total ThioS+ plaque volume in the cortex of 6-month-old 5xFAD (n=11), 5xFADiPKR (n=5), and 5xFAD*Eif2^A^* mice (n=4). The second shows BACE1+ axonal dystrophy volume normalized to ThioS+ plaque volume in the cortex of 6-month-old 5xFAD (n=9), 5xFADiPKR (n=5), and 5xFAD*Eif2^A^* mice (n=4). **(b)** Representative immunofluorescence images (left) and quantification (right). CD11b+ microglia: red, BACE1+ axonal dystrophies: green, DRAQ5+ nuclei: gray, ThioS+ dense-core plaques: blue. The bar graphs show the number of proximal microglia to plaques (nuclei of microglia less than 5 μm away from a plaque) per plaque and plaque infiltration by microglia calculated as CD11b+ microglial volume within ThioS+ plaques (normalized to plaque volume) in the cortex of 6-month-old 5xFAD (n=11), 5xFADiPKR (n=5), and 5xFAD*Eif2^A^* mice (n=4) quantified from (a). **(c)** Representative immunofluorescence images (left) and quantification (right) show Top: AT8+ hyperphosphorylated Tau: green, IBA1+ microglia: red, DAPI+ nuclei: blue; Bottom: AT8+ hyperphosphorylated Tau: green, NeuN+ neurons: red, CD31+ blood vessels: gray. The bar graphs show total AT8+ phospho-Tau volume within NeuN+ neurons in the cortex of 8-month-old PS19 (n=8), PS19iPKR (n=3), and PS19*Eif2^A^* mice (n=6) and percent AT8+ phospho-Tau volume within CD11b+ microglia in the cortex of 8-month-old PS19 (n=10), PS19iPKR (n=3), and PS19*Eif2^A^* mice (n=6). One outlier in the PS19*Eif2^A^* group was identified by Grubb’s test and removed. **(d)** Representative immunofluorescence images (left) and quantification (right) show presynaptic VGLUT2+ puncta: gray. The bar graph shows VGLUT2 density normalized in the layer V cortex of 6-month-old control (n=4), 5xFAD (n=6), 5xFADiPKR (n=5), and 5xFAD*Eif2^A^* mice (n=4). All analyses use one-way ANOVA with multiple comparisons. Bar graphs with individual data points show mean ± s.e.m.

In AD models, microglia are thought to restrict plaque-associated pathologies by contributing to the formation and compaction of plaques by engaging phagocytic clearance mechanisms^63,64^. The increased plaque load in 5xFADiPKR mice led us to postulate that ISR may impair microglia phagocytosis of Aβ. To investigate this, we assessed the uptake of fluorescently labeled Aβ42 oligomers by the immortalized microglia-like cell line, BV2 cells. We induced ISR by applying salubrinal (SLB), which prevents the dephosphorylation of eIF2α, thereby accumulating its phosphorylated form (Supplementary Fig. 2a, 11c-e). 24 hours after the SLB stimulation, we fed the cells oligomerized Aβ42 and recorded their uptake by live imaging over 6 hours. This analysis revealed that SLB-treated microglia phagocytosed Aβ42 slower than control microglia (Supplementary Fig. 11f). These data suggest that an impaired phagocytic response may contribute to worsened plaque pathology in 5xFADiPKR mice.

In AD, Tau pathology tightly correlates with neurodegeneration and cognitive decline and is therefore considered neurodegenerative^62^. Microglia have a complex role in Tau pathology that is poorly understood. They can be protective by taking up and breaking down pathological Tau released by neurons^65–67^; they can also contribute to Tau spread by secreting the phagocytosed Tau aggregates via exosomes^11^. To measure the impact of microglial ISR on Tau pathology, we bred our ISR models to the PS19 Tau model and generated PS19iPKR and PS19*Eif2^A^* mice. ISR induction in PS19iPKR mice by intermittent ASV treatment led to a significant increase in hyperphosphorylated Tau levels, specifically within neurons. Conversely, its occlusion in PS19*Eif2^A^* mice led to a significant decrease in hyperphosphorylated Tau levels (Fig. 4c, Supplementary Fig. 11g). Furthermore, we detected a higher proportion of Tau in the microglia of PS19*Eif2^A^* mice and a lower proportion in PS19-iPKR mice (Fig. 4c, Supplementary Fig. 11g), suggesting ISR may impact microglial uptake, degradation, or release of Tau, thus contributing to Tau pathology.

Microglia can contribute to cognitive impairment and neurodegeneration by causing synapse loss^9^, which is the strongest pathological correlate of cognitive decline^68^. To determine whether ISR plays a role in these detrimental outcomes of microglia, we quantified pre-synaptic terminal densities in layer V cortex of 6-month-old mice^69^. We found a significant reduction of pre-synaptic glutamate transporter VGLUT2 in 5xFADiPKR mice and a partial rescue in 5xFAD*Eif2^A^* mice (Fig. 4d). These results suggest that microglial ISR bidirectionally regulates the loss of specific pre-synaptic terminals in amyloidosis.

### ISR induces the dysfunction of other brain cell types via lipid secretion

An important question raised by our findings is what mediates the toxicity of microglial ISR on other cell types. As microglia can release cytotoxic factors under diverse stimuli^70,71^, we hypothesized that ISR-activated microglia could release toxic factors that exacerbate neuropathologies and contribute to microglial toxicity toward neighboring cells. Therefore, we tested the impact of ISR-activated microglia-conditioned media on OPCs, astrocytes, and neurons (Fig. 5a, Supplementary Fig. 12a-c). Based on previous reports suggesting non-cell-autonomous propagation of cellular stress pathways^72,73^, we first tested whether ISR-activated microglia could transmit ISR via secreted factors. We observed that SLB-conditioned media induced the expression of ATF4 targets in recipient naïve OPCs and neurons but not astrocytes (Fig. 5b, c, Supplementary Fig. 12d). Moreover, SLB-conditioned media increased the apoptosis rate in OPCs (as measured by %cCASP3+ cells, Fig. 5d, Supplementary Fig. 12e). As microglia play critical roles in suppressing neuronal hyperexcitability^74^, we next tested the impact of microglial ISR on neuron firing. Notably, SLB-conditioned media significantly increased neuronal firing and reduced the number of neurons in culture (Fig. 5e, Supplementary Fig. 12f). Together, these results showed that ISR activation in microglia results in the secretion of toxic factors that induce non-cell-autonomous ISR and impair homeostasis in two majorly impacted cell types in AD^75,76^, which may contribute to its neurodegenerative impact in vivo. Previous studies discovered that reactive astrocytes secrete toxic lipids, which impair neuron and OPC survival^15^, leading us to ponder the possibility of toxic lipid secretion by microglia. In line with this idea, our TRAP sequencing results revealed that ISR led to the bidirectional regulation of lipid metabolism genes, specifically fatty acid biosynthesis (*Acly, Aacs, Fasn),* fatty acid elongation and desaturation (*Hacd3, Fads1*, *Elovl4*), biosynthesis of di- and triglycerides (*Agpat4, Gpat4*), sphingolipid biosynthesis (*Cers4, Degs1*) and cholesterol biosynthesis (*Hmgcr, Hmgcs1*, Fig. 5f, g, Supplementary Fig. 6b-d). Of note, many of these genes are direct ATF4 targets^33^. To identify which lipid species may be secreted by ISR-activated microglia, we performed a lipidomic analysis on SLB-conditioned media. We found a strong upregulation of long-chain unsaturated phospholipids (including phosphatidylcholine, plasmenylcholine, phosphatidylglycerol, and phosphatidylserine), ceramides (including dihydro and hydroxy ceramides), sphingomyelins, long chain monounsaturated cholesterol esters, and di- and triglycerides (Fig. 5f, h, Supplementary Fig. 13), in agreement with our gene expression findings in vivo.

**Figure 5.**
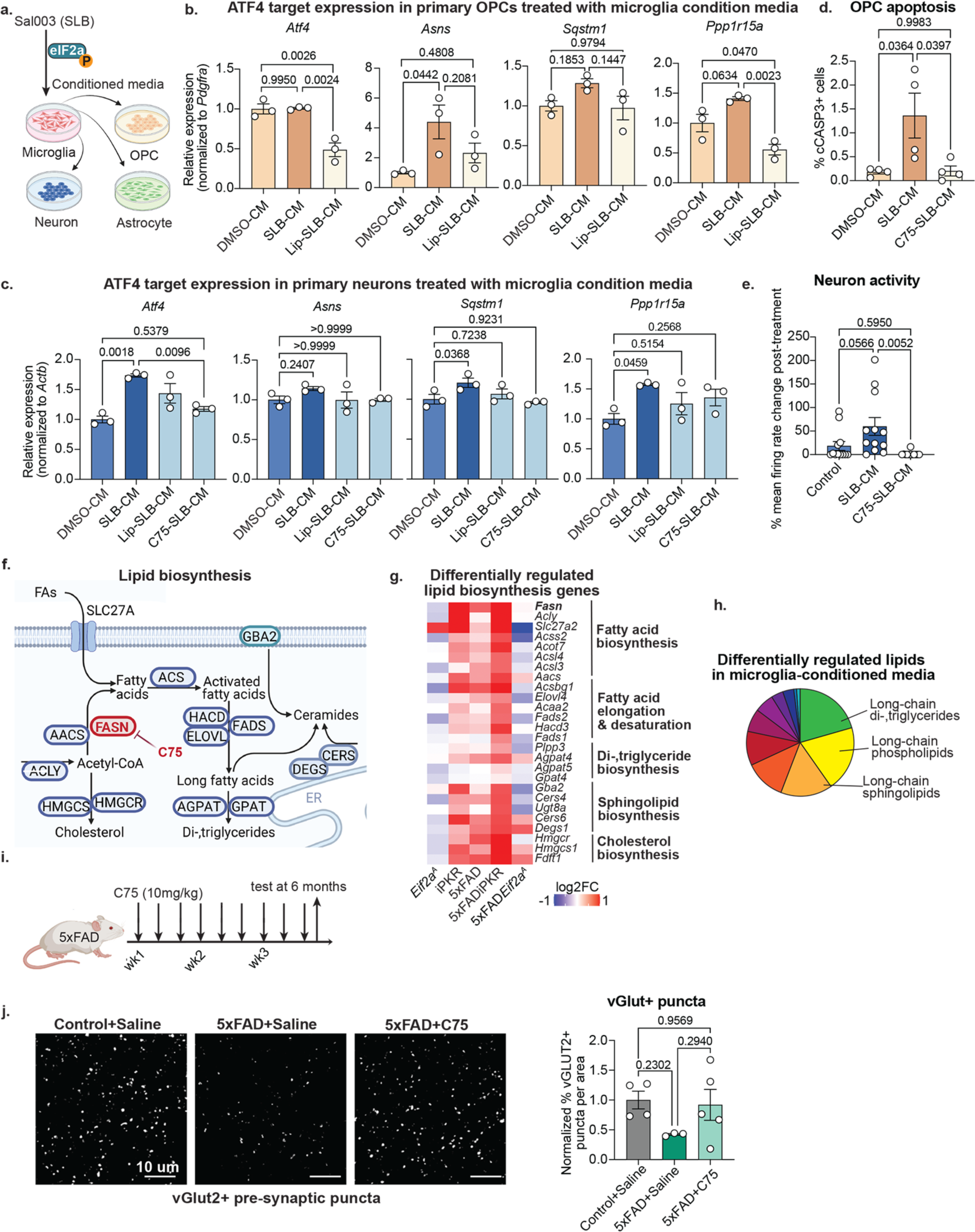
ISR-induced lipid secretion has a detrimental impact on other brain cell types. **(a)** The schematic shows the basic approach to test the predicted mechanism of microglial toxicity. Created with BioRender.com. **(b, c)** The bar graphs show the qPCR-based measurement of *Atf4* and ATF4 target genes in recipient primary OPCs (b) and primary neurons (c) 6 hours after receiving conditioned media from BV2 cells treated with DMSO (DMSO-CM), SLB (SLB-CM), SLB and C75 (C75-SLB-CM), and conditioned media from BV2 cells treated with SLB that was subjected to lipid removal (Lip-SLB-CM, n=3 wells/condition). **(d)** The bar graph shows the percentage of cCASP3+ apoptotic OLIG2+ OPCs 24 hours after receiving conditioned media from BV2 cells treated as in (b, c) (n=4 wells/condition). **(c)** Representative immunofluorescence images (left) and quantification (right) show NeuN+ neuronal cells: green, cCASP3+ apoptotic cells: red, DAPI+ nuclei: blue. The bar graphs show cCASP3+ apoptotic NeuN+ neuronal cells 24 hours after receiving conditioned media (n=4 wells/condition). **(e)** The bar shows the percentage change in the mean firing rate of primary neurons 1 hour after treatment compared to their baseline before drug treatment in an Axion multi-electrode array plate (n=12 wells/condition). **(f)** Schematic shows a simplified lipid synthesis pathway highlighting the upregulated genes in iPKR and 5xFAD microglia. Adapted from “Lipids and Proteins Involved in Lipid Uptake and Metabolism in Cardiac Lipotoxicity” by BioRender.com (2023). **(g)** Heatmap shows log2 (fold change) of lipogenesis genes that are differentially regulated (p value<0.05, baseMean>20 by DESeq2) by microglia-specific TRAP sequencing. **(h)** The pie chart shows the classification of differentially regulated lipid species in SLB-conditioned media compared to control. **(i)** Schematic shows the bi-weekly month-long treatment of iPKR mice as the intermittent treatment paradigm. **(j)** Representative immunofluorescence images (left) and quantification (right) show presynaptic VGLUT2+ puncta: gray. The bar graph shows VGLUT2 density normalized in the layer V cortex of 6-month-old vehicle-treated control (n=5), vehicle-treated 5xFAD (n=5), and C75-treated 5xFAD mice (n=5). One-way ANOVA with multiple comparisons. Bar graphs with individual data points show mean ± s.e.m.

Notably, lipid metabolism is tightly linked to protein homeostasis, and ATF4 targets include lipogenesis genes, most notably *Fasn*^77^. *Fasn,* which encodes a key rate-limiting enzyme in lipid biosynthesis, regulates palmitate synthesis from acetyl-CoA and malonyl-CoA into long-chain saturated fatty acids. *Fasn* is upregulated in 5xFAD microglia and more so in 5xFADiPKR microglia but remains low in 5xFAD-*Eif2a^A^* microglia (Fig. 5f, g). Importantly, FASN protein levels are induced in the brains of both AD mice and AD patients, and FASN-positive cells are frequently located around plaques in postmortem AD brains^78^. Therefore, we investigated whether FASN-produced fatty acids may contribute to the toxic effects of SLB-conditioned media. We inhibited FASN in BV2 cells by co-treating them with SLB and FASN inhibitor C75^79^. As an alternative approach, we removed fatty acids, lipoproteins, lipid-membrane bodies, and other lipids from the SLB-conditioned media using an adsorbent resin^80–82^. Both approaches to eliminating fatty acids in the SLB-conditioned media prevented its toxic effects on inducing ISR-associated OPC apoptosis and neuronal hyperactivity and cell loss (Fig. 5b-e, Supplementary Fig. 12e, f). Our results suggested that ISR triggers the secretion of fatty acids from microglia, which may impair the homeostasis of neurons and OPCs.

As C75 crosses the BBB^83^, we also tested whether FASN inhibition ameliorated neurodegenerative AD pathologies. To test the impact of FASN-mediated lipogenesis in vivo, we injected 5xFAD mice with 10mg/kg C75 for 3 weeks every other day and found that C75-injected 5xFAD mice had reduced loss of VGLUT2+ synaptic puncta (Fig. 5j). These results suggest that long-chain fatty acids may play an important role in neurodegeneration associated with AD.

## Discussion

Studies have consistently reported that microglia are a heterogeneous population in health and disease.^3^ However, this heterogeneity has obfuscated the connections between the molecular identity, structural phenotype, and functional outcome of microglial subsets. While much insight has been provided on DAM^5,7^ (also named MgND^6^)—a molecularly distinct subset with an associated protective function, the identity of microglia contributing to neurodegeneration is still uncertain. Here, we identify a microglia phenotype in two AD mouse models characterized by the ISR pathway. We provide evidence that ISR partly underlies the dark microglia phenotype, an ultrastructural microglial subset that has suggested neurotoxic faculties due to its frequent contact with synapses and neuronal dystrophies. In line with this, we further show that ISR-activated microglia compound several neurodegenerative pathologies characteristic of AD. Finally, we discover that the secretion of toxic lipids forms the basis for the toxicity of ISR+ microglia.

Our experiments provided direct, in vivo evidence that ISR renders microglia neurodegenerative. Cellular stress response pathways are evolutionarily conserved from prokaryotes to mammalian cells, and AD pathology likely induces the activation of many of these pathways in microglia. Our focus on ISR is based on its strong implication in AD^20,21^. ISR markers are elevated in AD^28–31^, and mechanistically, Aβ oligomers induce eIF2α phosphorylation in a PKR-dependent manner^84^. This phenomenon is not limited to AD but has also been observed in other neurodegenerative disease pathologies, including Parkinson’s and amyotrophic lateral sclerosis (ALS)^85,86^. This raises the question of whether ISR activation drives neurodegeneration or is merely a corollary of disease pathologies. AD genetics points to the former, as many AD risk variants impact EIF2 and ISR signaling pathways in microglia^87–90^, and ISR target *EIF2B* has been recently identified as an AD risk gene^27^. Notably, the strongest genetic and acquired risk factors for late-onset AD—the *APOE4* allele ^25,91^ and aging^91^, respectively—activate ISR in microglia. In contrast, exercise, a protective factor, decreases ISR^92^. Thus, a plausible scenario in AD causality could involve AD risk factors converging to trigger ISR in microglia, inducing early features of the dark microglia phenotype and thus exacerbating neurodegeneration.

Paralleling *APOE4* microglia, ISR-activated microglia in iPKR mice upregulate lipid metabolism and glycolysis-associated genes^93^ and enhance cholesterol biosynthesis^94^. The *APOE4* allele is associated with a “terminally inflammatory microglia, TIM” subset characterized by stress markers and defective Aβ phagocytosis similar to ISR-activated microglia^26^. On the pathological level, *APOE4* microglia and ISR-activated microglia both exhibit deficiencies in responding to Aβ plaques and exacerbate Tau pathology, suggesting an impaired protective response^6,25,93,95,96^. These findings thus merit further study of the interplay and potentially compounding effects of *APOE4* and ISR in microglia.

Our data provides compelling evidence that blocking microglial ISR has a substantial, although not fully restorative, impact on neurodegenerative pathologies. Previous functional studies strengthen the causal role of ISR in AD models, as genetic or pharmacological inhibition of eIF2α phosphorylation has a beneficial impact in mouse models of AD, traumatic brain injury, and other neurodegenerative diseases^84,85,97–102^. However, it is essential to note that these strategies fall short of complete ISR ablation, targeting only one of four ISR kinases or one of two eIF2α copies. A discrepancy has been observed in studies that use ISR inhibitor ISRIB in neurodegeneration models, showing conflicting results depending on the dose. When ISRIB is chronically administered at low doses, it rescues neurodegeneration in mouse models of AD and Prion disease^31,103^ and in vitro models of ALS^102^. In contrast, a single high dose of ISRIB does not have a beneficial effect^104^. In our hands, complete inhibition of microglial ISR failed to improve amyloid load and partially rescued synapse loss. While ISR was sufficient to autonomously induce other stress response pathways and dark microglia phenotype in an otherwise healthy brain, it was dispensable for the upregulation of other UPR pathway-associated genes and the generation of dark microglia in amyloidosis. This corroborates findings in liver cells, where ISR blockade triggers autonomous activation of UPR pathways^105^. These results and observations first suggest that multiple stress pathways are likely required to manifest the bona fide dark microglia phenotype, where ISR is just one of these pathways. Second, they suggest that while exorbitant ISR is detrimental, some level of ISR may be necessary to maintain cellular homeostasis. Hence, ISR may fall into a Goldilocks zone, whereby an optimum range confers beneficial outcomes, prompting future studies to target these pathways in a dose-dependent fashion.

Our findings demonstrate the ability of ISR-activated microglia to induce ISR in naïve neurons and OPCs, drawing interesting parallels with other studies describing non-cell-autonomous propagation of stress signaling. Similar to our ISR-activated microglia, tumor cells can transmit ER stress to naïve macrophages, resulting in cellular reprogramming into an immunosuppressive state that favors tumorigenesis^72^, and skeletal muscle cells can propagate UPR in a systemic fashion^73^. In the brain, inducing UPR in astrocytes impaired their synaptogenic function in vitro and induced non-cell-autonomous neurodegeneration in a model of Prion disease^106^. However, ISR induced by oxidative stress in neurons increases the resistance of naïve neurons to oxidative stress^107^. Thus, in the context of AD, the potential interplay between non-cell-autonomous stress signaling from various cell types remains to be untangled. We can speculate that this phenomenon could feed a positive feedback loop, where each stressed cell can induce cellular stress in neighboring cells and thus exacerbate the neurodegenerative process. In a broader context, our findings in microglia support a growing body of literature describing the role of non-cell-autonomous stress signaling.

Emerging evidence supports the role of lipids as extrinsic signaling molecules for non-cell-autonomous ISR, aligning with our findings in microglia. For example, stress-induced secretion of long-chain ceramides^73^ and extracellular cholesterol^108^ can propagate non-cell-autonomous ER stress in muscle cells. In the brain, reactive astrocytes can induce ISR-driven apoptosis in OPCs via the secretion of long-chain saturated fatty acids catalyzed by ELOVL enzymes in lipoprotein particles^15,109^. The large presence of cholesterol esters, triglycerides, and structural phospholipids secreted by ISR-activated microglia suggests that ISR-activated microglia, too, could secrete toxic lipids as lipoprotein particles^110^. Overall, our findings contribute to the emergence of lipids as extrinsic signaling molecules of non-cell-autonomous stress signaling. However, we cannot exclude the possibility that lipids may serve merely as transporters of other obscure toxic factors in the form of exosomes, specifically given the enrichment of structural phospholipids and sphingomyelin, which are major building blocks of exosomes and other extracellular vesicles^111^.

ISR has been previously linked to dysfunctional lipid metabolism and lipotoxicity, with a positive role of ATF4 in promoting lipogenesis^112–114^. Conversely, extracellular cholesterol and ceramides are well-characterized inducers of cellular stress and apoptotic ISR^115,116^. Our data revealed a striking bidirectional regulation of lipogenesis pathways by ISR. These data were supported by our in vitro lipidomics of microglia-secreted lipids, which revealed an increased release of cholesterol esters, ceramides, triglycerides, and structural phospholipids. Sphingolipids—particularly ceramides—are detrimentally associated with several pathological aspects of AD, including Aβ accumulation, Tau phosphorylation, ROS generation, mitochondrial dysfunction, and neurodegeneration^116,117^. Other lipid pathways, including triglycerides and cholesterol species, are dysregulated in AD^118–120^. AD genetics further implicate a disrupted cholesterol metabolism in AD casualty^121^, and elevated serum cholesterol levels have been proposed as an AD risk factor^122^. While the question of which lipid species drive ISR-induced microglial neurotoxicity warrants further study, our results showing the toxic effects of FASN-synthesized lipids align with recent preclinical findings that CMS121, a FASN inhibitor, alleviates cognitive loss^78^.

We discovered that ISR-activated microglia led to an increase in Aβ plaque load and neuronal pathological Tau levels. Our study provides several pieces of evidence that the detrimental impact of stressed microglia on amyloid pathology may be associated with defects in their engulfment and degradation: Our functional in vitro studies show reduced Aβ uptake by ISR-activated microglia. ISR strongly regulates phagocytosis pathways in vivo, and dark microglia have a characteristic accumulation of tertiary lysosomes—lysosomes that contain undigestible biomolecules indicative of defective degradation^19^. The role of microglia in Tau spread is less understood despite numerous studies. In postmortem tissues of AD patients, morphologically activated microglia were predicted to be detrimental to Tau pathology^13^, while microglial expression of DAM^5^ markers was attributed a protective role. The beneficial role of DAM was further substantiated functionally by mouse studies^123,124^. Mechanistically, microglia can uptake Tau (released by neurons or during phagocytosis of Tau-laden neurons/synapses). However, rather than fully degrading it, they secrete a portion, likely in exosomes^11,65^. Several factors, including NF-κB signaling and APOE isoforms, regulate the balance between their uptake, degradation, and secretion of Tau^67,125^. Based on the impaired engulfment and degradation and increased lipid secretion by ISR-activated microglia (potentially facilitating secretion through exosomes or other vesicles), one can speculate that ISR may also regulate this balance, shifting it toward secretion.

In neuroscience, the role of microglia in AD has been investigated mainly in the context of inflammation and DAM phenotype.^126–128^ At the same time, proteostasis and ISR are traditionally studied within neurons.^129,130^ Here, we shed light on how repeated ISR activation may transform microglia towards a neurodegenerative phenotype at the ultrastructural, molecular, and functional levels and provide a novel explanation for microglia-mediated neurotoxicity. Yet, our study has some limitations that should be addressed in future studies. For example, eIF2α phosphorylation has two critical outcomes: protein translation arrest and ISR gene expression downstream ATF4. We cannot pinpoint which of these outcomes underlies the neurodegenerative impact of ISR—though our study shows that ATF4 target FASN mediates the toxic effect of microglia and supports the latter. Future work will also be necessary to address several open questions raised by our study. How do other stress response pathways, such as those downstream of XBP1 or oxidative stress response, impact the dark microglia phenotype? What are the molecular mechanisms through which repeated ISR activation changes microglial identity? What are the lipid species that mediate the toxic impact of microglial ISR? What role do microglia-produced lipids play in Aβ and Tau pathology? What role does microglial ISR play in neurodevelopmental disorders^131,132^, where microglia and ISR have been heavily implicated? Overall, our research delineates a potentially targetable microglial phenotype whose functional outcome is neurodegenerative and expands the knowledge on the repertoire of microglial mechanisms of neurotoxicity.

## Author Contributions

A.F. and P.A. conceived and designed the study; A.F. performed mouse experiments, immunofluorescence, TRAP sequencing, and bioinformatics; A.F., L.A., and T.N. performed microglia isolation, qPCR, and Western blotting; A.F., L.A., S.A., H-J.P, J.M., D.A.-K., and P.C. designed and performed cell culture experiments; L.A. and J.L. performed single-nuclei sequencing; C.S., M.K., F.G.I., O.B., and M-E.T. designed, performed, and analyzed electron microscopy experiments; P.D. and N.J.H. designed and performed the lipidomics analysis; J.W.M. guided image analysis; J.D.M. and S.W. designed and performed synaptic terminal imaging and quantification; A.F. and P.A. wrote, and all authors edited the manuscript.

## Acknowledgments

We want to thank N. Heintz (Rockefeller University) and E. Klann (NYU) for generously providing the iPKR mice; N. Sonenberg and V. Sharma (McGill University) for generously providing the *Eif2^A^* mice; A.M. Goate (Icahn School of Mount Sinai) for generously providing the BV2 cell line; M. Parent and the brain bank at the CERVO Brain Research Center for generously providing the human brain sample; the ASRC Comparative Medicine Unit (CMU) for their assistance with mouse maintenance; the ASRC Epigenetics Core for flow cytometry and nucleic acid quality control; The Rockefeller University Genomics Core for sequencing and preliminary analysis; the CCNY Imaging Core and ASRC Live Imaging Core for microscopy and image analysis software; ASRC Mass Spectrometry core for mass spectrometry of ASV; A. Badimon (Rockefeller University), P. Casaccia (CUNY ASRC), and the ASRC Neuroscience Initiative for their helpful discussions and comments on the manuscript.

This work was supported by the CUNY Research Foundation, Alzheimer’s Association AARG-22-974642 (P.A.), Alfred P. Sloan Foundation JFRASE (P.A.), Provost’s Pre-Dissertation Science Research Fellowships (A.F., L.A.), CIHR Foundation FDN341846 (M.-E.T.), Tier II CRC in Neurobiology of Aging and Cognition CRC-2019-00407 (M.-E.T.), CFI John R. Evans Leaders Fund 39965 (M.-E.T.), National Multiple Sclerosis Society Grant TA-1905-34048 (S.W.), as well as generous support by Robin Chemers Neustein.

## Mice

Mice were housed at two to five animals per cage with a 12-hour light/dark cycle (lights on from 0700 to 1900 hours) at constant temperature (23°C) with *ad libitum* access to food and water. All animal protocols were approved by IACUC at CUNY Advanced Science Research Center CMU and performed according to NIH guidelines. *Eef1a1^LSL.NS3.eGFPL10a.iPKR/+^* iPKR mouse was generated as previously described^48,49^ and was generously provided by Drs. Eric Klann (New York University) and Nathaniel Heintz (The Rockefeller University). *Eif2s1^A/A^;* Tg*(fEif2s1*) *Eif2^A^* mouse was generously provided by Drs. Nahum Sonenberg and Vijendra Sharma (McGill University). *Cx3cr1^CreErt2/+(Litt)^* (#021160), 5xFAD (#030305), PS19 (#008169), and *Eef1a1^LSL.eGFPL10a/+^* TRAP (#030305) mice were purchased from The Jackson Laboratory. Wild-type CD-1 and C57BL/6 mice (for primary microglia, neurons, OPCs, and astrocytes) were purchased from Charles River Laboratories.

### Patients

The brain bank at CERVO Brain Research Center (QC, Canada) and handling of the post-mortem human tissues were approved by the Ethics Committee of the University of Victoria, Institut Universitaire en Santé Mentale de Québec, and Université Laval. Written consent was obtained for using post-mortem tissues, and all analyses were performed per the Code of Ethics of the World Medical Association.

### Mouse treatments

To activate Tamoxifen-inducible Cre-Ert2, all mice were gavaged at 4-6 weeks of age with six doses of 100 mg/kg of Tamoxifen (T5648, Sigma, St. Louis, MO) in corn oil (C8267, Sigma) with a separation of at least 48 hours between doses. Tamoxifen was dissolved directly in corn oil with 2 hours of incubation at 55°C.

ASV does not cross the blood-brain barrier (BBB), so we previously used an intracerebroventricular (i.c.v.) infusion of ASV^49^. Reasoning that this invasive surgery may initiate a non-specific, artificial response of microglia^43,133^, we established a less invasive drug delivery method by BBB permeabilization. Low-dose (1 μg/kg) VEGF injection can create a temporary window of BBB permeabilization and has been previously used to achieve efficient drug delivery to the brain^53,54^. Recombinant human VEGF (Fisher Scientific, 293-VE-050/CF) was reconstituted to a 100 ug/mL stock in water. Animals were intraperitoneally administered 1 μg/kg VEGF in 0.9% saline. ASV (Fisher Scientific, 50-202-9050) was reconstituted to 20mM stock in DMSO. 30 minutes after VEGF administration, we orally administered 5 mg/kg ASV in 1:9 PEG300 (MedChemExpress, 25322-68-3). Vehicle-treated animals received an intraperitoneal VEGF administration followed by 1:9 DMSO:PEG300 at 10ml/kg. For intermittent ISR, the injections were repeated biweekly for four weeks. In iPKR mice, treatment started at 4-5 months of age, in 5xFAD mice at 5 months, and in PS19 mice at 7 months.

C75 (Tocris, 2489) was dissolved in DMSO at 50mM. C75 was intraperitoneally injected at 10mg/kg in 0.9% saline every other day for 3 weeks. Vehicle-treated mice received 0.9% saline intraperitoneally at 10ml/kg. Evans Blue dye (Sigma, E2129-10G) was administered intraperitoneally 30 minutes after VEGF at 30mg/kg.

### Immunofluorescence staining and imaging of brain tissue samples

Animals were anesthetized with ketamine (120 mg/kg) and xylazine (24 mg/kg) and perfused transcardially with 10 ml PBS and 40 ml 4% paraformaldehyde (Electron Microscopy Sciences) as previously described^134^. Fixed brains were removed and dehydrated in 15% and 30% sucrose in PBS. Following dehydration, brains were frozen in Neg-50 (Life Technologies) on dry ice and stored at −80°C until further processing. Brains were cut using a cryostat, and 10- or 30-μm sections were mounted on Superfrost Plus microscope slides (Fisher Scientific), which were stored at −80°C until staining.

30-μm slides were washed with PBS, permeabilized with PBS + 0.2% Triton X-100 (PBST), and blocked with 2% normal goat serum in PBST. Slides were incubated with primary antibodies in 2% normal goat serum in PBST overnight at 4°C. Slides were then washed and incubated with Alexa Fluor-conjugated secondary antibodies (Alexa Fluor 488-, 546-, and 568-labeled goat anti-mouse, goat anti-rat, goat anti-chicken, goat anti-rabbit, or donkey anti-goat IgGs (H+L); 1:500, Life Technologies) in 2% normal goat serum in PBST for 1 h at RT. For nuclei and Aβ plaque staining, slides were washed and incubated with DRAQ5 (Thermo Scientific, 62-254, 1:2,500 in PBS) for 20 minutes and then Thioflavin S (Sigma Aldrich, T1892, 1% w/v Thioflavin S, 1:2,000 in PBS) for 2 minutes, respectively. Imaging was performed using a Zeiss LSM 800 Confocal Microscope (Zeiss, Oberkochen, DE). 20-μm z-stack confocal images were acquired at 2-μm intervals, with 20× objective. Image processing was performed using Zen 2011 software (Zeiss). Compressed z-stacked immunofluorescence images are used as representative images unless otherwise specified.

<u>For VGLUT2</u>, 10 µm coronal brain sections were blocked and permeabilized for 1h at room temperature in 10% normal goat serum/0.1M PB containing 0.3% Triton-X 100 (Sigma). Sections were then incubated with primary antibodies at room temperature overnight. The following day, sections were incubated with appropriate Alexa fluorophore-conjugated secondary antibodies (ThermoFisher Scientific), and Aβ plaques were visualized using 0.0005% Thioflavin S/0.1M PB (Sigma, T1892). After thorough washing, sections were mounted using vectashield without DAPI (Vector Laboratories, Burlingame, CA, USA). Three to four stained sections from each sample containing the somatosensory cortex were imaged on a Zeiss Observer 7 Laser Scanning Confocal microscope 900 equipped with an Airyscan 2 module, diode lasers (405 nm, 488 nm, 561 nm, 640 nm), and Zen Blue acquisition software (Zeiss; Oberkochen, Germany). Two to three randomly chosen 63x fields of view within the somatosensory cortex were acquired with three z-stack steps at 0.68 µm spacing for each hemisphere. Identical settings were used to obtain images from all samples within one experiment.

#### Primary antibodies

CD11b (1:1000, MCA711GT, Biorad, Hercules, CA), p-eIF2α (Ser 51, 1:300, Cell Signaling, 3398T), AT8 (1:500; Thermo Fisher Scientific, Cat No. MN1020B), IBA1 (Wako, Cat No. 011-27991, 1:2,000), BACE1 (Abcam, Cat No. ab108394, 1:500), NeuN (Millipore Cat No. ABN91MI, 1:1,000), CD31 (Invitrogen, 01-676-511, 1:250), and VGLUT2 (Synaptic Systems, 135404, 1:2000)

### Imaging quantification

Unless otherwise noted, z-stacked images from the brain tissues were analyzed by batch analysis using Imaris software (Bitplane, UK). In 5xFAD mice, a surface for plaques was first generated. <u>To quantify the</u> <u>plaque association of microglia</u>, a CD11b surface was generated, respectively, within 5 μm of the plaque surface. Next, surfaces were generated for the nuclei found within plaque-associated cells. The number of nuclei, number of plaques, and volume were recorded. The distal microglia were counted similarly for those more than 5 μm away from the plaque surface, and the numbers were normalized to the imaged area. <u>A CD11b surface was generated within plaques to quantify plaque infiltration by microglia</u>, and the volume was recorded. <u>For cytosolic p-eIF2α quantification</u>, the intensity of the p-eIF2α around microglial nuclei was recorded to capture endoplasmic reticulum staining. <u>For AT8 quantification</u>, an AT8+ surface was generated within CD11b+ microglia, CD31+ blood vessels, or NeuN+ neurons. <u>To quantify BACE1+</u> <u>dystrophic neurites</u>, a surface for BACE1 was generated.

<u>For the VGLUT2 density quantification</u>, image analyses were performed using ImageJ (NIH, version 1.53k) as described previously with some modifications^135^. In brief, first, a consistent threshold range was determined using sample images for each genotype and condition. All images were subjected to background subtraction. Then, single z-planes of the z-stacks (three z-planes per animal) were subjected to the same background subtraction and thresholding. Last, the total area of presynaptic inputs was measured from the thresholded images using the analyze particles function of ImageJ. Data from single z-planes was averaged for each z-stack of each field of view, and the mean of all fields of view from one animal was determined and normalized to control animals to assess changes in synaptic densities. For inhibitor experiments, manual thresholding blinded to condition and genotype for each channel within one experiment was performed (IsoData segmentation method, 85-255).

### Human sample cutting and preparation for scanning electron microscopy

Coronal hippocampal sections containing the entorhinal cortex, hippocampal head, CA1, CA2, and parahippocampal gyrus from an aged individual with Alzheimer’s disease (female, 79 years old, death by asphyxia; 18 h post-mortem interval) were obtained from the brain bank located at the CERVO Brain Research Center (QC, Canada). The brain was sectioned along the midline, and hemibrains were sectioned coronally into 2 cm thick samples. The latter were incubated for 3 days at 4°C in 4% paraformaldehyde (PFA), then stored at 4°C in phosphate-buffered saline (PBS; 50 mM, pH 7.4) with 15% sucrose and 0.1% sodium azide. Regions of interest were subsequently cut coronally into 50 µm-thick sections using a vibratome and stored at −20°C in cryoprotectant solution until immunostaining^18^.

Stained human brain samples containing the region of interest (CA1 and CA2 of the ventral hippocampus) were processed for scanning electron microscopy. Sections were incubated in 3% potassium ferrocyanide (cat# PFC232.250, BioShop, Burlington, ON, Canada) in phosphate buffer (PB; 100 mM, pH 7.4), combined (1:1) with 4% aqueous osmium tetroxide (cat# 19170, Electron Microscopy Sciences, Hatfield, PA, United States) for 1 h, then washed sequentially 5 min each in PB, 1:1 PB and double-distilled water (ddH_2_O), and ddH_2_O. Sections were then incubated in 1% thiocarbohydrazide (TCH; cat# 2231-57-4, Electron Microscopy Sciences) in ddH_2_O for 20 min, followed by 3 washes of 5 min in ddH_2_O. Sections were subsequently incubated in 2% osmium tetroxide (in ddH_2_O) for 30 min, then washed in ddH_2_O. Next, sections were dehydrated in ascending ethanol concentrations (2 times in 35%, 50%, 70%, 80%, 90%, and 3 times in 100%) for 10 min each, followed by 3 washes of 10 min in propylene oxide. Dehydrated sections were embedded in Durcupan ACM resin (cat# 44611–44614, MilliporeSigma) for 24 h, placed between two ACLAR® embedding sheets (cat# 50425-25, Electron Microscopy Sciences), and resin was polymerized at 55 °C for 5 days. The region of interest (CA1 and CA2 of ventral hippocampus) was excised, glued on a resin block, and cut into 73 nm thick ultrathin sections using a Leica ARTOS 3D ultramicrotome (Leica Biosystems).

### Mouse sample cutting and preparation for scanning electron microscopy

Animals were anesthetized at indicated ages with a ketamine (120 mg/kg) and xylazine (24 mg/kg) mixture and perfused transcardially with 10 ml PBS and 150 ml 0.6% glutaraldehyde/4% paraformaldehyde (Electron Microscopy Sciences). Brains were post-fixed overnight in 0.6% glutaraldehyde/4% paraformaldehyde and washed with PBS. Brains were cut with a vibratome into 50-um sections stored in cryoprotectant (30% glycerol and 30% ethylene glycol in PBS) until further processing.

Brain sections from female 5xFAD (n=1), male 5xFAD-*Eif2a^A^* (n=1), female 5xFAD-iPKR (n=1), and female iPKR (n=1) mice were processed for scanning electron microscopy. Coronal sections containing the deep layers of the isocortex (Bregma −0.46 mm to −0.70 mm)^136^ from 5xFAD, 5xFAD-*Eif2a^A^*, and iPKR animals were selected. Sagittal sections containing deep layers of the isocortex (Bregma 2.88 mm to 3.12 mm)^136^ from 5xFAD and 5xFAD-iPKR animals were selected. Sections were incubated in 3% potassium ferrocyanide in PB, combined (1:1) with 4% aqueous osmium tetroxide for 1 h, then washed sequentially 5 min each in PB, 1:1 PB and ddH_2_O, and ddH_2_O. Sections were then incubated in 1% TCH in ddH_2_O for 20 min, followed by 3 washes of 5 min in ddH_2_O. Sections were then incubated in 2% osmium tetroxide (in ddH2O) for 30 min, then washed in ddH_2_O. Next, sections were dehydrated in ascending ethanol concentration (2 times in 35%, 50%, 70%, 80%, 90%, and 3 times in 100%) for 5 min each, followed by 3 washes of 5 min in propylene oxide. Dehydrated sections were embedded in Durcupan ACM resin for 24 h, placed between two ACLAR® embedding sheets, and resin was polymerized at 55 °C for 3 days. The region of interest (deep layers of the cortex) was excised, glued on a resin block, and cut into 73 nm thick ultrathin sections using a Leica ARTOS 3D ultramicrotome (Leica Biosystems).

### P-eIF2*α* immunostaining of samples for scanning electron microscopy

Immunostaining was performed on post-mortem female human hippocampal samples (n=1) and female 5xFAD (Jackson laboratory) mouse samples (n=1). Human brain coronal section samples containing the CA1 and CA2 of the ventral hippocampus were selected for processing. Mouse brain sagittal section samples containing the deep layers of the isocortex were selected for processing (Bregma 2.88 mm to 3.12 mm) (Franklin and Paxinos, 2013). Sections were washed for 7 minutes in PBS 3 times. Sections were then incubated in sodium citrate buffer (10 mM, pH 6.0) for 15 min at 70°C, then allowed to cool to room temperature (RT), followed by 3 washes of 7 min in PBS. Sections were subsequently quenched in 0.3% ddH_2_O (Fisher Scientific, lot# 202762) in PBS for 5 min, followed by 3 washes of 7 min in PBS. Afterward, the sections were incubated in 0.1% NaBH_4_ in PBS for 30 min, followed by 3 washes of 7 min in PBS. Sections were next incubated in a blocking buffer containing 10% normal goat serum (Jackson ImmunoResearch Labs, Baltimore, USA cat# 005-000-121), 3% bovine serum albumin (Sigma-Aldrich, Oakville, cat# 9048-46-8,), and 0.01% Triton X-100 in Tris-buffered saline (TBS; 50 mM, pH 7.4) for 1 h at RT. They were incubated overnight in a blocking buffer solution with a rabbit anti-p-eIF2α primary antibody (1:300; Cell Signaling, 3398T) at 4°C. The following day, the brain sections were washed 3 times for 7 min with TBS, then incubated with a biotinylated goat anti-rabbit polyclonal secondary antibody (1:300; Jackson ImmunoResearch, 111-066-046) in TBS for 2 h at RT. Next, sections were washed in TBS and incubated with an avidin-biotin complex solution (ABC; 1:100; Vector Laboratories, PK-6100) in TBS for 1 h at RT. The staining was revealed for 90 seconds with 0.05% 3,3’-diaminobenzidine (DAB; Millipore Sigma, D5905-50TAB) and 0.015% H2O2 diluted in Tris buffer (TB; 0.05M, pH 8.0).

### Scanning electron microscopy imaging and ultrastructural identification of dark microglia

Ultrathin-processed sections were collected on a dust-free silicon nitride chip, glued on specimen mounts, and inserted into a Zeiss Crossbeam 350 focused ion beam scanning electron microscope (FIB-SEM). The regions of interest (CA1 and CA2 of the ventral hippocampus for human sections, deep layers of the isocortex region for mouse sections) were identified using backscattered electrons (ESB) and secondary electrons (SE2) detectors at 10 kV for initial focus, then at 1.4 kV when samples were at a sufficient working distance to perform high-resolution focus on the brain ultrastructure. These steps were controlled using SmartSEM software (Zeiss). Microglia were identified by their distinctive nuclear heterochromatin pattern and specific ultrastructure features such as long and narrow stretches of endoplasmic reticulum (ER) cisternae and lipid inclusions (e.g., lysosomes and lipofuscin granules), as well as association with pockets of extracellular space^137^. Dark microglia were differentiated from typical microglia by their distinctive electron-dense cytoplasm and nucleoplasm, loss of the nuclear heterochromatin pattern, and various cellular stress markers like dilated ER cisternae and altered mitochondrial cristae ultrastructure^17,19,137^. Dark microglia were then imaged at 5 nm resolution and 1.4 kV, and the images were exported as TIFF files using the Zeiss ATLAS Engine 5 software (Fibics).

### Qualitative electron microscopy analysis

For dark microglia interaction analysis, using the 5 nm images, the area examined, the number of dark microglia, and their proximity to dystrophic neurites and plaques were manually recorded.

For the quantification of p-eIF2α+ cell types, using all the 5nm images from a stained female 5xFAD (n=1), the complete and incomplete cells in each image, if the cell had p-eIF2α staining, and the subcellular location of the staining were manually recorded.

For the quantification of the dark and intermediate microglia density, using the 25nm images, the number of the dark microglia population and the area examined were manually recorded.

All analyses were conducted by an experimenter blind to the conditions.

### Translating Ribosome Affinity Purification (TRAP)

We chose to use TRAP because it can unmask changes at the translational level that may not be evident at the transcriptional level—especially relevant for ISR genes^20,33^. It also overcomes limitations associated with cell isolation^43^, where distinct microglia populations may disproportionately respond to or not survive the isolation process. The following genotypes for biological groups were used:

_TRAP: *Cx3cr1CreErt2/+; Eef1a1LSL.eGFPL10a/+*_

*Eif2^A^*: *Cx3cr1^CreErt2/+^; Eef1a1^LSL.eGFPL10a/+^*;*Eif2s1^A/A^;* Tg*(fEif2s1*)

5xFAD: 5xFAD;*Cx3cr1^CreErt2/+^; Eef1a1^LSL.eGFPL10a/+^*

5XFAD*Eif2^A^*: 5xFAD;*Cx3cr1^CreErt2/+^; Eef1a1^LSL.eGFPL10a/+^*;*Eif2s1^A/A^;* Tg*(fEif2s1*)

_iPKR: *Cx3cr1CreErt2/+; Eef1a1LSL.NS3.eGFPL10a.iPKR/+*_

_5xFADiPKR: 5xFAD;*Cx3cr1CreErt2/+; Eef1a1LSL.NS3.eGFPL10a.iPKR/+*_

Microglia-specific ribosomal profiling was performed from *Cx3cr1^CreErt2/+(Litt)^* mice crossed to *Eef1a1^LSL.eGFPL10a/+^* TRAP or *Eef1a1^LSL.iPKR/+^* iPKR mice, which contain the gene encoding eGFP-L10a. Mice were euthanized with CO_2_, and brain regions of interest were dissected and flash-frozen for TRAP. Ribosome-associated mRNA from microglia was isolated from each region as previously described,^42^ where each sample corresponds to a single mouse. We used the cortex, which is among the brain regions most affected in AD and is significant to memory and higher functions^138–140^. Briefly, the brain tissue was thawed in an ice-cold Wheaton 33 low extractable borosilicate glass homogenizer containing 1 mL cell-lysis buffer (20 mM HEPES-KOH, pH 7.3, 150 mM KCl, 10 mM MgCl2, and 1% NP-40, 0.5 mM DTT, 100 μg/ml cycloheximide, and 10 μl/ml rRNasin (Promega) and Superasin (Invitrogen). After manually homogenizing the samples with 3-5 gentle strokes using the PTFE homogenizer (grinding chamber clearance 0.1 to 0.15mm). Samples were then homogenized in a motor-driven overhead stirrer at 900 r.p.m. with 12 complete strokes. The samples were transferred to chilled Eppendorf tubes, and a post-nuclear supernatant was prepared by centrifugation at 4°C, 10 minutes, 2,000 x g. NP-40 (final concentration = 1%) and DHPC (final concentration = 30 mM) were added to the supernatant. A post-mitochondrial supernatant was prepared by centrifugation at 4°C, 10 minutes, 16,000 x *g.* 200uL Streptavidin MyOne T1 Dynabeads (Invitrogen), conjugated to 1 μg/μl biotinylated Protein L (Pierce) and 50 μg each of anti-eGFP antibodies Htz-GFP-19F7 and Htz-GFP-19C8, bioreactor supernatant (Memorial-Sloan Kettering Monoclonal Antibody Facility) were added to each supernatant. After overnight incubation at 4°C with gentle end-over-end rotation, the unbound fraction was collected using a magnetic stand. The polysome-bound beads were washed with the high-salt buffer (20 mM HEPES-KOH, pH 7.3, 350 mM KCl, 10 mM MgCl2, 1% NP-40, 0.5 mM DTT, and 100 μg/ml cycloheximide). RNA clean-up from the washed polysome-bound beads and 5% of the unbound fractions was performed using RNeasy Mini Kit (Qiagen) following the manufacturer’s instructions. RNA integrity was assayed using an RNA Pico chip on a Bioanalyzer 2100 (Agilent, Santa Clara, CA), and only samples with RIN>9 were considered for subsequent analysis.

### Library preparation and sequencing

Libraries were prepared from 0.6-1.0ng starting RNA using SMART-Seq® mRNA LP (Takara, 634768), multiplexed using Unique Dual Index Kit 1–96 (Takara, 634752), and size-selected using NucleoMag® NGS Clean-up and Size Select beads (Takara 744970.5) as per the manufacturer’s recommendations. Concentrations and quality controls were performed using High Sensitivity RNA TapeStation and High Sensitivity DNA D5000 TapeStation. Multiplexed libraries were directly loaded on NextSeq 500 (Ilumina) with single-read sequencing for 75 cycles. Raw sequencing data were processed by using Illumina bcl2fastq2 Conversion Software v2.17.

### Bioinformatic analysis of TRAP sequencing

Quality control was performed on raw sequencing reads with Trim Galore, followed by alignment to the mouse genome (mm10) with HISAT2 (version 2.2.1) using default parameters^141^. featureCounts (version 2.0.3) was used to obtain the read counts per gene against the mm10 GENCODE annotation^142^. We pooled two batches of TRAP sequencing samples before differential gene expression analysis. For differential expression analysis, read counts were input into DESeq2^143^ (version 1.38.3). Mapping statistics, including read count, quality assessment, and read alignment, were performed in parallel. The raw counts were normalized using the variance stabilizing transformation (vst) DESeq2 package and further transformed into z-scores for heatmap visualization. Principal component analysis (PCA) was performed on the top 500 most variable genes across all samples based on the VST data for outlier assessment.

As microglial enrichment in iPKR mice was lower than in TRAP mice (likely due to lower expression of eGFP-L10a downstream of the 2A sequence^48^), we performed pairwise comparisons within each line to avoid artificial enrichment/depletion of microglia-specific genes. Namely, 5xFAD, *Eif2^A^*, and 5XFAD*Eif2^A^* samples were compared to TRAP samples. ASV-treated iPKR and 5xFADiPKR samples were compared to vehicle-treated iPKR samples. To identify differentially enriched genes, we used a cutoff of the p-value < 0.05 and mean expression > 20 (DESeq2). We did not select TRAP-enriched genes against the unbound fraction to fully represent ubiquitously expressed stress-associated and metabolic genes. All heat maps and scatter plots for bulk sequencing were made using R (v3.1.1; https://www.R-project.org). For all heatmaps, the expression of each gene in vst was normalized to the mean across all samples (z-scored). Pathway and upstream regulator analyses were performed using Ingenuity Pathway Analysis software (Qiagen). P values and z scores are represented in balloon graphs and were made using R (v3.1.1; https://www.R-project.org).

### Isolation of microglial nuclei by FANS for single nuclei sequencing

Microglial nuclei were isolated based on the eGFP-L10a fluorescence of newly formed ribosomes in the microglia nucleoli as described in Ayata et al.^43^ Briefly, one male and one female mouse for each sample were sacrificed by cervical dislocation. Brain regions were quickly dissected and homogenized in 0.25 M sucrose, 150 mM KCl, 5 mM MgCl2, 20 mM Tricine pH 7.8 with a glass Dounce homogenizer (1984-10002, Kimble Chase, Vineland, NJ). The buffers were supplemented with 10 μl/ml RNasin, Superasin, and EDTA-free protease inhibitor cocktail (11836170001, Roche). The homogenate was then spun through a 29% iodixanol cushion. The resulting nuclear pellet was resuspended in 0.25 M sucrose, 150 mM KCl, 5 mM MgCl2, 20 mM Tricine pH 7.8, supplemented with 10 μM DyeCycle Ruby (V10304, Invitrogen), and 10% donkey serum (017-000-121, Jackson Immunoresearch, West Grove, PA). Microglial nuclei were sorted in a BD FACS Aria cell sorter by gating for the lowest DyeCycle Ruby, which indicates nuclei singlets and a high GFP signal. For single nuclei RNA sequencing, isolated nuclei were used immediately for 10X sequencing.

### Single nuclei sequencing

Single-nucleus RNA-seq (snRNA-seq) was performed on isolated microglial nuclei (at least 5,000 nuclei) using the Chromium platform (10x Genomics, Pleasanton, CA) with the 3′ gene expression (3′ GEX) V2 kit, using a targeted input of ∼8,000 nuclei per sample. In brief, gel-bead in emulsions (GEMs) were generated on the sample chip in the Chromium controller. Barcoded cDNA was extracted from the GEMs using Post-GEM RT-cleanup and amplified for 12 cycles. Amplified cDNA was fragmented and subjected to end-repair, poly-A-tailing, adaptor ligation, and 10X-specific sample indexing following the manufacturer’s protocol. Libraries were quantified using Bioanalyzer (Agilent) and QuBit (Thermofisher) analysis and then sequenced in paired-end mode on a NextSeq 2000 instrument (Illumina, San Diego, CA) targeting a depth of 30,000 reads per nucleus.

### Bioinformatic analysis of single-nuclei sequencing

.fastq files containing barcoded reads for single-nuclei sequencing were aligned to the mouse mm10 genome and processed into filtered feature barcode matrices (Market Exchange Format) using CellRanger Pipeline v7.1.0 with default parameters. To capture unspliced pre-mRNA in the single-nucleus RNA expression assay, intronic regions in the 10X Cell Ranger mm10 v1.2.0 reference were marked as exonic as suggested by 10X for pre-mRNA reference generation. Among the 27,950 sequenced nuclei, the median number of UMIs detected per nucleus was 2,375, with a median of 1,367 genes per nucleus.

The filtered feature barcode matrices were then used downstream processing with the Seurat bundle^144^. First, the Seurat object was initialized with the raw (non-normalized data) for each dataset. Features detected in less than 5 cells and cells with less than 150 unique RNAs or more than 5% mitochondrial RNA were excluded using min.cells, subset, and percent.mt functions. Then the following clustering pipeline steps were applied: NormalizeData(), FindVariableFeatures(), ScaleData(), RunPCA(), FindNeighbors(), and FindClusters(). Next, the non-microglial clusters that do not express *Hexb* were excluded. After this pipeline was applied to all datasets, they were integrated using the following pipeline: FindIntegrationAnchors() with dims 1:20, IntegrateData() with dims 1:2, ScaleData(), FindVariableFeatures(), RunPCA(), RunUMAP(), FindNeighbors(), and FindClusters() with resolution 1.2. Subset markers were identified with FindAllMarkers() with min.pct = 0.1 and logfc.threshold = 0.2. The number of cells in each cluster was calculated using Idents(). Dot plots were generated by the DotPlot() function, dim plots with the DimPlot() function, and feature plots with the FeaturePlot() function. Homeostatic, Stressed, DAM1, DAM2, IFN, and High-metabolism clusters were formed by merging clusters that share gene expression and gene ontologies. Gene ontology enrichments were calculated using EnrichR^145–147^.

### Mass spectrometry for ASV

Animals were sacrificed by cervical dislocation, and brain tissue was flash-frozen in liquid N_2_ until further processing. Samples were homogenized in ice-cold HPLC-grade 80% methanol in glass homogenizers using automatic pestle mortar at 4°C, centrifuged at 14,000g at 4°C for 5 minutes, and stored at −80°C until analysis. The samples were analyzed using Bruker’s Daltonics maXis-II UHR-ESI-QqTOF mass spectrometer coupled to a Thermo-Scientific Ultimate-3000 UHPLC system. Agilent Acclaim 120 Agilent’s 300SB-C18 2.1 x 75mm column was used for separation at 300C and the flow rate at 150 μL/min. The gradient used was 0–2 min 2% solvent B (acetonitrile, 0.15% formic acid) and 98% solvent A (water, 0.15% formic acid) followed by a gradient 2–50% B from 2 to 18 min, 50–95% B from 18 to 20 min, then held at 95% B from 20 to 25 min. All experimental data were acquired over the range m/z 50– 1500 in positive ion mode. ASV concentrations were calculated using a standard dilution series.

### Acute isolation of adult microglia

Adult microglia were isolated using a protocol adapted from Badimon et al.^74^ Animals were sacrificed by cervical dislocation, and the cortex was dissected for mechanical homogenization in HBSS (Sigma, H6648-6X500ML). All steps were performed on ice or at 4℃ to minimize *ex vivo* microglia activation.^43,74^. Dissociated cells were filtered through a 70-μm mesh filter (Fisher Scientific, 22363548). Myelin removal was performed using Percoll (pH 7.4) density gradient separation. The homogenate was supplemented with 90% Percoll (17-0891-02, Amersham, Amersham, UK) with PBS (pH 7.4). The resulting homogenate in 21% Percoll gradient was centrifuged at 500*g* for 15 min at 4 °C. Supernatant was removed, and the cell pellet was resuspended in filter-sterilized MACS buffer (1X PBS, 0.5%BSA, 2mM EDTA). CD11b+ microglia/myeloid cells were selected by anti-CD11b-coated microbeads (130-093-636, Miltenyi) with the LS columns (Miltenyi, 130-042-401) and QuadroMACs separator following the manufacturer’s recommendations. The ensuing microglia suspension was centrifuged at 300g for 15 minutes and resuspended in MACS buffer for manual counting with a hemocytometer.

### BV2 cell culture and conditioned media preparation

BV2 cells were chosen over primary microglia for conditioned media experiments for two reasons: 1. They can be grown in high numbers to produce sufficient treatments for multiple experiments with technical replicates. 2. They survive well despite several washes and media changes. The BV2 mouse microglial cell line was kindly provided by Dr. Alison Goate (Icahn School of Medicine at Mount Sinai). The BV2 cells were cultured in Dulbecco’s modified Eagle’s medium (DMEM) (Gibco #11965) supplemented with 5% heat-inactivated <u>fetal bovine serum</u> (FBS, Gibco #16140) and 100 U/ml penicillin-streptomycin (Pen/Strep, Gibco #15140). The BV2 cells were routinely tested for mycoplasma contamination using MycoAlert PLUS mycoplasma detection kit (Lonza).

For conditioned media (CM) preparation, BV2 cells were treated with 10μM salubrinal/SLB (Sal 003, Tocris #3657, prepared as 50 mM stock in DMSO) or vehicle DMSO (Fisher Scientific #BP231) for 24 hours. To prepare CM with C75 (Tocris, 2489), BV2 cells were treated with 25μM C75 24 hours after plating and 1 hour before treatment with DMSO or SLB. The DMSO-/SLB-containing medium was removed, the cells were washed with PBS, and a fresh recipient cell medium was added to the cells. For example, if the target cell was OPCs, a fresh SATO medium was given (see below). This CM for each cell type was collected after 6 hrs. To ensure that this conditioned media represented ISR-activated microglia, the persistent (albeit reduced) expression of *Atf4* in BV2 cells after media change was confirmed by qPCR (Supplementary Fig. 12b). Each target cell media was tested to verify that they did not artificially induce ISR in microglia (Supplementary Fig. 12c). The CM was filtered using the Steriflip® vacuum filter tubes (Millipore, #SCGP00525), flash frozen, and stored at −80°C until use.

Lipid depletion of CM was done using Cleanascite™ (Biotech Support Group #X2555) following the manufacturer’s recommendations. Briefly, 1:4 of the Cleanascite™ reagent was added to the CM, incubated for 10 min at room temperature, mixed periodically, and then centrifuged at 1,600 g for 2 min. The supernatant was carefully collected, centrifuged again, and filtered through a 0.22 µm-syringe filter (Avantor, 76479-024) before applying it to recipient cells.

### A*β* uptake experiments

Aβ was prepared by aggregating fluorescently labeled Aβ peptides (Beta-Amyloid 1-42, HiLyte Fluor 488-labeled, Anaspec AS-60479-01), as previously described^148^. Lyophilized Aβ was resuspended at 2 mg/mL in PBS with 10 mM NaOH at pH 7.5. Protein was shaken at 1500 rpm overnight at 37°C.

BV2 cells were treated for 24 hours with 10 μM salubrinal/SLB (Sal 003, Tocris #3657, prepared as 50 mM stock in DMSO) or vehicle DMSO (Fisher Scientific #BP231). After 24 hours, SLB/DMSO media was replaced with 0.1 μg/mL Aβ for 6 hours in a live cell imaging system (Incucyte SX1 Live-Cell Analysis System, Sartorius), with imaging in both phase and green channels at 10X magnification every 15 minutes. Total green cell and total cell numbers were recorded (both quantified by Incucyte Live-Cell Analysis AI software manufacturers’ settings).

### Primary OPC cultures

Primary mouse OPCs were isolated from the brain of C57BL/6 mice at postnatal day P7 through immunopanning with a rat anti-mouse CD140a antibody, recognizing PDGFRα, as previously described^106^. Briefly, cerebral cortices were digested in papain solution for 20 min at 37°C, followed by trituration and filtration. After a series of selection process (2 negative selection with BSL1 followed by 1 positive selection with CD140a), cells were cultured in SATO medium (DMEM containing 100 μg/ml bovine serum albumin (BSA), 10 μg/ml apotransferrin, 16 μg/ml putrescine, 62.5 ng/ml progesterone, 39.6 ng/ml sodium selenite, 5 μg/ml insulin, 1 mM sodium pyruvate, 2 mM l-glutamine, 100 U/ml penicillin, 100 μg/ml streptomycin, B27 Supplement, 5 μg/ml N-acetyl-cysteine, Trace Element B, 10 ng/ ml biotin, 50 μM forskolin) supplemented with growth factors (10 ng/ml PDGF-AA and 20 ng/m/ bFGF), and maintained in a poly-D-lysine-coated 12-well plate (Corning, #3513) in a 37°C, 5% CO_2_ incubator for further expansion. For qPCR, cells were plated onto poly-D-lysine coated 24-well plates (Corning) at a density of 3×10^4^ cells/well. Once they reached 80% confluency, OPCs were treated with CM for 6 hrs, lysed using TRIzol™ reagent (Fisher Scientific #15596026), flash frozen, and stored at −80°C for RNA isolation. For immunocytochemistry, cells were plated onto poly-D-lysine coated CC^2^-eight-chamber slides at a density of 3×10^5^ cells/well. Once they reached 80% confluency, OPCs were treated with CM for 6 or 24 hrs and fixed with 4% paraformaldehyde.

### Primary astrocyte cultures

Primary astrocytes were isolated from CD-1 mice at P3, as previously described^149^. Mouse dissections were done in ice-cold HBSS (Gibco, 14065056). After dissection and removal of meninges, cortex tissue was placed in 2 ml of warm astrocyte media (Advanced DMEM/F12 (Gibco #12634010) supplemented with 5% FBS, 100 U/ml GlutaMAX™ (Gibco #35050061), and 100 U/ml penicillin-streptomycin). The tissue was mechanically dissociated using a 16-gauge BD Precision Glide™ needle attached to a 1mL syringe, followed by gentle pipetting. Dissociated cells were then passed through a 70-μm nylon mesh filter, centrifuged at 400 g for 5 minutes, and resuspended in 5ml warm fresh astrocyte media. Resuspended cells were plated onto T175-cm flasks (Millipore #CLS431466) coated with 0.1% gelatin (Millipore-Sigma #1288485-500MG) and placed in the cell culture incubator at 37°C, 5% CO. The growth of astrocytes was monitored over a week and used for experiments accordingly. If not in use, the astrocytes were frozen in 90% FBS with 10% DMSO and kept at −80°C for long-term storage. For qPCR, astrocytes were plated in a 12-well plate and treated with CM for 6 hours. After 6 hours, cells were dissociated with TRIzol™ and stored at −80°C for RNA isolation.

### Primary neuron cultures

Primary cultures of cortical neurons were isolated from mouse embryos on embryonic day E16 as described previously^107^ with slight modifications. Briefly, embryonic cortices were digested with 0.025% trypsin–EDTA for 15 min at 37°C, followed by incubation with NM10 medium (DMEM with 10% fetal bovine serum) for 5 min at room temperature. After a series of dissociation processes (trituration followed by filtration and centrifugation), cells were resuspended in a neurobasal medium supplemented with B27, 2 mM L-glutamine, and 100 U/mL penicillin, 100 μg/mL streptomycin and plated. At 3 days post-plating, cells were exposed to an anti-mitotic agent (5 μM AraC) for 16 hours to avoid contamination of non-neuronal cells, followed by fresh media change. Cells were then cultured at 37°C in a 5% CO_2_ incubator until 14 days in vitro (DIV), and the medium was replaced every other day. Cells were plated onto poly-d-lysine-coated 48-well plates at a density of 3.0 × 10^4^ cells/well. For qPCR, cells were treated with CM for 6 hours and imaged every hour until cell collection by the Incucye live imaging platform (Sartorius) in the incubator at 37°C at 5% CO. After 6 hours, cells were dissociated with TRIzol™ and stored at −80°C for RNA isolation. For immunostaining, cells were treated for 24 hours and were fixed with 4% PFA.

### Multi-electrode array recordings

For testing endogenous spontaneous electrical neuronal activity, cells were plated onto polyethyleneimine (PEI)/laminin-coated Axion 48-well multi-electrode array plates at a density of 7.5 × 10^4^ cells/well. After two recordings, neurons were given the CM, and their endogenous electrical activity was recorded using a Maestro Pro multi-well multi-electrode array system (Axion Biosystems) every 30 minutes for 2 hours. Briefly, on the day of the experiment, cells were challenged with conditioned media for up to 6 hours, and spontaneous neuronal activity was recorded for 10-15 minutes at 1 hour post-exposure. Before activity acquisition, cells were allowed to equilibrate for 5 min. Signals were simultaneously acquired across 768 channels (16 electrodes per well, 48-well plate), and data acquisition was managed with Axion’s integrated software, AxiS Navigator.

### Quantitative PCR (qPCR)

To obtain RNA from cell culture, cell media was removed, and cells were washed with ice-cold PBS twice and lysed in Trizol (Invitrogen) for 5 minutes on ice. Phase separation was performed using chloroform:isoamyl alcohol 49:1 (Sigma). RNA was precipitated using isopropanol, sodium acetate, and GlycolBlue (Invitrogen). The pellet was washed with freshly prepared, ice-cold 80% ethanol, air-dried, and resuspended in nuclease-free water. Up to 500ng of RNA was used as starting material for cDNA synthesis using Applied Biosystems High-Capacity RNA-to-cDNA Kit (Applied Biosystems) following the manufacturer’s recommendations. cDNA was then used as input for qPCR using PerfeCTa FastMix II Low ROX (QuantaBio) following the manufacturer’s recommendations. Cycle counts for mRNA quantification were normalized to *Hexb* unless otherwise stated. Quantification (RQ=2^ΔCt^) and relative expression (experimental RQ/average control RQs) for each gene were calculated.

Probes: *Atf4* (Mm00515325_g1), *Ppp1r15a*: (Mm01205601_g1), *Sqstm1* (Mm00448091_m1), *Asns* (Mm00803785_m1), *Hexb*: (Mm01282432_m1), *Pdgfra* (Mm00440701_m1), *Gapdh*: (Mm99999915_g1), *Actb* (Mm02619580_g1).

### Protein preparation from BV2 cells and Western blot analysis

Cell pellets containing over 10^6^ BV2 cells were resuspended in RIPA buffer supplemented with PhoSTOP phosphatase inhibitor (Sigma, 4906845001), a cOmplete™, EDTA-free Protease Inhibitor Cocktail (Sigma, 11873580001), and 2.7uL/10mL Benzonase (MilliporeSigma, 712053) at 10,000 cells/uL and incubated on ice for 1 hour with gentle vortexing every 15 minutes. NaCl was added to achieve a final concentration of 0.385M, and the samples were nutated at 4°C for 1 hour. Samples were centrifuged at 14,000 g, 4°C for 15 minutes. Protein concentration was quantified using a BCA protein assay kit (Pierce, 23225) according to the manufacturer’s recommendations. The lysate was mixed with NuPage LDS Sample Buffer (Invitrogen) to achieve 1X and DTT (Sigma) to achieve 0.1M and heated for 10 minutes at 70°C. 6.5μg of the sample was loaded into a 4-12% NuPAGE gel (Invitrogen) and ran at 80V. Samples were transferred to (PVDF membranes, Invitrogen) for 2 hours at 25V in an ice-cold transfer buffer, following the manufacturer’s recommendations. Membranes were incubated with blocking buffer (5% non-fat milk in TBS-Tween) for 1 hr at room temperature and primary antibodies overnight at 4°C. Membranes were washed with TBS-Tween and incubated with secondary HRP-coupled antibodies (Invitrogen, 1:10,000) for 1 hr at room temperature. Proteins were detected using the Western Lightning Plus-ECL (Perkin Elmer), following the manufacturer’s recommendations, and chemiluminescence images were processed using the ChemiDoc Imaging System (Bio-Rad).

#### Primary antibodies

p-eIF2a (1:500 in 5% BSA, CST #9721), eIF2a (1:1000 in 5% BSA, CST, #2103).

### Immunocytochemistry of neurons and OPCs

Cells were cultured on CC^2^ (8-well) chamber slides (Thermo Fisher Scientific, 154941PK). 24 hours after conditioned media treatment, they were fixed with 4% PFA for 15 minutes at room temperature and washed three times with ice-cold PBS. Fixed cells were incubated in blocking buffer (5% normal goat serum and 0.02% Triton in PGBA buffer [0.1 M phosphate buffer, 0.1% gelatin, 1% bovine serum albumin, 0.002% sodium azide]) for 1 hr at room temperature and then incubated with primary antibodies in blocking buffer overnight at 4°C in the dark. Cells were washed three times with ice-cold PBS and incubated with Alexa Fluor-conjugated secondary antibodies (Alexa Fluor-labelled 488 goat anti-rabbit, 546-goat anti-mouse, and 647 goat anti-rat, IgGs (H+L); 1:500, Life Technologies) in blocking buffer for 1 hr at room temperature. Slides were then washed, mounted using a Prolong Gold anti-fade mounting medium containing DAPI (Invitrogen, P36931), and dried overnight in the dark at room temperature. Imaging was performed using a fluorescence microscope (EVOS M5000 Invitrogen) at 10x magnification. For the analysis, ImageJ (NIH, version 1.53k) was used as described previously^150^. Briefly, the number of DAPI-, cCASP3-, OLIG2, and NeuN-positive cells were counted in each image (n=5-8 images per sample)

#### Primary antibodies

OLIG2 (Millipore, MABN50, 1:500); NeuN (Millipore Cat No. ABN91MI, 1:1,000), cCASP3 (Cell Signaling, 9661, 1:1,000).

### Untargeted full-scale lipidomics

A crude lipid fraction was extracted from cell media using a slightly modified Bligh and Dyer procedure^151,152^. In brief, cell media were diluted with ddH_2_O to make up a final volume of 1 mL. They were then added 2.9mL of methanol/dichloromethane (2:0.9, v/v) containing the twelve internal standards: Cer d18:1/12:0 −6 ng/mL, SM d18:1/12:0 −0.5 ng/mL, GlcCer d18:1/12:0 −3.0 ng/mL, LacCer 18:1/12:0 −10.0 ng/mL, d5-DAG d16:0/16:0 −15 ng/mL, d5-TAG 16:0/18:0/16:0 −2 ng/mL, cholesteryl-d7 ester 16:0 −30 ng/mL, PA d12:0/12:0 −500 ng/mL, PC 12:0/12:0 −0.2 ng/mL, PE d12:0/12:0 −2 ng/mL, PG d12:0/12:0 −200 ng/mL, PS d12:0/12:0 −900 ng/mL in a glass tube. To obtain a biphasic mixture, an additional 1mL of ddH_2_O and 900μL dichloromethane was added and vortexed. The resultant mixture was incubated on ice for 30 min and centrifuged (10 min, 3,000 g, 4°C) to separate the organic and aqueous phases. The organic phase was collected and stored at −20°C. Just before analysis, 1mL of the organic layer was dried using a nitrogen evaporator and re-suspended in 150µl of running solvent (dichloromethane:methanol, 1:1) containing 5mM ammonium acetate and 5mg/mL of ceramide C17:0 to track the instrument performance. Lipid analysis was conducted in MS/MS^ALL^ mode on a TripleTOF 5600 (AB Sci-ex, Redwood City, CA). Samples (50μL injection volume) were directly infused by HPLC at a constant flow rate of 7 µL/min using an LC-20AD pump and SIL-20AC XR autosampler (Shimazu, Canby, OR). The mass spectrometer was operated at a mass resolution of 30,000 for the time of flight (TOF) MS scan and 15,000 for the product ion scan in the high sensitivity mode and automatically calibrated every 10-sample injections using APCI positive calibration solution delivered via a calibration delivery system (AB SCIEX). The TOF MS and MS/MSALL data obtained were post-aligned to internal standards using Analyst TF 1.8 (AB Sciex) with a mass error of less than 5 ppm. LipidView (version 1.3, AB SCIEX, Concord, Ontario, Canada AB Sciex) database was used for the identification and annotation of lipid species. Lipid identifications were validated using a pooled sample extract based on the precursor and fragment matchings from the experimental pooled sample runs^152^. These pre-validated lipid species were then quantified in each experimental sample using MultiQuant software (version 3.0, AB Sciex, Concord, ON, Canada). Each lipid species response was normalized to its corresponding internal standard response, and these normalized values were used for statistical analysis.

### Statistical analysis

Statistics were analyzed using GraphPad Prism v9.5.1, and significance was determined at p-value < 0.05. For two-group comparisons, an unpaired t-test was used; for comparisons with more than two groups, a one-way ANOVA was performed. All statistical analyses were two-tailed. Normal distribution was assessed by the Shapiro-Wilk (SW) normality test. Grubbs’ test was used to identify outliers. Normally distributed data with unequal variance was analyzed with Welch’s correction. Data that was not normally distributed was analyzed using a two-tailed Mann-Whitney U test. No statistical methods were used to predetermine the sample size, but our sample sizes are similar to those generally employed in the field.

### Data availability

TRAP sequencing, single-nuclei sequencing, electron microscopy, and lipidomics datasets will be deposited to public databases and will be available upon publication. Other datasets and any additional information are available from the lead contact upon request.

---

**Supplementary Figure 1.**
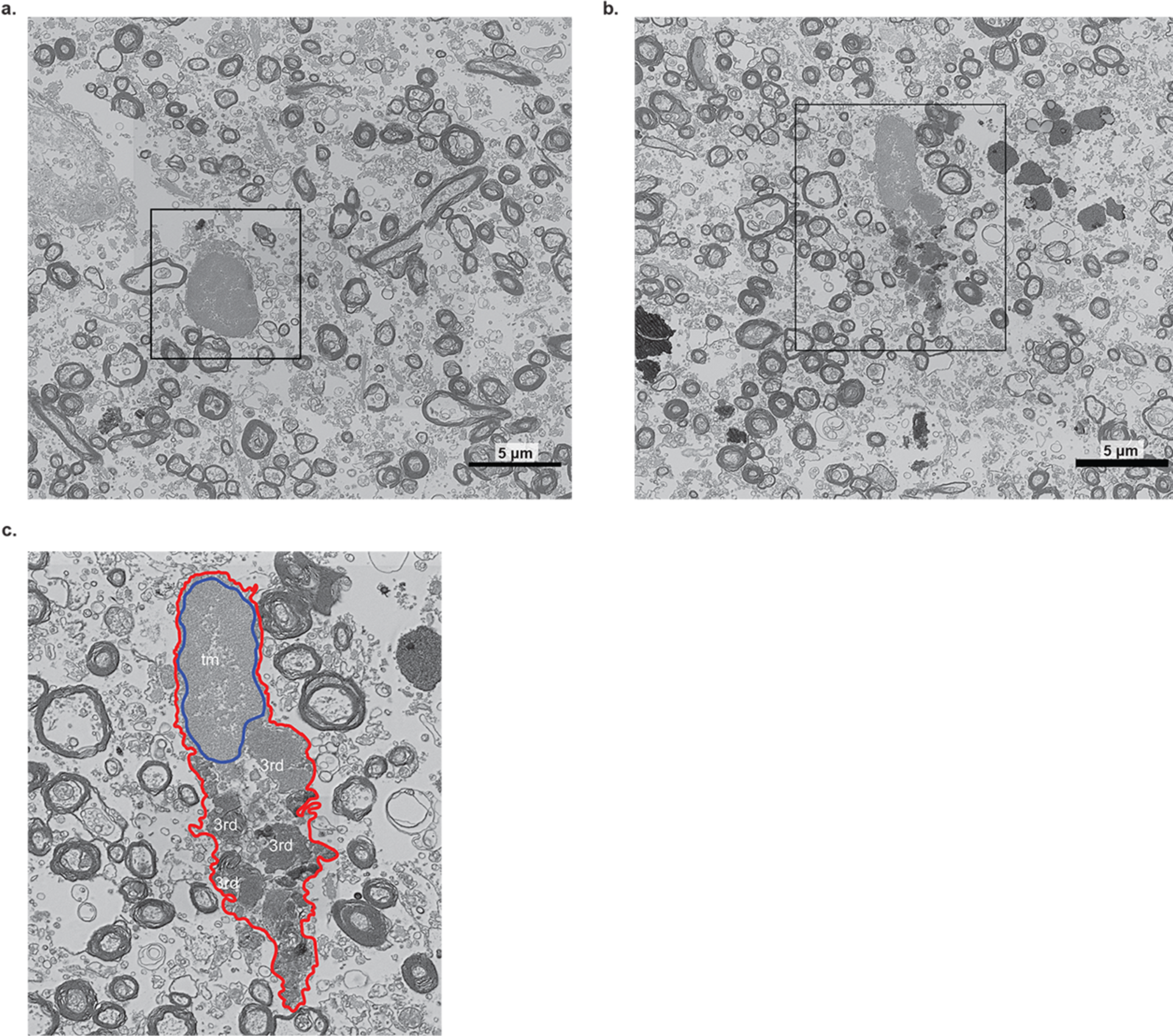
Dark microglia are present in the AD postmortem brain. Representative electron microscopy images of microglia from the hippocampus of a 79-year-old female AD patient. **(a)** The image shows dark microglia. A black square indicates a region of interest. The zoomed-in region of interest is shown in Fig. 1a. **(b)** The image shows typical microglia (tm). A black square indicates a region of interest. **(c)** The zoomed-in region of interest is indicated in (b). Red outline: cellular membrane. Blue outline: nuclear membrane. Tertiary lysosomes: 3rd (scale bars = 5 μm).

**Supplementary Figure 2.**
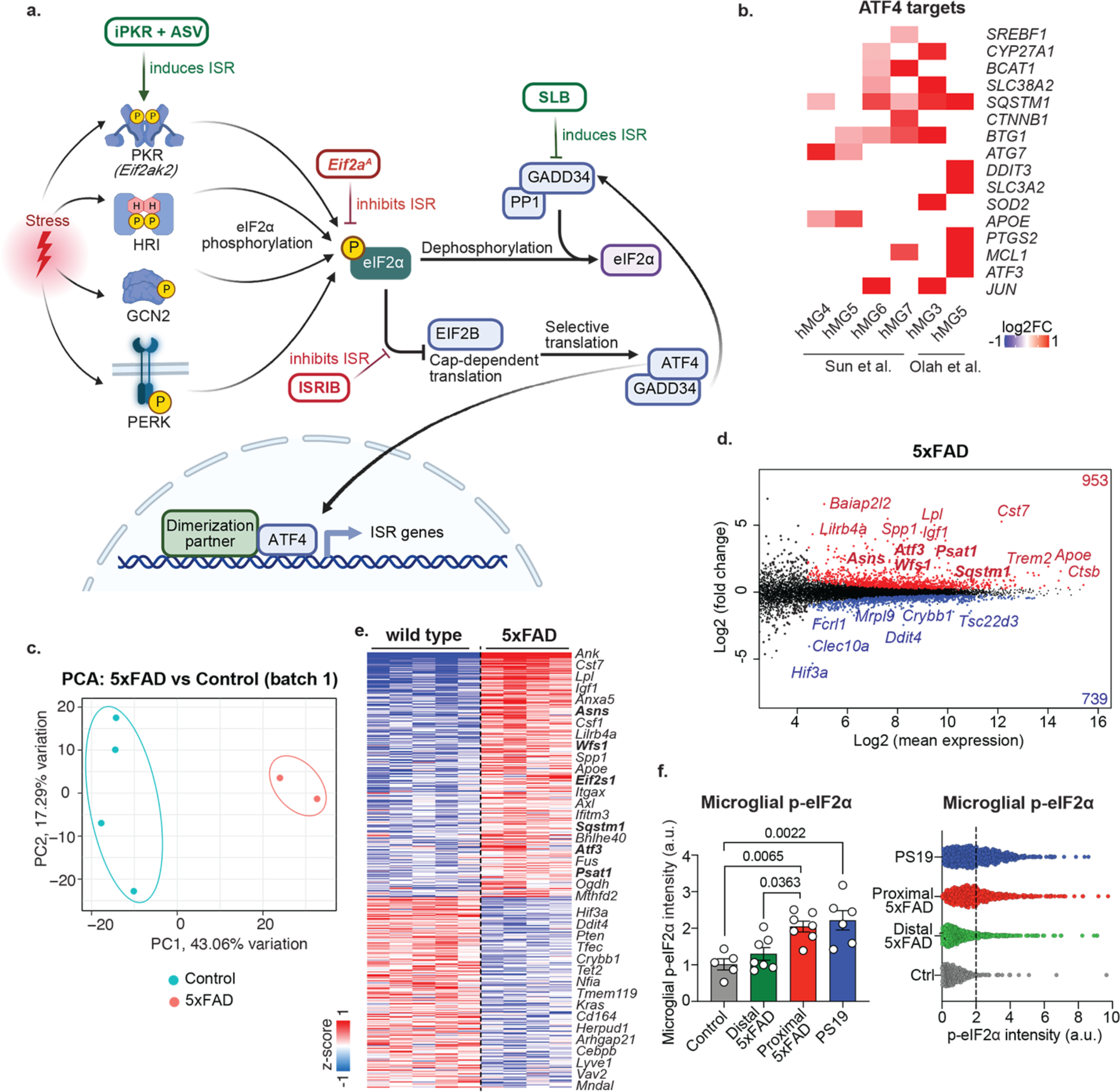
ISR is activated in microglia in AD mouse models and patients. **(a)** Schematic shows central events in integrated stress response (ISR). Adapted from “Stress Response Mechanism in Cells” by BioRender.com (2023). **(b)** Heatmap shows gene log2 (fold change) of ATF4 target genes in human microglia subsets identified by Sun et al.^36^ and Olah et al^35^. **(c)** Principle component analysis (PCA) of TRAP sequencing data from one batch of 6-month-old control (n=4) and 5xFAD samples (n=2). **(d)** The MA plot (representing log-ratio (M) on the *y*-axis and mean average (A) on the *x*-axis) shows differentially regulated genes (p value<0.05, baseMean>20 by DESeq2) in translating ribosome affinity purification (TRAP) sequencing of microglia from 6-month-old control (n=5) and 5xFAD (n=4) mice. **(e)** Heatmap shows z-scored variance stabilizing transformation (vst, gene expression score) of differentially regulated genes (p value<0.05, baseMean>20 by DESeq2). **(f)** The bar graph shows the mean p-eIF2α intensity in microglial cytosol (normalized to control) in the cortex of 6-month-old control (n=5), 6-month-old 5xFAD (n=7), and 8-month-old control PS19 mice (n=6). In 5xFAD mice, p-eIF2α intensity in microglia proximal to plaques (microglial surfaces less than 5 μm away from a plaque) and distal microglia (microglial surfaces more than 5 μm away from a plaque) are shown separately—one-way ANOVA with multiple comparisons. The bar graph with individual data points shows mean ± s.e.m. The dot plot shows the pooled mean cytosolic p-eIF2α intensity per microglia in the cortex of 6-month-old control (n=5), 6-month-old 5xFAD (n=7), and 8-month-old control PS19 mice (n=6), related to Fig. 1e. Dotted line represents the divide between p-eIF2α^high^ and p-eIF2α^low^ microglia.

**Supplementary Figure 3.**
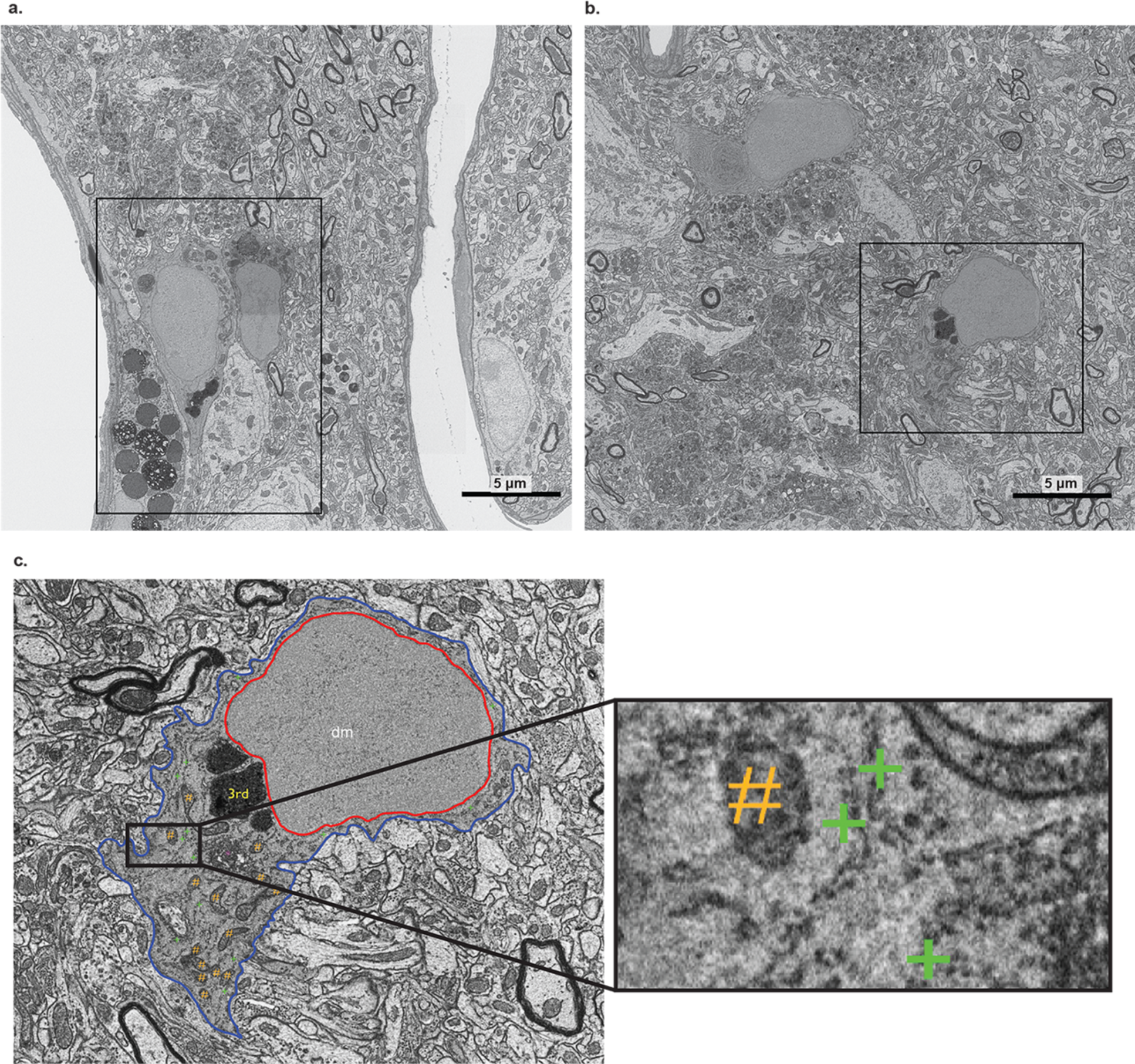
ISR is activated in microglia in the 5xFAD mouse model. **(a)** Two adjacent dark microglia close to a blood vessel contacting a p-eIF2α+ neuronal dendrite located near a blood vessel in a 5xFAD mouse. A black rectangle is shown to indicate the region of interest. The zoomed region of interest is shown in Fig 1h. **(b)** Two dark microglia in the isocortex of a 5xFAD mouse. A black rectangle is shown to indicate the region of interest. **(c)** The zoomed-in region of interest is indicated with a black rectangle in (b). The inset shows a zoomed-in view illustrating examples of p-eIF2α+ ER cisternae in dark microglia (green plus) and mitochondria (orange dash). The immunostaining is observed as a dotted pattern decorating the ER cisternae. dm: dark microglia, 3rd: tertiary lysosome. Red outline: cellular membrane. Blue outline: nuclear membrane. Purple asterisk: Golgi apparatus. (scale bars = 5 μm).

**Supplementary Figure 4.**
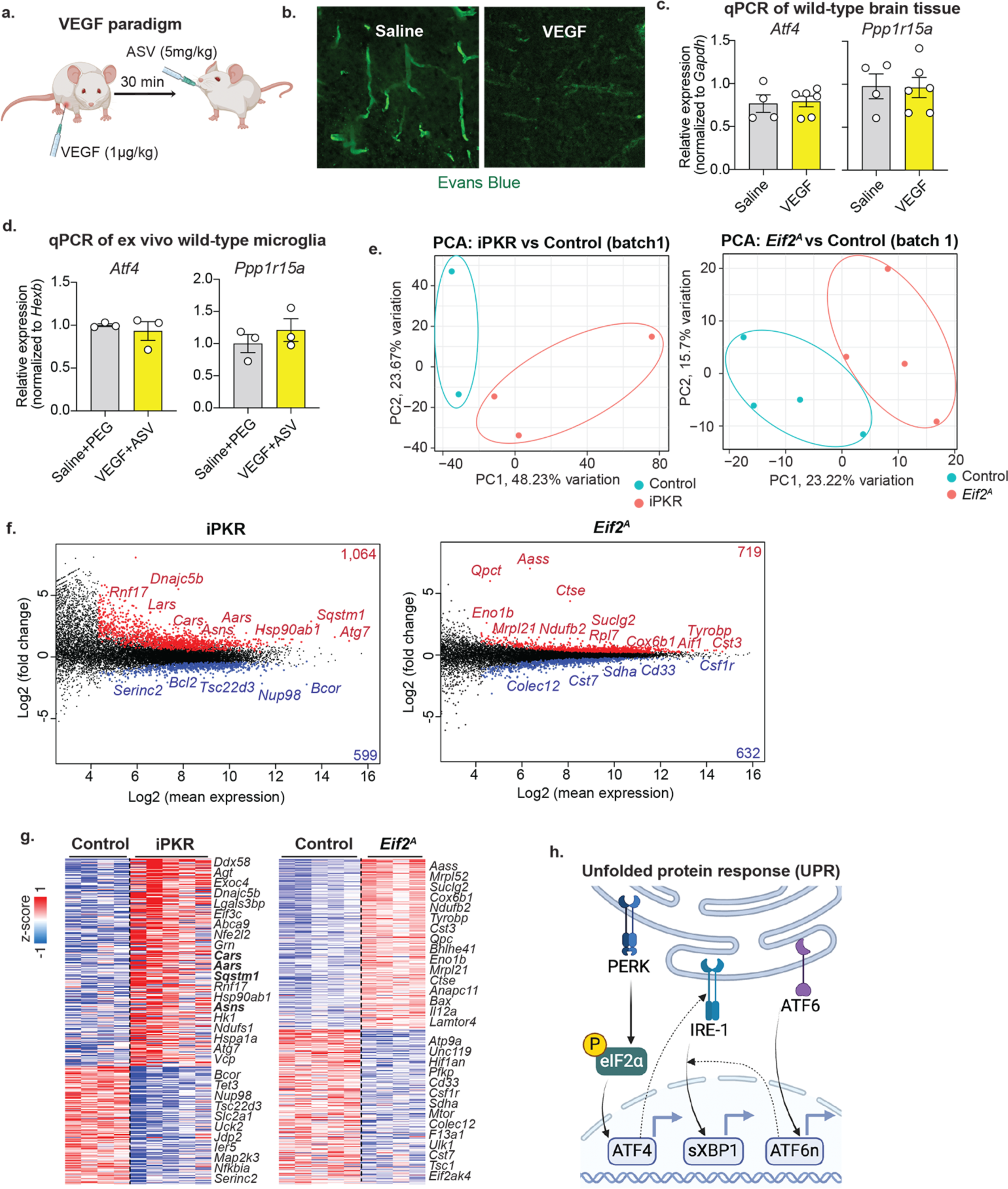
Validation and characterization of the microglia-specific chemogenetic ISR model. **(a)** Schematic shows the treatment paradigms to deliver ASV. **(b)** Evans Blue (green) in a mouse intraperitoneally injected with vehicle PBS:Saline or 1 μg/kg VEGF. **(c)** The bar graphs show qPCR-based measurement levels of selected mRNAs in the brains of mice treated as in (a). Brain tissue was isolated 2.5 hours after PEG (n=4) or ASV (n=6) oral administration—unpaired two-tailed t-test. **(d)** The bar graphs show qPCR-based measurement levels of selected mRNAs in microglia isolated from mice treated as in (a). Microglia were isolated 2.5 hours after PEG or ASV oral administration (n=3/group)—unpaired two-tailed t-test. Bar graphs with individual data points show mean ± s.e.m. **(e)** Principle component analysis (PCA) of TRAP sequencing data from one batch of 6-month-old control (vehicle-treated iPKR, n=2) and ASV-treated iPKR samples (n=3), and one batch of control TRAP (n=4) and *Eif2^A^* TRAP mice (n=4). **(f)** The MA plot (representing log-ratio (M) on the *y*-axis and mean average (A) on the *x*-axis) shows differentially regulated genes (p value<0.05, baseMean>20 by DESeq2) in TRAP sequencing of microglia from left: 6-month-old (vehicle-treated iPKR, n=4) and ASV-treated iPKR samples (n=5) and right: 6-month-old control TRAP (n=5) and *Eif2^A^* TRAP mice (n=4). **(g)** Heatmap shows z-scored variance stabilizing transformation (vst, gene expression score) of differentially regulated genes (p value<0.05, baseMean>20 by DESeq2). **(h)** Schematic shows unfolded protein pathways and their crosstalk. Created with BioRender.com (2023).

**Supplementary Figure 5.**
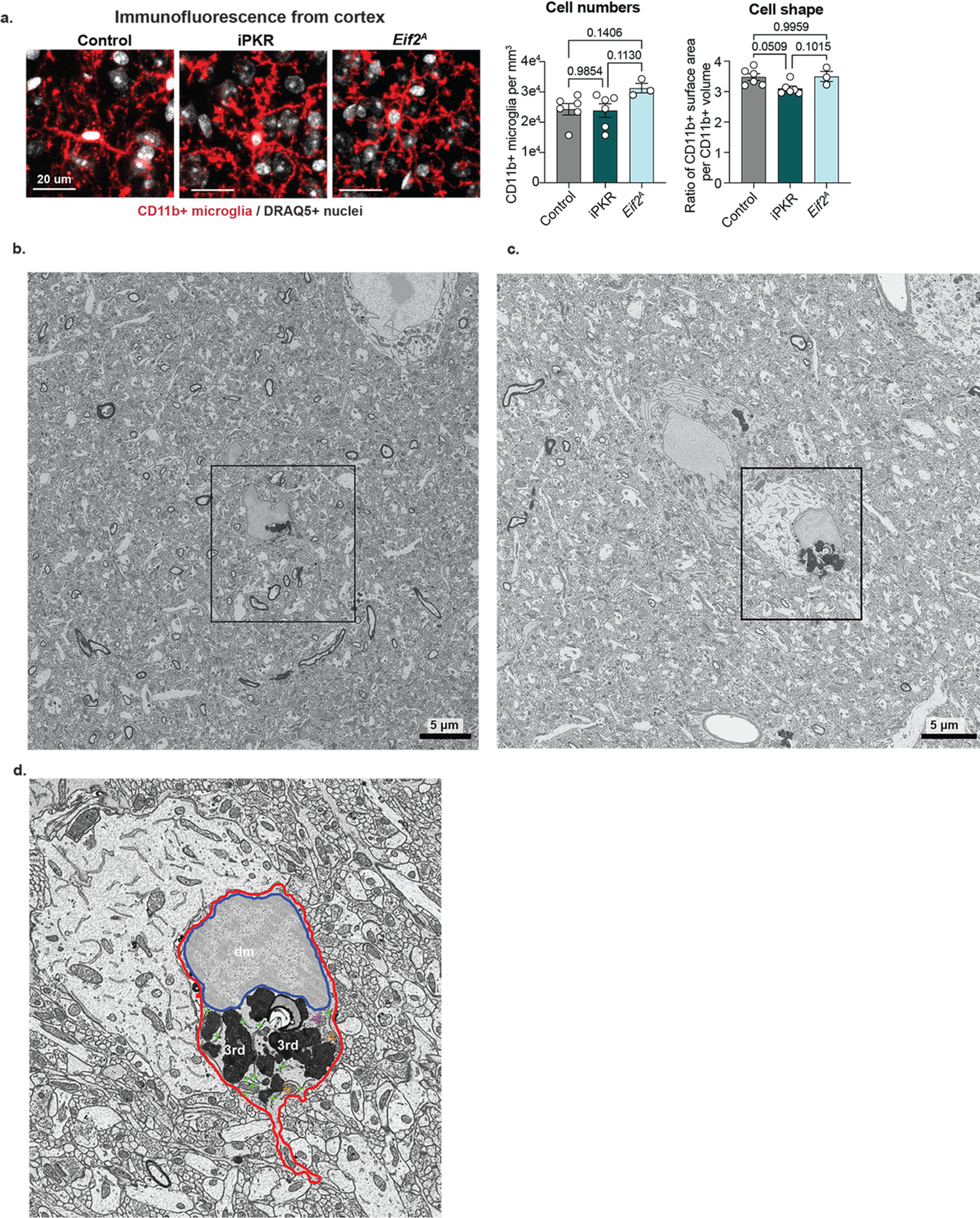
Activation of ISR autonomously generates features of dark microglia. **(a)** Representative immunofluorescence images (left) and quantifications (right). CD11b+ microglia: red, DRAQ5+ nuclei: gray. The first bar graph shows cell numbers (number of microglial nuclei), and the second graph shows the area-to-volume ratio of microglia in the cortex of 6-month-old ASV-treated control (n=6), ASV-treated iPKR (n=6), and *Eif2^A^* mice (n=3)—one-way ANOVA with multiple comparisons. Bar graphs with individual data points show mean ± s.e.m. **(b, c)** A dark microglia with intermediate phenotype in a female iPKR mouse (scale bars = 5 μm). A black square indicates a region of interest. **(d)** The zoomed-in region of interest, as indicated in (c), shows dark microglia (dm) containing several tertiary lysosomes (3rd). Red outline: cellular membrane. Blue outline: nuclear membrane. Green plus: ER. Orange hash: mitochondria. Purple asterisk: Golgi apparatus.

**Supplementary Figure 6.**
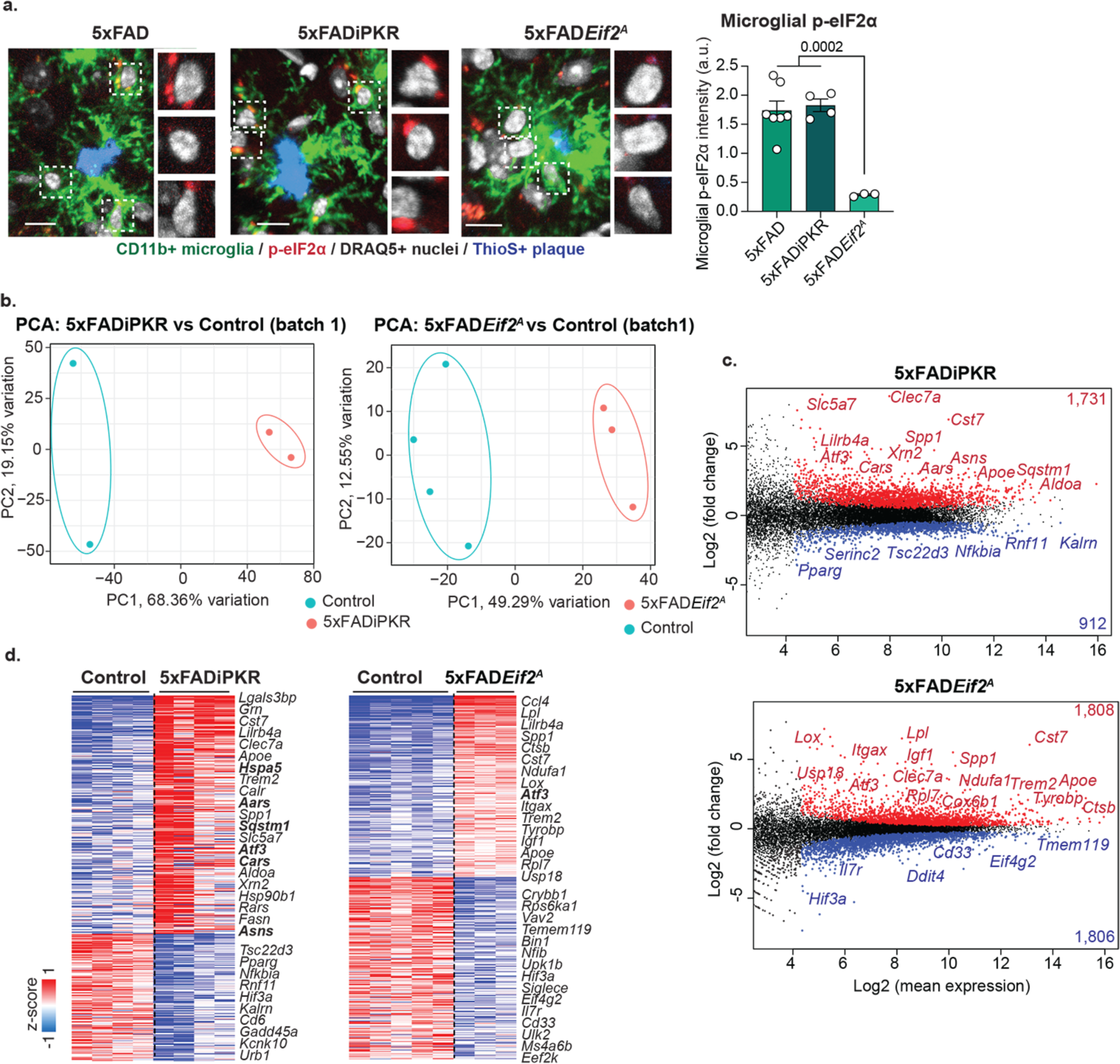
Modulation of ISR induces distinct gene expression profiles by microglia-specific TRAP sequencing. **(a)** Representative immunofluorescence images (left) and quantifications (right). p-eIF2α: red, CD11b+ microglia: green, ThioS+ dense core plaques: blue, DRAQ5+ nuclei: gray. The bar graph shows the mean p-eIF2α intensity around microglial nuclei (inset) in the cortex of 6-month-old ASV-treated control 5xFAD (n=7), ASV-treated 5xFADiPKR (n=4), and 5xFAD*Eif2^A^* mice (n=3)—one-way ANOVA with multiple comparisons. **(b)** Principle component analysis (PCA) of TRAP sequencing data from one batch of 6-month-old (vehicle-treated iPKR, n=2) and ASV-treated 5xFADiPKR samples (n=2) and one batch of 6-month-old control TRAP (n=4) and 5xFAD*Eif2^A^* TRAP mice (n=3). **(c)** The MA plot (representing log-ratio (M) on the *y*-axis and mean average (A) on the *x*-axis) shows differentially regulated genes (p value<0.05, baseMean>20 by DESeq2) in TRAP sequencing of microglia from 6-month-old (vehicle-treated iPKR, n=4) and ASV-treated iPKR samples (n=4) and control TRAP (n=5) and 5xFAD*Eif2^A^* TRAP mice (n=3). **(d)** Heatmap shows z-scored variance stabilizing transformation (vst, gene expression score) of differentially regulated genes (p value<0.05, baseMean>20 by DESeq2).

**Supplementary Figure 7.**
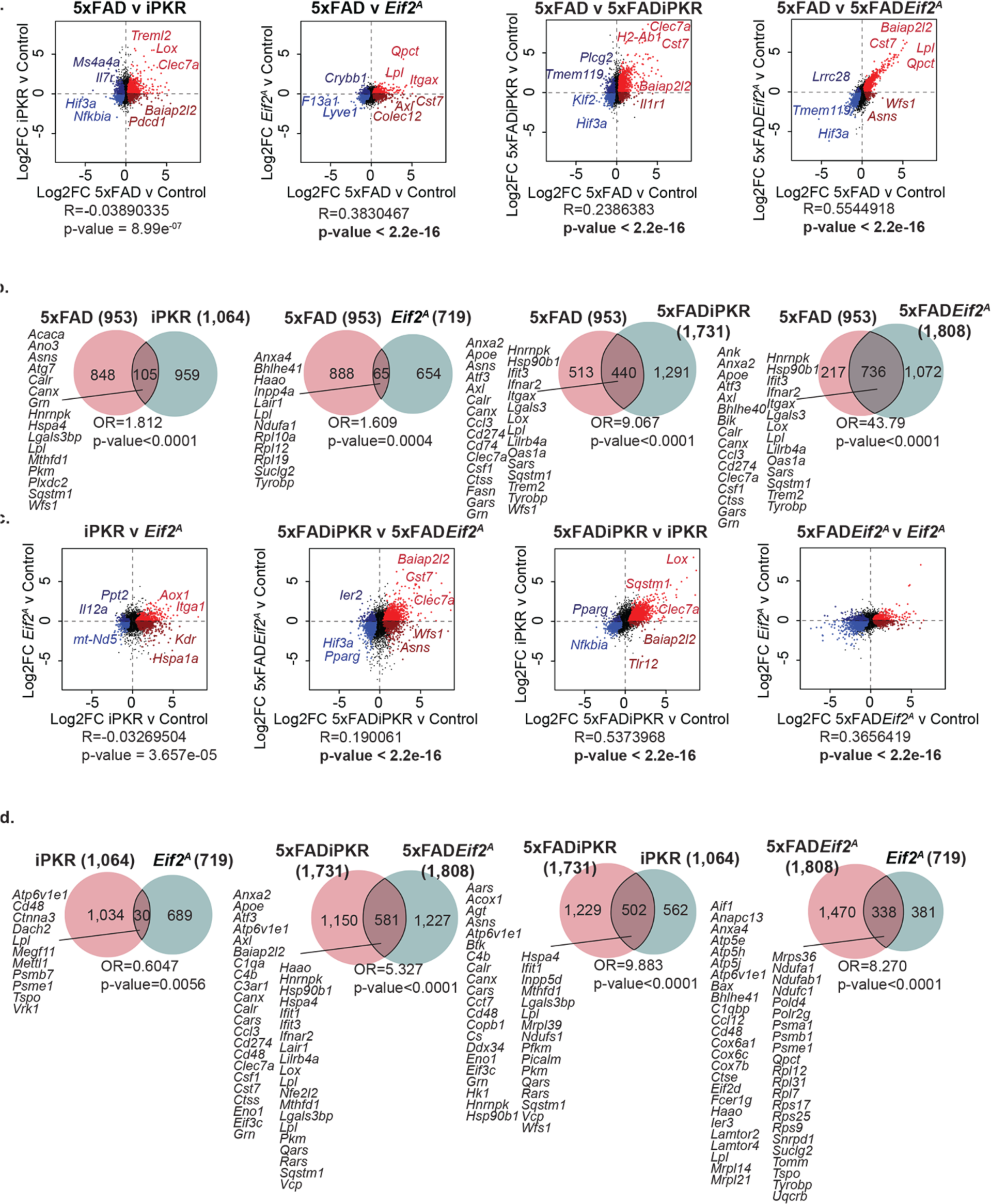
Comparisons of ISR-dependent gene expression changes by microglia-specific TRAP sequencing. **(a, c)** The scatter plots compare the log2 (fold change) between different differentially enriched genes identified by TRAP. Genes significantly upregulated in both groups: red. Genes upregulated in the x-axis group only: dark red. Genes downregulated in both groups: blue. Genes downregulated in the x-axis group only: dark blue. Pearson correlation was used. **(b, d)** Venn diagrams show overlap between upregulated genes and all other genesets. χ^2^ test was used.

**Supplementary Figure 8.**
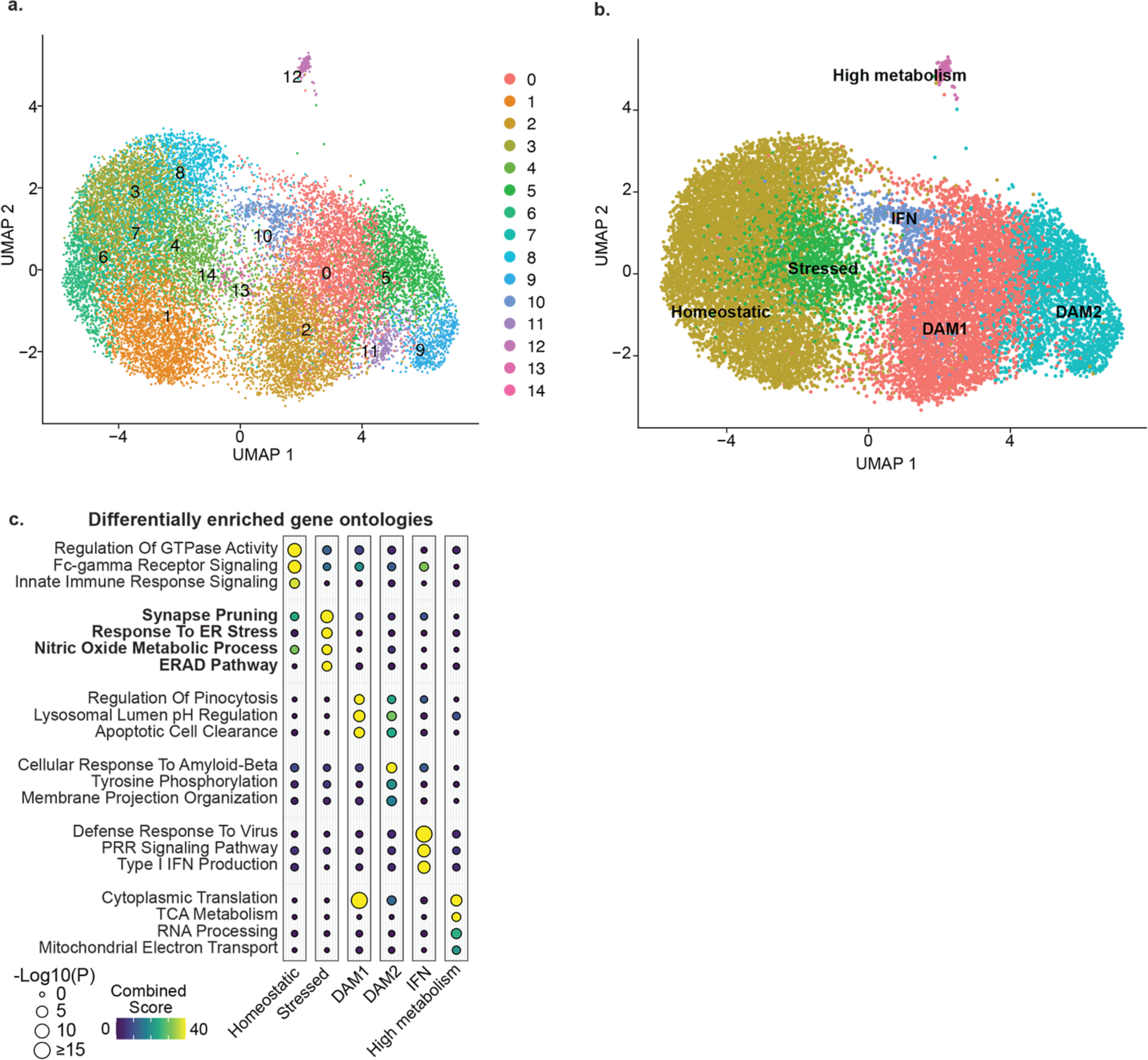
Analysis of microglia-specific single nuclei sequencing. **(A-b)** Uniform manifold approximation and projection (UMAP) visualizations from the Chromium 10X single nuclei RNA-sequencing of cortical microglial nuclei from 5xFAD, 5xFADiPKR, and 5xFAD*Eif2^A^* mice show automatically generated clusters (a) and clusters organized into functional groups (b). **(c)** The balloon plot represents the p-value (size) and combined score (color) for gene ontologies of cluster markers in microglia-specific single nuclei sequencing.

**Supplementary Figure 9.**
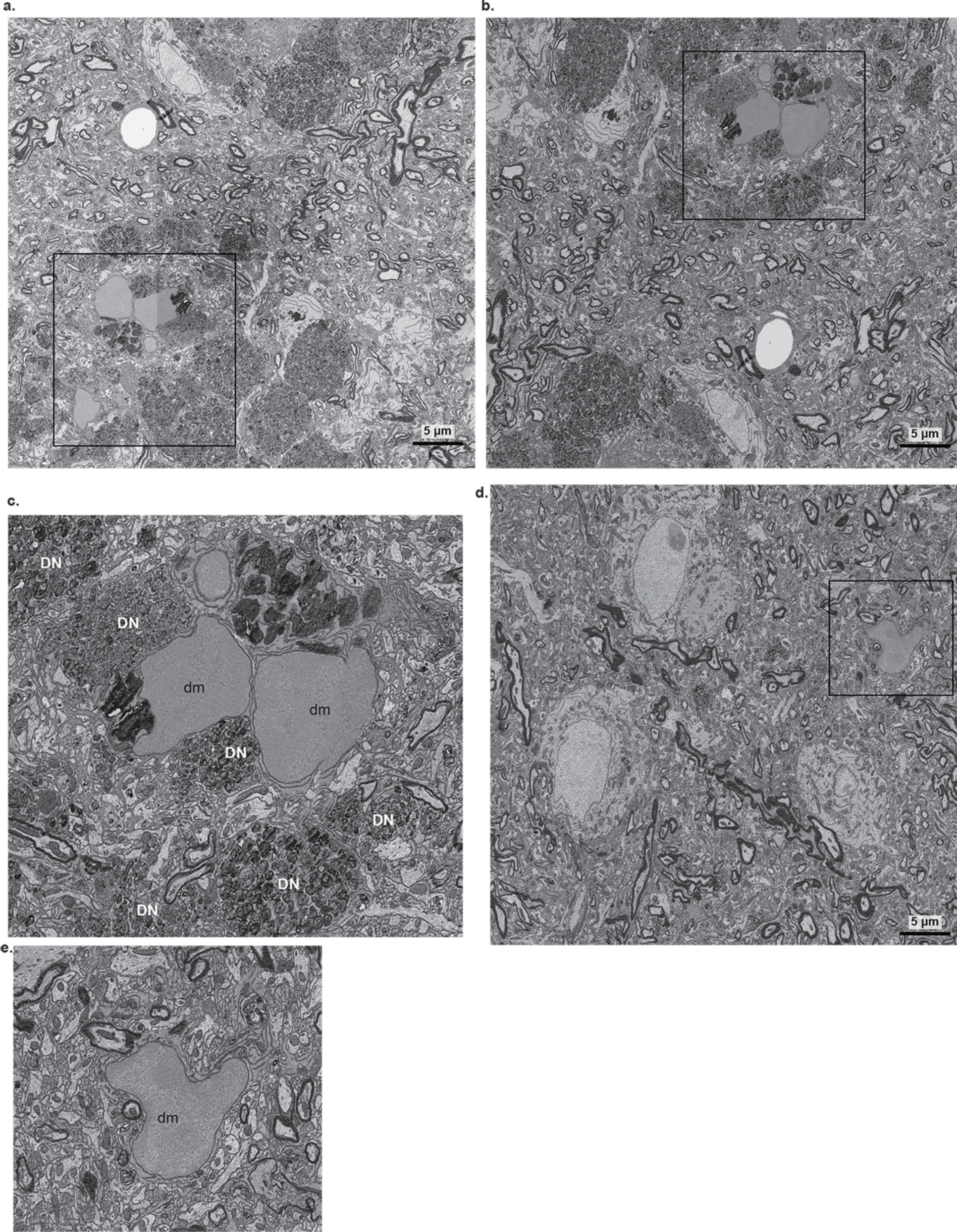
Dark microglia in the 5xFADiPKR model. **(a, b, d)** Dark microglia (dm) were observed in the isocortex of a 6-month-old male 5xFADiPKR mouse. The black rectangle indicates a region of interest that is shown zoomed in in Fig. 3g for (a), in (c) for (b), and in (e) for (d). Scale bars = 5 μm. **(c, e)** Examples of dark microglia (dm) near dystrophic neurites (DN, c) or distal to dystrophic neurites (e).

**Supplementary Figure 10.**
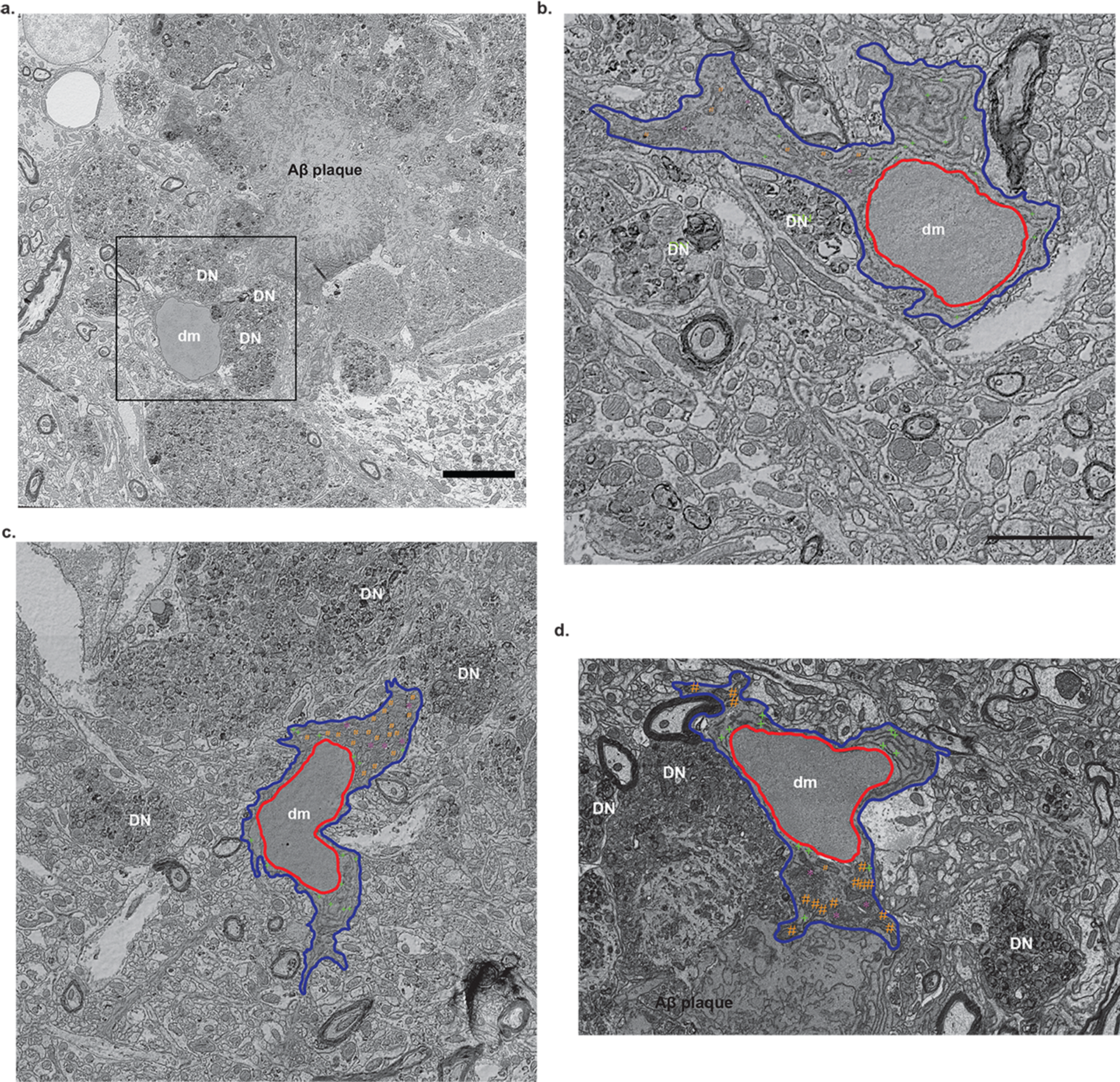
Dark microglia in the 5xFAD*Eif2^A^* model. **(a)** Dark microglia (dm) in the isocortex of a 6-month-old female 5xFAD*Eif2^A^* mouse that juxtaposes an Aβ plaque and dystrophic neurites (DN). The black rectangle denotes the region of interest shown zoomed in in Fig. 3g. **(b-d)** Examples of dark microglia in the isocortex of a 6-month-old female 5xFAD*Eif2^A^* mouse. 3rd: tertiary lysosome. Blue outline: cellular membrane. Red outline: nuclear membrane. Green plus: ER. Orange hashtag: mitochondria. Purple star: Golgi apparatus (scale bars = 5 μm).

**Supplementary Figure 11.**
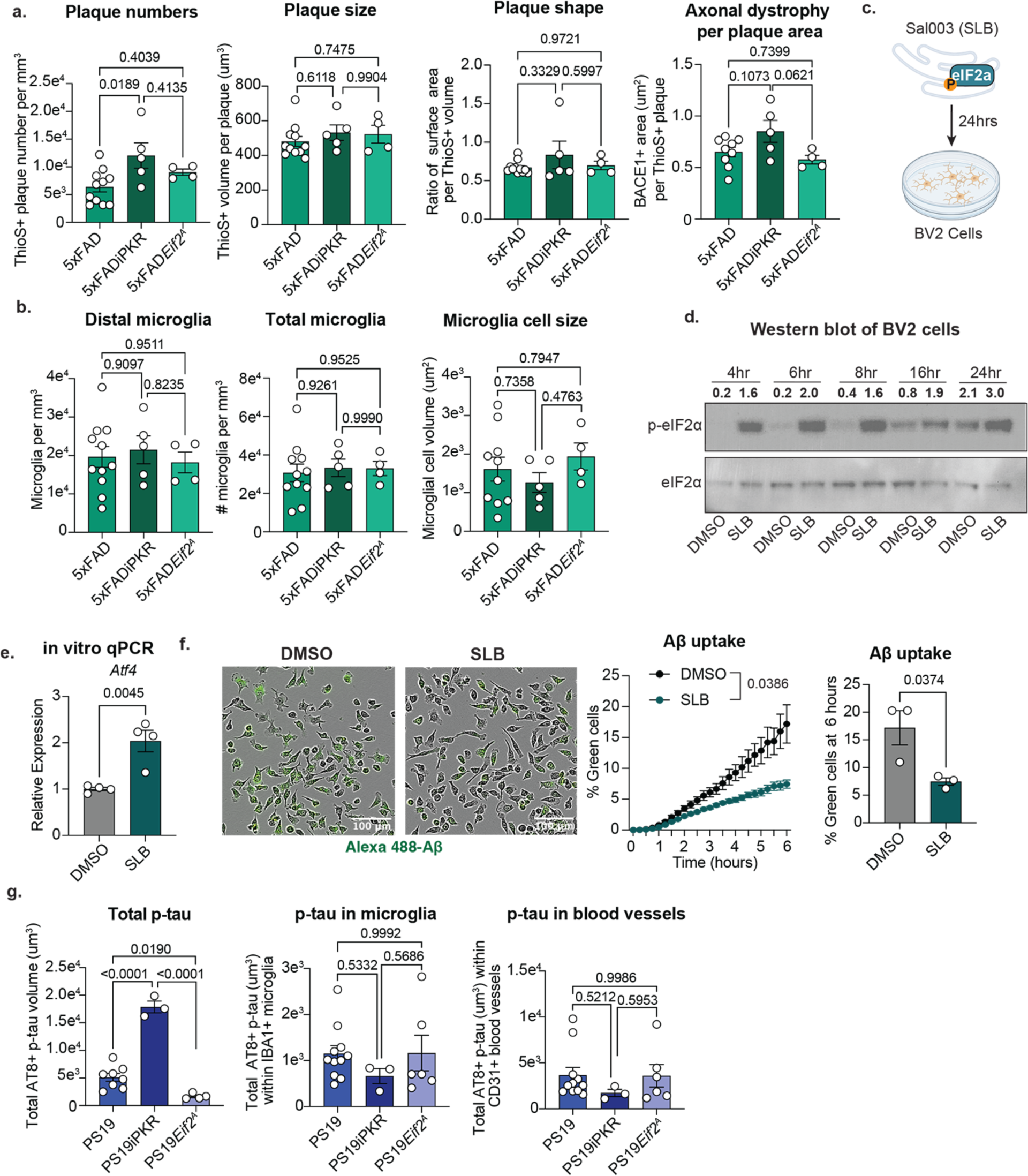
ISR has a strong impact on microglial response to amyloid and tau pathology. **(a)** Bar graphs show total ThioS+ plaque numbers, ThioS+ plaque volume per plaque, ThioS+ plaque surface area per volume, and BACE1+ axonal dystrophy volume normalized to ThioS+ plaque volume in the cortex of 6-month-old 5xFAD (n=11, n=9 for BACE1+axonal dystrophy), 5xFADiPKR (n=5), and 5xFAD*Eif2^A^* mice (n=4) quantified from Fig. 4a. **(b)** Bar graphs show CD11b+ distal microglia (nuclei of microglia more than 5 μm away from a plaque) per plaque, the total number of CD11b+ microglia, and CD11b+ surface area per volume in the cortex of 6-month-old 5xFAD (n=11), 5xFADiPKR (n=5), and 5xFAD*Eif2^A^* mice (n=4) quantified from Fig. 4a. **(c)** Schematic shows induction of ISR in BV2 cells by SLB. **(d)** Representative Western blots of p-eIF2α and total eIF2α (t-eIF2α) in BV2 microglia stimulated with SLB at different time points. The quantification shows the ratio of area under the curve for p-/t eIF2α. **(e)** The bar graph shows the qPCR-based measurement of *Atf4* in BV2 cells stimulated with SLB for 24 hours (n=4 wells/condition)—unpaired two-tailed t-test. **(f)** Representative composite brightfield and green fluorescence images (left) and quantifications (right). BV2 microglia cells in brightfield and Alexa-488-labeled Aβ42: green. Cells that uptake Aβ42 appear bright green. The line graph shows the % green cells in DMSO and SLB-treated cells. The bar graph shows the % green cells in DMSO and SLB-treated cells after 6 hours of treatment (n=3 wells/condition)—unpaired two-tailed t-test. **(g)** The first bar graph shows the total volume of AT8+ phospho-Tau in the imaged region in the cortex of 8-month-old ASV-treated PS19 (n=8), PS19iPKR (n=3), and PS19*Eif2^A^* mice (n=4). The next bar graphs show the volume of AT8+ phospho-Tau in IBA1+ microglia and CD31+ blood vessels in the cortex of 8-month-old ASV-treated PS19 (n=11), PS19iPKR (n=3), and PS19*Eif2^A^* mice (n=6). Bar graphs with individual data points show mean ± s.e.m. One-way ANOVA with multiple comparisons.

**Supplementary Figure 12.**
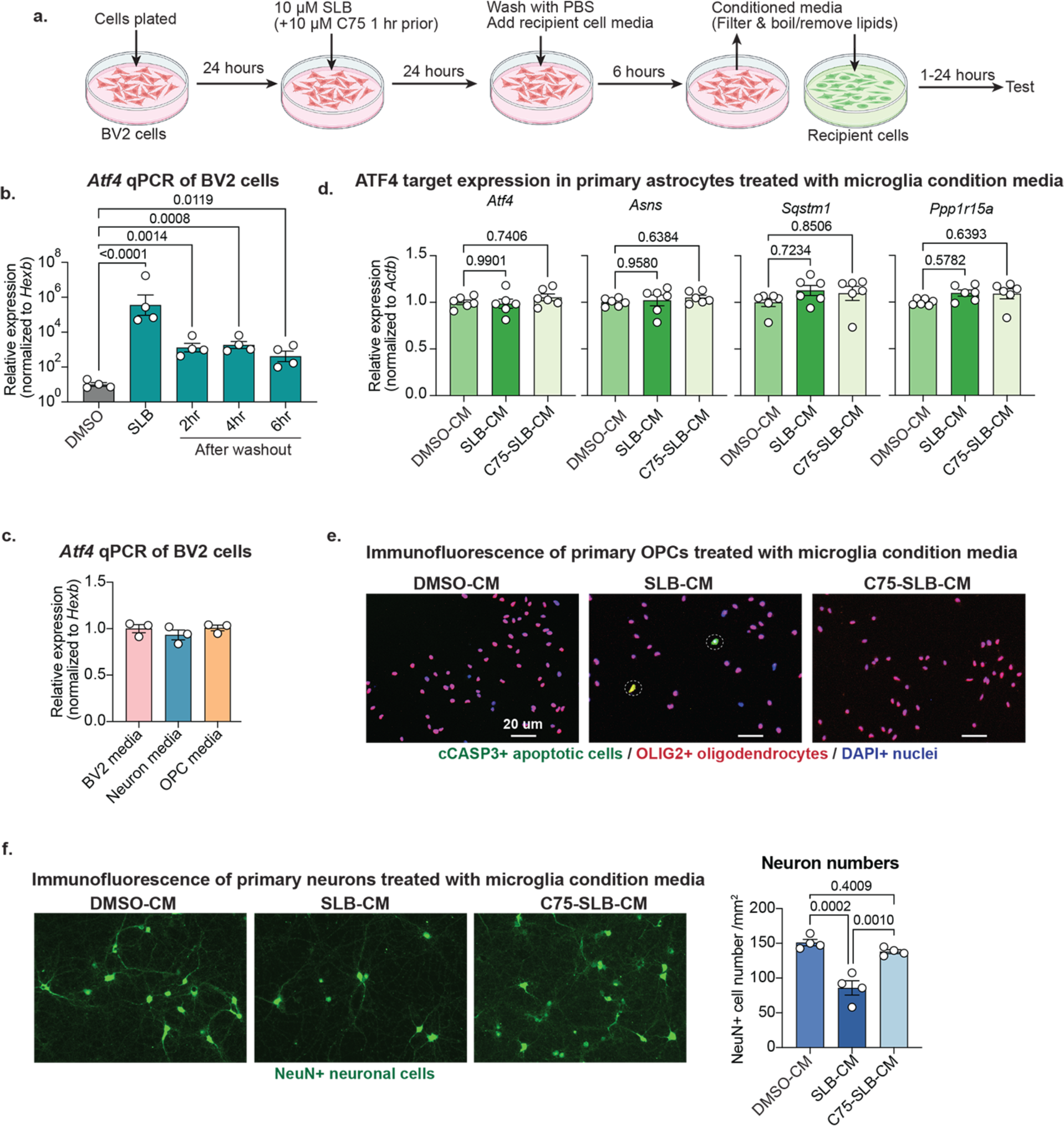
ISR-induced lipid secretion has a detrimental impact on other brain cell types. **(a)** Schematic shows the experimental design of conditioned media experiments. **(b)** The bar graph shows the qPCR-based measurement of *Atf4* in BV2 cells treated as in (a) at different time points after washing out SLB (n=4 wells/ condition)—one-way ANOVA with multiple comparisons. **(c)** The bar graph shows the qPCR-based measurement of *Atf4* in BV2 microglia after adding different recipient cell media (n=3 wells/condition)—one-way ANOVA with multiple comparisons. **(d)** The bar graphs show the qPCR-based measurement of *Atf4* and ATF4 target genes in recipient primary astrocytes treated with SLB-conditioned media from BV2 cells as in (a) (n=6 wells/condition)— one-way ANOVA with multiple comparisons. **(e)** Representative immunofluorescence images show OLIG2+ oligodendrocyte lineage cells: red, cCASP3+ apoptotic cells: green, DAPI+ nuclei: blue. **(f)** Representative immunofluorescence images (left) and quantification (right). NeuN+ neuronal cells: green. The bar graph shows the number of NeuN+ cells per mm^2^ (n=4 wells/group). Bar graphs with individual data points show mean ± s.e.m. One-way ANOVA with multiple comparisons.

**Supplementary Figure 13.**
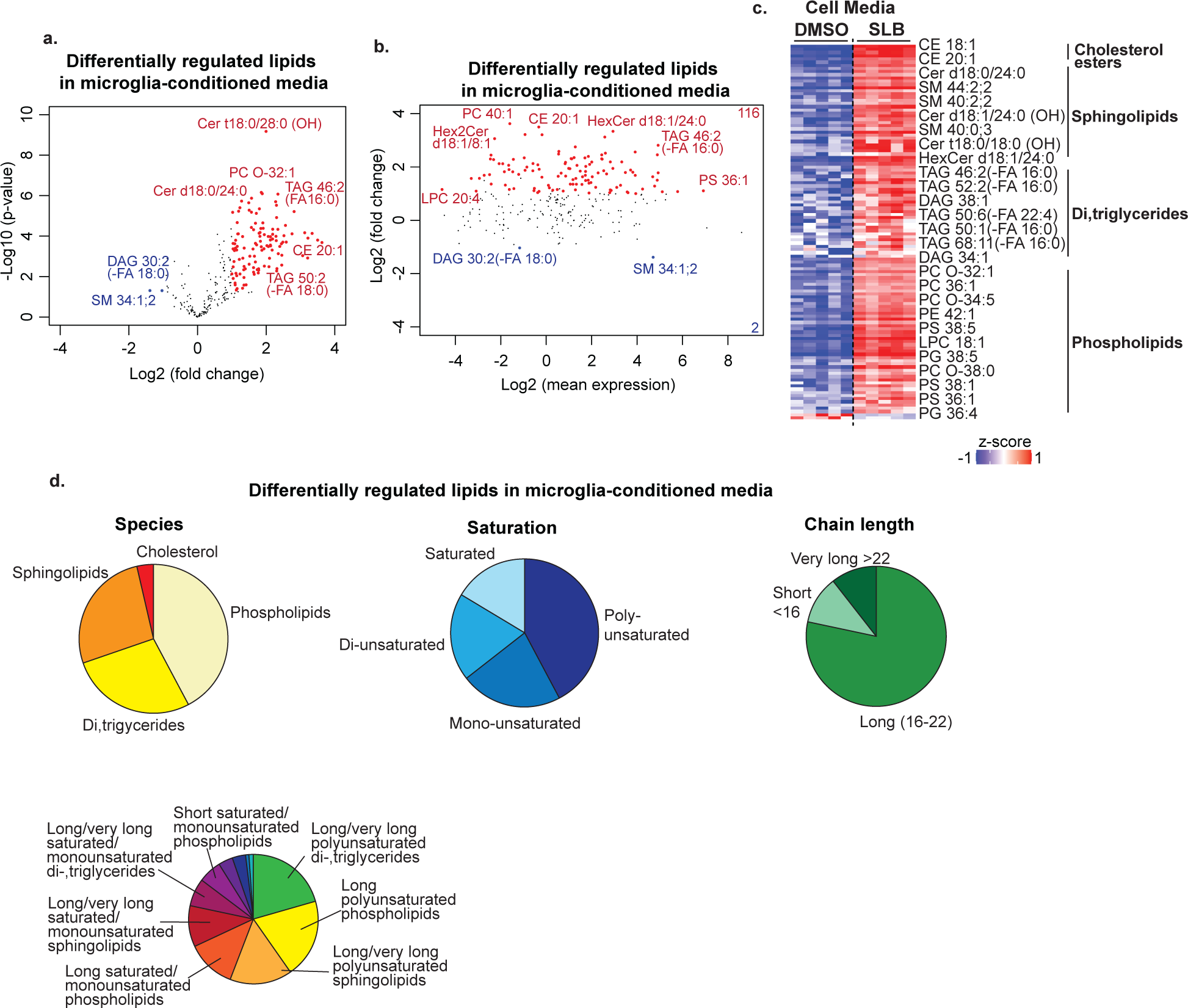
ISR induces the secretion of diverse long-chain lipid species. **(a, b)** Volcano plot (a) and MA plot (b) show differentially regulated (p-value <0.05, fold change > 2 by t-test) lipid species detected by untargeted lipidomics in BV2 cell media treated as in Fig. S12a (n=5/group). Upregulated lipids are in red, and downregulated lipids are in blue. **(c)** Heatmap shows z-scored relative amounts of differentially regulated lipids in cell media from SLB-treated microglia as in Supplementary Fig 12a. **(d)** Pie charts classify significantly upregulated lipids in (a-c).

## Bibliography

1 Efthymiou, A. G. & Goate, A. M. Late onset Alzheimer’s disease genetics implicates microglial pathways in disease risk. Molecular neurodegeneration 12, 43–43, doi:10.1186/s13024-017-0184-x (2017).

2 Nott, A. et al. Brain cell type–specific enhancer–promoter interactome maps and disease <strong>-</strong>risk association. Science 366, 1134–1139, doi:10.1126/science.aay0793 (2019).

3 Masuda, T., Sankowski, R., Staszewski, O. & Prinz, M. Microglia Heterogeneity in the Single-Cell Era. Cell Reports 30, 1271–1281, 10.1016/j.celrep.2020.01.010 (2020).

4 Chen, Y. & Colonna, M. Microglia in Alzheimer’s disease at single-cell level. Are there common patterns in humans and mice? Journal of Experimental Medicine 218, doi:10.1084/jem.20202717 (2021).

5 Keren-Shaul, H. et al. A Unique Microglia Type Associated with Restricting Development of Alzheimer’s Disease. Cell 169, 1276–1290.e1217, 10.1016/j.cell.2017.05.018 (2017).

6 Krasemann, S. et al. The TREM2-APOE Pathway Drives the Transcriptional Phenotype of Dysfunctional Microglia in Neurodegenerative Diseases. Immunity 47, 566–581.e569, 10.1016/j.immuni.2017.08.008 (2017).

7 Deczkowska, A. et al. Disease-Associated Microglia: A Universal Immune Sensor of Neurodegeneration. Cell 173, 1073–1081, 10.1016/j.cell.2018.05.003 (2018).

8 Azevedo, E. P. et al. Activated microglia mediate synapse loss and short-term memory deficits in a mouse model of transthyretin-related oculoleptomeningeal amyloidosis. Cell Death & Disease 4, e789–e789, doi:10.1038/cddis.2013.325 (2013).

9 Hong, S. et al. Complement and microglia mediate early synapse loss in Alzheimer mouse models. Science, doi:10.1126/science.aad8373 (2016).

10 Lui, H. et al. Progranulin Deficiency Promotes Circuit-Specific Synaptic Pruning by Microglia via Complement Activation. Cell 165, 921–935, doi:10.1016/j.cell.2016.04.001 (2016).

11 Asai, H. et al. Depletion of microglia and inhibition of exosome synthesis halt tau propagation. Nature Neuroscience 18, 1584–1593, doi:10.1038/nn.4132 (2015).

12 Hopp, S. C. et al. The role of microglia in processing and spreading of bioactive tau seeds in Alzheimer’s disease. Journal of Neuroinflammation 15, 269, doi:10.1186/s12974-018-1309-z (2018).

13 Felsky, D. et al. Neuropathological correlates and genetic architecture of microglial activation in elderly human brain. Nature Communications 10, 409, doi:10.1038/s41467-018-08279-3 (2019).

14 Gibson, E. M. et al. Methotrexate Chemotherapy Induces Persistent Tri-glial Dysregulation that Underlies Chemotherapy-Related Cognitive Impairment. Cell 176, 43–55.e13, doi:10.1016/j.cell.2018.10.049 (2019).

15 Guttenplan, K. A. et al. Neurotoxic reactive astrocytes induce cell death via saturated lipids. Nature 599, 102–107, doi:10.1038/s41586-021-03960-y (2021).

16 Fernández-Castañeda, A. et al. Mild respiratory COVID can cause multi-lineage neural cell and myelin dysregulation. Cell 185, 2452–2468.e2416, 10.1016/j.cell.2022.06.008 (2022).

17 Bisht, K. et al. Dark microglia: A new phenotype predominantly associated with pathological states. Glia 64, 826–839, doi:10.1002/glia.22966 (2016).

18 El Hajj, H., et al. Ultrastructural evidence of microglial heterogeneity in Alzheimer’s disease amyloid pathology. Journal of Neuroinflammation 16, 87, doi:10.1186/s12974-019-1473-9 (2019).

19 St-Pierre, M.-K. et al. Ultrastructural characterization of dark microglia during aging in a mouse model of Alzheimer’s disease pathology and in human post-mortem brain samples. Journal of Neuroinflammation 19, 235, doi:10.11 (2022).

20 Bond, S., Lopez-Lloreda, C., Gannon, P. J., Akay-Espinoza, C. & Jordan-Sciutto, K. L. The Integrated Stress Response and Phosphorylated Eukaryotic Initiation Factor 2α in Neurodegeneration. Journal of Neuropathology & Experimental Neurology 79, 123–143, doi:10.1093/jnen/nlz129 (2020).

21 Costa-Mattioli, M. & Walter, P. The integrated stress response: From mechanism to disease. Science 368, doi:10.1126/science.aat5314 (2020).

22 Chang, R. C., Wong, A. K., Ng, H. K. & Hugon, J. Phosphorylation of eukaryotic initiation factor-2alpha (eIF2alpha) is associated with neuronal degeneration in Alzheimer’s disease. Neuroreport 13, 2429–2432, doi:10.1097/00001756-200212200-00011 (2002).

23 Paccalin, M. et al. Activated mTOR and PKR kinases in lymphocytes correlate with memory and cognitive decline in Alzheimer’s disease. Dement Geriatr Cogn Disord 22, 320–326, doi:10.1159/000095562 (2006).

24 Di Battista, A. M., Heinsinger, N. M. & Rebeck, G. W. Alzheimer’s Disease Genetic Risk Factor APOE-ε4 Also Affects Normal Brain Function. Curr Alzheimer Res 13, 1200–1207, doi:10.2174/1567205013666160401115127 (2016).

25 Machlovi, S. I. et al. APOE4 confers transcriptomic and functional alterations to primary mouse microglia. Neurobiology of Disease 164, 105615, 10.1016/j.nbd.2022.105615 (2022).

26 Millet, A., Ledo, J. H. & Tavazoie, S. F. An exhausted-like microglial population accumulates in aged and APOE4 genotype Alzheimer’s brains. Immunity 57, 153–170.e156, 10.1016/j.immuni.2023.12.001 (2024).

27 Novikova, G. et al. Integration of Alzheimer’s disease genetics and myeloid genomics identifies disease risk regulatory elements and genes. Nat Commun 12, 1610, doi:10.1038/s41467-021-21823-y (2021).

28 Chang, R. C. C., Wong, A. K. Y., Ng, H.-K. & Hugon, J. Phosphorylation of eukaryotic initiation factor-2α (eIF2α) is associated with neuronal degeneration in Alzheimer’s disease. NeuroReport 13 (2002).

29 Page, G. et al. Activated double-stranded RNA-dependent protein kinase and neuronal death in models of Alzheimer’s disease. Neuroscience 139, 1343–1354, 10.1016/j.neuroscience.2006.01.047 (2006).

30 Kim, H.-S. et al. Swedish amyloid precursor protein mutation increases phosphorylation of eIF2α in vitro and in vivo. Journal of Neuroscience Research 85, 1528–1537, 10.1002/jnr.21267 (2007).

31 Oliveira Mauricio, M., et al. Correction of eIF2-dependent defects in brain protein synthesis, synaptic plasticity, and memory in mouse models of Alzheimer’s disease. Sci Signal 14, eabc5429, doi:10.1126/scisignal.abc5429 (2021).

32 Pakos-Zebrucka, K. et al. The integrated stress response. EMBO Rep 17, 1374–1395, doi:10.15252/embr.201642195 (2016).

33 Neill, G. & Masson, G. R. A stay of execution: ATF4 regulation and potential outcomes for the integrated stress response. Front Mol Neurosci 16, 1112253, doi:10.3389/fnmol.2023.1112253 (2023).

34 Lau, S.-F., Cao, H., Fu, A. K. Y. & Ip, N. Y. Single-nucleus transcriptome analysis reveals dysregulation of angiogenic endothelial cells and neuroprotective glia in Alzheimer’s disease. Proceedings of the National Academy of Sciences 117, 25800–25809, doi:10.1073/pnas.2008762117 (2020).

35 Olah, M. et al. Single cell RNA sequencing of human microglia uncovers a subset associated with Alzheimer’s disease. Nature Communications 11, 6129, doi:10.1038/s41467-020-19737-2 (2020).

36 Sun, N. et al. Human microglial state dynamics in Alzheimer’s disease progression. Cell 186, 4386–4403.e4329, doi:10.1016/j.cell.2023.08.037 (2023).

37 Krämer, A., Green, J., Pollard, J., Jr. & Tugendreich, S. Causal analysis approaches in Ingenuity Pathway Analysis. Bioinformatics 30, 523–530, doi:10.1093/bioinformatics/btt703 (2014).

38 Oakley, H. et al. Intraneuronal β-Amyloid Aggregates, Neurodegeneration, and Neuron Loss in Transgenic Mice with Five Familial Alzheimer’s Disease Mutations: Potential Factors in Amyloid Plaque Formation. The Journal of Neuroscience 26, 10129–10140, doi:10.1523/jneurosci.1202-06.2006 (2006).

39 Richard, B. C. et al. Gene Dosage Dependent Aggravation of the Neurological Phenotype in the 5XFAD Mouse Model of Alzheimer’s Disease. J Alzheimers Dis 45, 1223–1236, doi:10.3233/jad-143120 (2015).

40 Doyle, J. P. et al. Application of a Translational Profiling Approach for the Comparative Analysis of CNS Cell Types. Cell 135, 749–762, doi:10.1016/j.cell.2008.10.029 (2008).

41 Heiman, M. et al. A Translational Profiling Approach for the Molecular Characterization of CNS Cell Types. Cell 135, 738–748, doi:10.1016/j.cell.2008.10.028 (2008).

42 Heiman, M., Kulicke, R., Fenster, R. J., Greengard, P. & Heintz, N. Cell type–specific mRNA purification by translating ribosome affinity purification (TRAP). Nat. Protocols 9, 1282–1291, doi:10.1038/nprot.2014.085 (2014).

43 Ayata, P. et al. Epigenetic regulation of brain region-specific microglia clearance activity. Nature Neuroscience 21, 1049–1060, doi:10.1038/s41593-018-0192-3 (2018).

44 Yoshiyama, Y. et al. Synapse loss and microglial activation precede tangles in a P301S tauopathy mouse model. Neuron 53, 337–351, doi:10.1016/j.neuron.2007.01.010 (2007).

45 Parkhurst, Christopher N. et al. Microglia Promote Learning-Dependent Synapse Formation through Brain-Derived Neurotrophic Factor. Cell 155, 1596–1609, doi:10.1016/j.cell.2013.11.030 (2013).

46 Zhao, X. F. et al. Targeting Microglia Using Cx3cr1-Cre Lines: Revisiting the Specificity. eNeuro 6, doi:10.1523/eneuro.0114-19.2019 (2019).

47 Faust, T. E. et al. A comparative analysis of microglial inducible Cre lines. bioRxiv, 2023.2001.2009.523268, doi:10.1101/2023.01.09.523268 (2023).

48 Ayata, P. Decoding 5HMC as an Active Chromatin Mark in the Brain and its Link to Rett Syndrome PhD thesis, The Rockefeller University, (2013).

49 Shrestha, P. et al. Cell-type-specific drug-inducible protein synthesis inhibition demonstrates that memory consolidation requires rapid neuronal translation. Nature Neuroscience 23, 281–292, doi:10.1038/s41593-019-0568-z (2020).

50 Dar, A. C., Dever, T. E. & Sicheri, F. Higher-Order Substrate Recognition of eIF2&#x3b1; by the RNA-Dependent Protein Kinase PKR. Cell 122, 887–900, doi:10.1016/j.cell.2005.06.044 (2005).

51 Back, S. H. et al. Translation Attenuation through eIF2α Phosphorylation Prevents Oxidative Stress and Maintains the Differentiated State in β Cells. Cell Metabolism 10, 13–26, 10.1016/j.cmet.2009.06.002 (2009).

52 Sharma, V. et al. eIF2α controls memory consolidation via excitatory and somatostatin neurons. Nature 586, 412–416, doi:10.1038/s41586-020-2805-8 (2020).

53 Jiang, S., Xia, R., Jiang, Y., Wang, L. & Gao, F. Vascular endothelial growth factors enhance the permeability of the mouse blood-brain barrier. PLoS One 9, e86407–e86407, doi:10.1371/journal.pone.0086407 (2014).

54 Lundy, D. J. et al. Inducing a Transient Increase in Blood–Brain Barrier Permeability for Improved Liposomal Drug Therapy of Glioblastoma Multiforme. ACS Nano 13, 97–113, doi:10.1021/acsnano.8b03785 (2019).

55 Gentile, I. et al. Asunaprevir, a protease inhibitor for the treatment of hepatitis C infection. Ther Clin Risk Manag 10, 493–504, doi:10.2147/tcrm.S66731 (2014).

56 Eley, T., Garimella, T., Li, W. & Bertz, R. J. Asunaprevir: A Review of Preclinical and Clinical Pharmacokinetics and Drug-Drug Interactions. Clin Pharmacokinet 54, 1205–1222, doi:10.1007/s40262-015-0299-6 (2015).

57 Hou, Y. et al. Ageing as a risk factor for neurodegenerative disease. Nature Reviews Neurology 15, 565–581, doi:10.1038/s41582-019-0244-7 (2019).

58 Leak, R. K. Adaptation and sensitization to proteotoxic stress. Dose Response 12, 24–56, doi:10.2203/dose-response.13-016.Leak (2013).

59 Horowitz, M. Epigenetics and cytoprotection with heat acclimation. J Appl Physiol (1985) 120, 702–710, doi:10.1152/japplphysiol.00552.2015 (2016).

60 Li, C. & Casanueva, O. Epigenetic inheritance of proteostasis and ageing. Essays in biochemistry 60, 191–202, doi:10.1042/EBC20160025 (2016).

61 Zhao, E. et al. KDM4C and ATF4 Cooperate in Transcriptional Control of Amino Acid Metabolism. Cell Reports 14, 506–519, doi:10.1016/j.celrep.2015.12.053 (2016).

62 Salvadores, N., Gerónimo-Olvera, C. & Court, F. A. Axonal Degeneration in AD: The Contribution of Aβ and Tau. Front Aging Neurosci 12, 581767, doi:10.3389/fnagi.2020.581767 (2020).

63 Condello, C., Yuan, P., Schain, A. & Grutzendler, J. Microglia constitute a barrier that prevents neurotoxic protofibrillar Aβ42 hotspots around plaques. Nature communications 6, 6176–6176, doi:10.1038/ncomms7176 (2015).

64 Spangenberg, E. E. et al. Eliminating microglia in Alzheimer’s mice prevents neuronal loss without modulating amyloid-β pathology. Brain : a journal of neurology 139, 1265–1281, doi:10.1093/brain/aww016 (2016).

65 Hopp, S. C. et al. The role of microglia in processing and spreading of bioactive tau seeds in Alzheimer’s disease. J Neuroinflammation 15, 269, doi:10.1186/s12974-018-1309-z (2018).

66 Litvinchuk, A. et al. Amelioration of Tau and ApoE4-linked glial lipid accumulation and neurodegeneration with an LXR agonist. Neuron, doi:10.1016/j.neuron.2023.10.023 (2023).

67 Chen, Y. et al. APOE3ch alters microglial response and suppresses Aβ-induced tau seeding and spread. Cell 187, 428–445.e420, doi:10.1016/j.cell.2023.11.029 (2024).

68 Kashani, A. et al. Loss of VGLUT1 and VGLUT2 in the prefrontal cortex is correlated with cognitive decline in Alzheimer disease. Neurobiol Aging 29, 1619–1630, doi:10.1016/j.neurobiolaging.2007.04.010 (2008).

69 Buskila, Y., Crowe, S. E. & Ellis-Davies, G. C. R. Synaptic deficits in layer 5 neurons precede overt structural decay in 5xFAD mice. Neuroscience 254, 152–159, doi:10.1016/j.neuroscience.2013.09.016 (2013).

70 Klegeris, A. et al. Alpha-synuclein activates stress signaling protein kinases in THP-1 cells and microglia. Neurobiol Aging 29, 739–752, doi:10.1016/j.neurobiolaging.2006.11.013 (2008).

71 Bernath, A. K., Murray, T. E., Shirley Yang, S., Gibon, J. & Klegeris, A. Microglia secrete distinct sets of neurotoxins in a stimulus-dependent manner. Brain Res 1807, 148315, doi:10.1016/j.brainres.2023.148315 (2023).

72 Mahadevan, N. R. et al. Cell-extrinsic effects of tumor ER stress imprint myeloid dendritic cells and impair CD8⁺ T cell priming. PLoS One 7, e51845, doi:10.1371/journal.pone.0051845 (2012).

73 McNally, B. D. et al. Long-chain ceramides are cell non-autonomous signals linking lipotoxicity to endoplasmic reticulum stress in skeletal muscle. Nat Commun 13, 1748, doi:10.1038/s41467-022-29363-9 (2022).

74 Badimon, A. et al. Negative feedback control of neuronal activity by microglia. Nature 586, 417–423, doi:10.1038/s41586-020-2777-8 (2020).

75 Zhou, Y. et al. Human and mouse single-nucleus transcriptomics reveal TREM2-dependent and TREM2-independent cellular responses in Alzheimer’s disease. Nature Medicine 26, 131–142, doi:10.1038/s41591-019-0695-9 (2020).

76 Mathys, H. et al. Single-cell atlas reveals correlates of high cognitive function, dementia, and resilience to Alzheimer’s disease pathology. Cell 186, 4365–4385.e4327, doi:10.1016/j.cell.2023.08.039 (2023).

77 Moncan, M. et al. Regulation of lipid metabolism by the unfolded protein response. J Cell Mol Med 25, 1359–1370, doi:10.1111/jcmm.16255 (2021).

78 Ates, G., Goldberg, J., Currais, A. & Maher, P. CMS121, a fatty acid synthase inhibitor, protects against excess lipid peroxidation and inflammation and alleviates cognitive loss in a transgenic mouse model of Alzheimer’s disease. Redox Biol 36, 101648, doi:10.1016/j.redox.2020.101648 (2020).

79 Kuhajda, F. P. et al. Synthesis and antitumor activity of an inhibitor of fatty acid synthase. Proc Natl Acad Sci U S A 97, 3450–3454, doi:10.1073/pnas.97.7.3450 (2000).

80 Castro, A. R., Morrill, W. E. & Pope, V. Lipid removal from human serum samples. Clin Diagn Lab Immunol 7, 197–199, doi:10.1128/cdli.7.2.197-199.2000 (2000).

81 Nguyen, D. C. et al. Extracellular vesicles from bone marrow-derived mesenchymal stromal cells support ex vivo survival of human antibody secreting cells. J Extracell Vesicles 7, 1463778, doi:10.1080/20013078.2018.1463778 (2018).

82 Chen, R. R. et al. Targeting of lipid metabolism with a metabolic inhibitor cocktail eradicates peritoneal metastases in ovarian cancer cells. Commun Biol 2, 281, doi:10.1038/s42003-019-0508-1 (2019).

83 Loftus, T. M. et al. Reduced food intake and body weight in mice treated with fatty acid synthase inhibitors. Science 288, 2379–2381, doi:10.1126/science.288.5475.2379 (2000).

84 Lourenco, M. V. et al. TNF-α mediates PKR-dependent memory impairment and brain IRS-1 inhibition induced by Alzheimer’s β-amyloid oligomers in mice and monkeys. Cell Metab 18, 831–843, doi:10.1016/j.cmet.2013.11.002 (2013).

85 Hoozemans, J. J. M. et al. Activation of the unfolded protein response in Parkinson’s disease. Biochemical and Biophysical Research Communications 354, 707–711, 10.1016/jbbrc.2007.01.043 (2007).

86 Ilieva, E. V. et al. Oxidative and endoplasmic reticulum stress interplay in sporadic amyotrophic lateral sclerosis. Brain 130, 3111–3123, doi:10.1093/brain/awm190 (2007).

87 Perry, D. C. et al. Progranulin Mutations as Risk Factors for Alzheimer Disease. JAMA Neurology 70, 774–778, doi:10.1001/2013.jamaneurol.393 (2013).

88 Ulland, T. K. et al. TREM2 Maintains Microglial Metabolic Fitness in Alzheimer’s Disease. Cell 170, 649–663.e613, 10.1016/j.cell.2017.07.023 (2017).

89 Marschallinger, J. et al. Lipid-droplet-accumulating microglia represent a dysfunctional and proinflammatory state in the aging brain. Nature Neuroscience 23, 194–208, doi:10.1038/s41593-019-0566-1 (2020).

90 Pimenova, A. A. et al. Alzheimer’s-associated PU.1 expression levels regulate microglial inflammatory response. Neurobiology of Disease 148, 105217, 10.1016/jnbd.2020.105217 (2021).

91 Segev, Y., Michaelson, D. M. & Rosenblum, K. ApoE ε4 is associated with eIF2α phosphorylation and impaired learning in young mice. Neurobiol Aging 34, 863–872, doi:10.1016/j.neurobiolaging.2012.06.020 (2013).

92 Kang, E.-B. et al. Treadmill exercise represses neuronal cell death and inflammation during Aβ-induced ER stress by regulating unfolded protein response in aged presenilin 2 mutant mice. Apoptosis 18, 1332–1347, doi:10.1007/s10495-013-0884-9 (2013).

93 Lee, S. et al. APOE modulates microglial immunometabolism in response to age, amyloid pathology, and inflammatory challenge. Cell Rep 42, 112196, doi:10.1016/j.celrep.2023.112196 (2023).

94 Tcw, J. et al. Cholesterol and matrisome pathways dysregulated in astrocytes and microglia. Cell 185, 2213–2233.e2225, doi:10.1016/j.cell.2022.05.017 (2022).

95 Shi, Y. et al. ApoE4 markedly exacerbates tau-mediated neurodegeneration in a mouse model of tauopathy. Nature 549, 523–527, doi:10.1038/nature24016 (2017).

96 Yin, Z. et al. APOE4 impairs the microglial response in Alzheimer’s disease by inducing TGFβ-mediated checkpoints. Nature Immunology 24, 1839–1853, doi:10.1038/s41590-023-01627-6 (2023).

97 Ma, T. et al. Suppression of eIF2α kinases alleviates Alzheimer’s disease–related plasticity and memory deficits. Nature Neuroscience 16, 1299–1305, doi:10.1038/nn.3486 (2013).

98 Ohno, M. Roles of eIF2α kinases in the pathogenesis of Alzheimer’s disease. Frontiers in Molecular Neuroscience 7, doi:10.3389/fnmol.2014.00022 (2014).

99 Carret-Rebillat, A.-S. et al. Neuroinflammation and Aβ Accumulation Linked To Systemic Inflammation Are Decreased By Genetic PKR Down-Regulation. Scientific Reports 5, 8489, doi:10.1038/srep08489 (2015).

100 Sidrauski, C., McGeachy, A. M., Ingolia, N. T. & Walter, P. The small molecule ISRIB reverses the effects of eIF2α phosphorylation on translation and stress granule assembly. Elife 4, e05033, doi:10.7554/eLife.05033 (2015).

101 Tible, M. et al. PKR knockout in the 5xFAD model of Alzheimer’s disease reveals beneficial effects on spatial memory and brain lesions. Aging Cell 18, e12887, doi:10.1111/acel.12887 (2019).

102 Bugallo, R. et al. Fine tuning of the unfolded protein response by ISRIB improves neuronal survival in a model of amyotrophic lateral sclerosis. Cell Death & Disease 11, 397, doi:10.1038/s41419-020-2601-2 (2020).

103 Halliday, M. et al. Partial restoration of protein synthesis rates by the small molecule ISRIB prevents neurodegeneration without pancreatic toxicity. Cell Death & Disease 6, e1672–e1672, doi:10.1038/cddis.2015.49 (2015).

104 Briggs, D. I. et al. Role of Endoplasmic Reticulum Stress in Learning and Memory Impairment and Alzheimer’s Disease-Like Neuropathology in the PS19 and APP^Swe^ Mouse Models of Tauopathy and Amyloidosis. eneuro 4, ENEURO.0025-0017.2017, doi:10.1523/eneuro.0025-17.2017 (2017).

105 Back, S. H. et al. Translation attenuation through eIF2alpha phosphorylation prevents oxidative stress and maintains the differentiated state in beta cells. Cell Metab 10, 13–26, doi:10.1016/j.cmet.2009.06.002 (2009).

106 Smith, H. L. et al. Astrocyte Unfolded Protein Response Induces a Specific Reactivity State that Causes Non-Cell-Autonomous Neuronal Degeneration. Neuron 105, 855–866.e855, doi:10.1016/j.neuron.2019.12.014 (2020).

107 Chung, K. M. et al. A systemic cell stress signal confers neuronal resilience toward oxidative stress in a Hedgehog-dependent manner. Cell Rep 41, 111488, doi:10.1016/j.celrep.2022.111488 (2022).

108 Chattopadhyay, A. et al. Cholesterol-Induced Phenotypic Modulation of Smooth Muscle Cells to Macrophage/Fibroblast–like Cells Is Driven by an Unfolded Protein Response. Arteriosclerosis, Thrombosis, and Vascular Biology 41, 302–316, doi:10.1161/ATVBAHA.120.315164 (2021).

109 Liddelow, S. A. et al. Neurotoxic reactive astrocytes are induced by activated microglia. Nature 541, 481–487, doi:10.1038/nature21029 (2017).

110 Huynh, T.-P. V. et al. Lack of hepatic apoE does not influence early Aβ deposition: observations from a new APOE knock-in model. Molecular Neurodegeneration 14, 37, doi:10.1186/s13024-019-0337-1 (2019).

111 Ghadami, S. & Dellinger, K. The lipid composition of extracellular vesicles: applications in diagnostics and therapeutic delivery. Front Mol Biosci 10, 1198044, doi:10.3389/fmolb.2023.1198044 (2023).

112 Oyadomari, S., Harding, H. P., Zhang, Y., Oyadomari, M. & Ron, D. Dephosphorylation of translation initiation factor 2alpha enhances glucose tolerance and attenuates hepatosteatosis in mice. Cell Metab 7, 520–532, doi:10.1016/j.cmet.2008.04.011 (2008).

113 Wang, C. et al. ATF4 regulates lipid metabolism and thermogenesis. Cell Research 20, 174–184, doi:10.1038/cr.2010.4 (2010).

114 Han, J. & Kaufman, R. J. The role of ER stress in lipid metabolism and lipotoxicity. J Lipid Res 57, 1329–1338, doi:10.1194/jlr.R067595 (2016).

115 Hannun, Y. A. & Luberto, C. Ceramide in the eukaryotic stress response. Trends in Cell Biology 10, 73–80, 10.1016/S0962-8924(99)01694-3 (2000).

116 Sozen, E. & Ozer, N. K. Impact of high cholesterol and endoplasmic reticulum stress on metabolic diseases: An updated mini-review. Redox Biology 12, 456–461, 10.1016/jredox.2017.02.025 (2017).

117 Jazvinšćak Jembrek, M., Hof, P. R. & Šimić, G. Ceramides in Alzheimer’s Disease: Key Mediators of Neuronal Apoptosis Induced by Oxidative Stress and Aβ Accumulation. Oxid Med Cell Longev 2015, 346783, doi:10.1155/2015/346783 (2015).

118 Snowden, S. G. et al. Association between fatty acid metabolism in the brain and Alzheimer disease neuropathology and cognitive performance: A nontargeted metabolomic study. PLoS Med 14, e1002266–e1002266, doi:10.1371/journal.pmed.1002266 (2017).

119 Huo, Z. et al. Brain and blood metabolome for Alzheimer’s dementia: findings from a targeted metabolomics analysis. Neurobiol Aging 86, 123–133, doi:10.1016/j.neurobiolaging.2019.10.014 (2020).

120 Varma, V. R. et al. Abnormal brain cholesterol homeostasis in Alzheimer’s disease—a targeted metabolomic and transcriptomic study. npj Aging and Mechanisms of Disease 7, 11, doi:10.1038/s41514-021-00064-9 (2021).

121 Romero-Molina, C., Garretti, F., Andrews, S. J., Marcora, E. & Goate, A. M. Microglial efferocytosis: Diving into the Alzheimer&#x2019;s disease gene pool. Neuron 110, 3513–3533, doi:10.1016/j.neuron.2022.10.015 (2022).

122 Wood, W. G., Li, L., Müller, W. E. & Eckert, G. P. Cholesterol as a causative factor in Alzheimer’s disease: a debatable hypothesis. J Neurochem 129, 559–572, doi:10.1111/jnc.12637 (2014).

123 Gratuze, M. et al. Activated microglia mitigate Aβ-associated tau seeding and spreading. J Exp Med 218, doi:10.1084/jem.20210542 (2021).

124 Lee, S.-H. et al. Trem2 restrains the enhancement of tau accumulation and neurodegeneration by &#x3b2;-amyloid pathology. Neuron 109, 1283–1301.e1286, doi:10.1016/j.neuron.2021.02.010 (2021).

125 Wang, C. et al. Microglial NF-κB drives tau spreading and toxicity in a mouse model of tauopathy. Nature Communications 13, 1969, doi:10.1038/s41467-022-29552-6 (2022).

126 Wendeln, A.-C. et al. Innate immune memory in the brain shapes neurological disease hallmarks. Nature 556, 332–338, doi:10.1038/s41586-018-0023-4 (2018).

127 Salani, F., Sterbini, V., Sacchinelli, E., Garramone, M. & Bossù, P. Is Innate Memory a Double-Edge Sword in Alzheimer’s Disease? A Reappraisal of New Concepts and Old Data. Front Immunol 10, 1768, doi:10.3389/fimmu.2019.01768 (2019).

128 Zhang, X. et al. Epigenetic regulation of innate immune memory in microglia. Journal of Neuroinflammation 19, 111, doi:10.1186/s12974-022-02463-5 (2022).

129 Höhn, A., Tramutola, A. & Cascella, R. Proteostasis Failure in Neurodegenerative Diseases: Focus on Oxidative Stress. Oxid Med Cell Longev 2020, 5497046–5497046, doi:10.1155/2020/5497046 (2020).

130 Sonninen, T. M., Goldsteins, G., Laham-Karam, N., Koistinaho, J. & Lehtonen, Š. Proteostasis Disturbances and Inflammation in Neurodegenerative Diseases. Cells 9, doi:10.3390/cells9102183 (2020).

131 Xu, Z.-X. et al. Elevated protein synthesis in microglia causes autism-like synaptic and behavioral aberrations. Nature communications 11, 1797–1797, doi:10.1038/s41467-020-15530-3 (2020).

132 Kalish, B. T. et al. Maternal immune activation in mice disrupts proteostasis in the fetal brain. Nat Neurosci 24, 204–213, doi:10.1038/s41593-020-00762-9 (2021).

133 Donat, C. K., Scott, G., Gentleman, S. M. & Sastre, M. Microglial Activation in Traumatic Brain Injury. Front Aging Neurosci 9, 208, doi:10.3389/fnagi.2017.00208 (2017).

134 Schaefer, A. et al. Control of Cognition and Adaptive Behavior by the GLP/G9a Epigenetic Suppressor Complex. Neuron 64, 678–691, doi:10.1016/j.neuron.2009.11.019 (2009).

135 Werneburg, S. et al. Targeted Complement Inhibition at Synapses Prevents Microglial Synaptic Engulfment and Synapse Loss in Demyelinating Disease. Immunity 52, 167–182.e167, doi:10.1016/j.immuni.2019.12.004 (2020).

136 Paxinos, G. & Franklin, K. B. J. Paxinos and Franklin’s the mouse brain in stereotaxic coordinates. (Elsevier Science, 2013).

137 Nahirney, P. C. & Tremblay, M. E. Brain Ultrastructure: Putting the Pieces Together. Front Cell Dev Biol 9, 629503, doi:10.3389/fcell.2021.629503 (2021).

138 Raji, C. A., Lopez, O. L., Kuller, L. H., Carmichael, O. T. & Becker, J. T. Age, Alzheimer disease, and brain structure. Neurology 73, 1899–1905, doi:10.1212/WNL.0b013e3181c3f293 (2009).

139 Liu, J. et al. Impaired adult myelination in the prefrontal cortex of socially isolated mice. Nature neuroscience 15, 1621–1623, doi:10.1038/nn.3263 (2012).

140 Rivera, D. S. et al. Effects of long-lasting social isolation and re-socialization on cognitive performance and brain activity: a longitudinal study in Octodon degus. Scientific Reports 10, 18315, doi:10.1038/s41598-020-75026-4 (2020).

141 Kim, D., Paggi, J. M., Park, C., Bennett, C. & Salzberg, S. L. Graph-based genome alignment and genotyping with HISAT2 and HISAT-genotype. Nature Biotechnology 37, 907–915, doi:10.1038/s41587-019-0201-4 (2019).

142 Liao, Y., Smyth, G. K. & Shi, W. featureCounts: an efficient general purpose program for assigning sequence reads to genomic features. Bioinformatics 30, doi:10.1093/bioinformatics/btt656 (2014).

143 Love, M. I., Huber, W. & Anders, S. Moderated estimation of fold change and dispersion for RNA-seq data with DESeq2. Genome Biology 15, 550, doi:10.1186/s13059-014-0550-8 (2014).

144 Hao, Y. et al. Dictionary learning for integrative, multimodal and scalable single-cell analysis. Nat Biotechnol 42, 293–304, doi:10.1038/s41587-023-01767-y (2024).

145 Chen, E. Y. et al. Enrichr: interactive and collaborative HTML5 gene list enrichment analysis tool. BMC Bioinformatics 14, 128, doi:10.1186/1471-2105-14-128 (2013).

146 Kuleshov, M. V. et al. Enrichr: a comprehensive gene set enrichment analysis web server 2016 update. Nucleic Acids Research 44, W90–W97, doi:10.1093/nar/gkw377 (2016).

147 Xie, Z. et al. Gene Set Knowledge Discovery with Enrichr. Current Protocols 1, e90, doi:10.1002/cpz1.90 (2021).

148 Rostami, J. et al. Crosstalk between astrocytes and microglia results in increased degradation of α-synuclein and amyloid-β aggregates. J Neuroinflammation 18, 124, doi:10.1186/s12974-021-02158-3 (2021).

149 Albuquerque, C., Joseph, D. J., Choudhury, P. & MacDermott, A. B. Dissection, plating, and maintenance of cortical astrocyte cultures. Cold Spring Harb Protoc 2009, pdb.prot5273, doi:10.1101/pdb.prot5273 (2009).

150 Hernandez-Ono, A. et al. Dynamic regulation of hepatic lipid metabolism by torsinA and its activators. JCI Insight 9, doi:10.1172/jci.insight.175328 (2024).

151 Bligh, E. G. & Dyer, W. J. A rapid method of total lipid extraction and purification. Can J Biochem Physiol 37, 911–917, doi:10.1139/o59-099 (1959).

152 Chen, S. et al. Lipidomic characterization of extracellular vesicles in human serum. J Circ Biomark 8, 1849454419879848, doi:10.1177/1849454419879848 (2019).

